# A novel pathway linking plasma membrane and chloroplasts is co-opted by pathogens to suppress salicylic acid-dependent defences

**DOI:** 10.1101/837955

**Authors:** Laura Medina-Puche, Huang Tan, Vivek Dogra, Mengshi Wu, Tabata Rosas-Diaz, Liping Wang, Xue Ding, Dan Zhang, Xing Fu, Chanhong Kim, Rosa Lozano-Duran

**Affiliations:** Shanghai Center for Plant Stress Biology, CAS Center for Excellence in Molecular Plant Sciences, Chinese Academy of Sciences, Shanghai 201602, China; University of the Chinese Academy of Sciences, Beijing 100049, China

**Keywords:** Plasma membrane, chloroplast, retrograde signalling, salicylic acid, defence, geminivirus, effector, pathogen, plant

## Abstract

Chloroplasts are crucial players in the activation of defensive hormonal responses during plant-pathogen interactions. Here, we show that a plant virus-encoded protein re-localizes from the plasma membrane to chloroplasts upon triggering plant defence, interfering with the chloroplast-dependent activation of anti-viral salicylic acid (SA) biosynthesis. Strikingly, we have found that plant pathogens from different kingdoms seem to have convergently evolved to target chloroplasts and impair SA-dependent defences following an association with membranes, which is based on the co-existence of two subcellular targeting signals, an N-myristoylation site and a chloroplast transit peptide. This pattern is also present in plant proteins, at least one of which conversely activates SA defences from the chloroplast. Taken together, our results suggest that a pathway linking plasma membrane to chloroplasts and activating defence exists in plants, and that such pathway has been co-opted by plant pathogens during host-pathogen co-evolution to promote virulence through suppression of SA responses.

## INTRODUCTION

Beyond their role as photosynthetic organelles enabling photoautotrophy, chloroplasts are emerging as hubs in the integration of environmental stimuli and determinants of downstream responses (Chan et al., 2016, de Souza et al., 2017, Zhu, 2016, de Torres Zabala et al., 2015, Xiao et al., 2013, Zhao et al., 2016). A growing body of evidence substantiates the fundamental function of chloroplasts in orchestrating defence responses: upon perception of a biotic threat, this organelle acts as the source of calcium and reactive oxygen species (ROS) bursts and communicates with the nucleus through retrograde signalling, initiating a signalling cascade that leads to the expression of defence-related genes, including those responsible for the biosynthesis of the defense hormone salicylic acid (SA), subsequently produced in the chloroplast stroma (Serrano et al., 2016; Nomura et al., 2012). The chloroplast-nucleus communication during Pattern-Triggered Immunity (PTI) involves the so-called Calcium Sensing Receptor (CAS), a thylakoid membrane-associated protein; although the exact molecular function of CAS in unclear, this protein is required for PTI-induced transcriptional reprograming, SA biosynthesis, callose deposition, and anti-bacterial resistance (Nomura et al., 2012). How the information of pathogen attack is relayed from the cell periphery (e.g. during a bacterial infection) or possibly other subcellular compartments (e.g. during viral infections) to chloroplasts, however, remains elusive, while the molecular basis of retrograde signalling during defence responses are largely unexplored.

In agreement with a prominent role of chloroplast function in defence and in the context of the arms race between pathogens and hosts, a number of virulence factors from evolutionarily unrelated pathogens belonging to different kingdoms of life, including bacteria, viruses, fungi, and oomycetes, have been described to target this organelle (e.g. de Torres Zabala et al., 2015, Fondong et al., 2007, Rodriguez-Herva et al., 2012, Rosas-Diaz et al., 2018, Jelenska et al., 2007, Jelenska et al., 2010, Li et al., 2014, Liu et al., 2018, Park et al., 2017, Petre et al., 2016). Some plant viruses co-opt chloroplast membranes to build viral replication factories (Jin et al., 2018; Bhattacharyya and Chakraborty, 2018) but, intriguingly, plant proteins from other viruses can also be found in chloroplasts (Vaira et al., 2018; Bhattacharyya and Chakraborty, 2018; Rosas-Diaz et al., 2018; Zhan et al., 2018; Bhattacharyya et al., 2015; Liu et al., 2015). In a few cases, it has been demonstrated or suggested that chloroplast-localized viral proteins can promote viral pathogenesis (Gnanasekaran et al., 2019; Bhattacharyya et al., 2015; Krenz et al., 2010). We recently showed that the C4 protein from the geminivirus *Tomato yellow leaf curl virus* (TYLCV), a DNA virus replicating in the nucleus, contains two overlapping localization signals, namely an N-myristoylation motif that tethers it to the plasma membrane (PM) and a chloroplast transit peptide (cTP) that targets it to the chloroplast (Rosas-Diaz et al., 2018). While at least one of the roles of C4 at the PM is to suppress the cell-to-cell movement of RNA interference (RNAi) (Rosas-Diaz et al., 2018; Fan et al., 2019), the function of C4 in the chloroplast is so far enigmatic.

Here, we show that C4 shifts its localization from the PM to chloroplasts upon activation of immune signalling by the replication-associated viral protein (Rep) or by exogenous treatments with the bacterial peptide elicitor flg22 or the endogenous peptide Pep1. Once inside the organelle, C4 associates with the thylakoid transmembrane protein CAS. The effect of C4 is consistent with a suppression of CAS function in retrograde signalling, as expression of this viral protein leads to decreased CAS-dependent immune responses, namely lower amplitude in cytosolic calcium burst upon pathogen perception, depressed transcriptional changes, reduced SA accumulation, and compromised resistance against the plant pathogenic bacterium *Pseudomonas syringae*. The C4-facilitated manipulation of chloroplast-mediated defences is biologically relevant, since knocking down *CAS* or depleting downstream SA promotes viral accumulation and partially complements a C4 null mutation in the virus, pointing at the suppression of SA responses as one of the main roles of this virus-encoded protein.

Strikingly, we have found the coexistence of the overlapping targeting signals contained in C4, the N-myristoylation site and the cTP, in a number of evolutionarily unrelated plant pathogen-encoded effector proteins, including those from DNA viruses, RNA viruses, and bacteria. Importantly, some of these effectors show dual membrane/chloroplast localization and suppress chloroplast-dependent defences when targeted to this organelle. Moreover, co-occurrence of these two targeting signals can also be found in a conserved set of plant proteins, many of which have a described role in the regulation of defence responses. We demonstrate that one of these proteins, Calcium Protein Kinase 16 (CPK16), re-localizes from the PM to chloroplasts upon flg22 treatment to promote chloroplast-dependent defences. Based on the results presented here, we propose that a protein relocalization-dependent pathway physically linking PM and chloroplasts and regulating defence exists in plants, and that this pathway has been co-opted by pathogens during evolution to suppress defence responses and promote virulence.

## RESULTS

### The C4 protein from the geminivirus *Tomato yellow leaf curl virus* shifts its localization from the plasma membrane to chloroplasts upon activation of defence

When expressed in plant cells, the C4 protein from TYLCV fused to GFP at its C-terminus localizes preferentially at the PM, with a minor fraction visible in chloroplasts (Figure 1A, B; Rosas-Diaz et al., 2018). However, when co-expressed with a TYLCV infectious clone, C4-GFP strongly accumulates in chloroplasts and is depleted from the PM (Figure 1C); co-expression with the viral Replication-associated protein (Rep) alone, but not other virus-encoded proteins, is sufficient to trigger this shift in localization (Figure 1D). Expression of Rep in *Nicotiana benthamiana* leads to the activation of defence responses, probably owing to the recognition of the protein or its activity by the plant (Ding et al., 2019); in order to test whether activation of defence could lead to the re-localization of C4 from the PM to chloroplasts, we treated *N. benthamiana* leaves transiently expressing C4-GFP with the bacterial peptide elicitor flg22, which is recognized by a receptor complex at the cell surface and activates pattern-triggered immunity (PTI) (Felix et al., 1999; Gomez-Gomez and Boller, 2000). As shown in Figure 1E and Supplemental figure 1A, flg22 treatment results in a clear accumulation of C4-GFP in chloroplasts, at the expense of the PM pool; treatment with the translation inhibitor cycloheximide (CHX) supports the idea that the shift in PM/chloroplast C4-GFP accumulation ratio is due to physical re-localization of the protein, and not to differential targeting following synthesis *de novo*. The endogenous immunogenic peptide Pep1 similarly triggers the chloroplast re-localization of C4 in *N. benthamiana*, while treatment with the bacterial derived peptide elf18, for which *N. benthamiana* lacks a receptor, has no noticeable effect of the subcellular distribution of the protein (Supplemental figure 1B-D). The increase in chloroplast-localized C4-GFP following flg22 treatment can also be detected by organelle purification and western blot (Supplemental figure 1E); additionally, a 33 KDa variant of C4, corresponding to C4-GFP after the chloroplast import-coupled cleavage of the cTP (mature chloroplast form), accumulates at the expense of the full-length version of the fusion protein (Supplemental figure 1F). Chloroplast fractionation indicates that C4 is a peripheral thylakoid membrane protein facing the stroma (Figure 1F, G; Supplemental figure 1G). Nevertheless, accumulation of C4 in the chloroplast in transgenic Arabidopsis lines expressing the non-myristoylable C4_G2A_ mutant version, which localizes to chloroplasts exclusively (Figure 1B; Rosas-Diaz et al., 2018), does not affect photosynthetic efficiency, chloroplast ultrastructure, or general plant development (Figure 1H-L).

**Figure 1.**
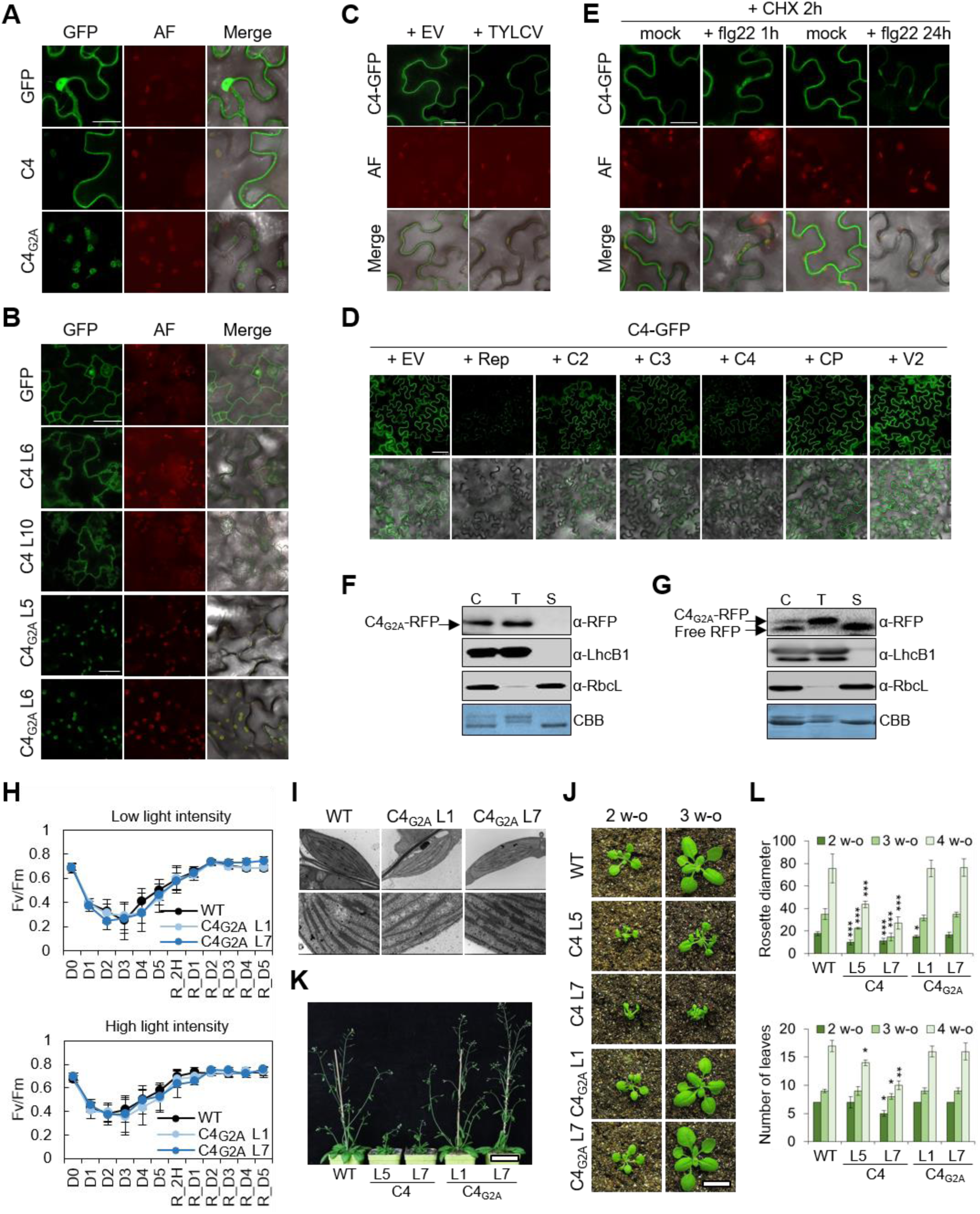
The C4 protein from the geminivirus TYLCV shifts its localization from plasma membrane to chloroplasts upon activation of defence. (**A**) C4 shows a plasma membrane (PM)/chloroplast dual localization upon transient expression in *N. benthamiana* leaves. Localization of the wild-type and the non-myristoylable (C4_G2A_) C4 versions fused to GFP at two days post-infiltration (dpi). Scale bar = 25 µm. AF: Autofluorescence. **(B**) C4 shows a PM/chloroplast dual localization in Arabidopsis transgenic lines. Localization of the wild-type and the non-myristoylable (C4_G2A_) C4 versions fused to GFP in cotyledons of 7-day-old seedlings. Scale bar = 25 µm. AF: Autofluorescence. (**C**) C4 re-localizes from PM to chloroplasts during TYLCV infection in *N. benthamiana* leaves. Localization of C4-GFP was compared when expressed alone or co-infiltrated with a TYLCV infectious clone at 2dpi. Scale bar = 25 µm. AF: Autofluorescence. (**D**) The virus-encoded Rep protein, but not other viral proteins, triggers chloroplast localization of C4-GFP. C4-GFP was co-infiltrated with *Agrobacterium* clones to express each TYLCV protein independently in *N. benthamiana* leaves and localization was determined at 2dpi. SMD settings: gating 1%, laser 10, gain 90%. With V2: gating 1%, laser 10, gain 30%. EV: empty vector. Scale bar = 50 µm. (**E**) C4 re-localizes from PM to chloroplasts in response to treatment with the bacterial peptide elicitor flg22 upon transient expression in *N. benthamiana* leaves. Localization of C4-GFP was compared upon treatment with 1 µM flg22 or mock treatment (1h and 24 h post-treatment) in the presence of CHX (50 µg/ml, 2 h). Scale bar = 25 µm. AF: Autofluorescence. (**F**) Transiently expressed C4_G2A_-RFP in *N. benthamiana* is associated to the chloroplast thylakoid membrane. Isolated chloroplasts were separated into membrane and stroma fractions upon transient expression of the non-myristoylable C4 version (C4_G2A_) fused to RFP in *N. benthamiana* leaves (2dpi). (C: total chloroplast; T: thylakoid; S: stroma; LhcB1: light harvesting complex protein B1 (25 kDa), a thylakoid membrane protein; RbcL: rubisco large subunit (52.7 kDa), a stromal protein). CBB: Comassie brilliant blue. (**G**) Stably expressed C4_G2A_-RFP in Arabidopsis is associated to the chloroplast thylakoid membrane. Isolated chloroplasts from three-week-old Arabidopsis transgenic lines expressing the non-myristoylable C4 version (C4_G2A_) fused to RFP were separated into membrane and stroma fractions. (C: total chloroplast; T: thylakoid; S: stroma; LhcB1: light harvesting complex protein B1 (25 kDa), a thylakoid membrane protein; RbcL: rubisco large subunit (52.7 kDa), a stromal protein). CBB: Comassie brilliant blue. (**H**) Chloroplast-localized C4 does not affect photosynthetic efficiency. Ten-day-old wild-type or 35S:C4_G2A_ (Lines 1 (L1) and 7 (L7)) seedlings were grown in ½ MS-sucrose agar medium under constant light (low light intensity: 40 µmol m^−2^ s^−1^; high light intensity: 80 µmol m^−2^ s^−1^) for 5 days at 22°C, followed by 5 days under constant light (very high intensity: 300 µmol m^−2^ s^−1^) in cold stress conditions (10°C), when Fv/Fm was recorded every day (D: day; D0 to D5: from day 0 to day 5); seedlings were then transferred to the initial conditions (recovery), and Fv/Fm was recorded at the indicated time points (R: recovery; R_2h: 2 h after recovery; R_D1 to R_D5: from day 1 to day 5 after recovery). (**I**) Chloroplast-localized C4 does not affect chloroplast ultrastructure. Ten-day-old wild-type or 35S:C4_G2A_ (Lines 1 (L1) and 7 (L7)) seedlings were grown in ½ MS -sucrose agar medium under long day conditions and used for transmission electron microscopy imaging. (**J**) Expression of chloroplast-localized C4 does not visibly affect development of 2- and 3-week-old transgenic Arabidopsis lines (Bar = 1 cm). (**K**) Expression of chloroplast-localized C4 does not visibly affect development of five week-old transgenic Arabidopsis lines (Bar = 7 cm). (**L**) Developmental phenotypes of transgenic Arabidopsis plants expressing C4 or C4_G2A_ grown in long day conditions. Rosette diameter is in mm. Bars represent SE of n = 3. Asterisks indicate a statistically significant difference (\**P* < 0.05, \*\**P* < 0.01, \*\*\**P* < 0.001) according to a one-way ANOVA with post-hoc Dunnett’s multiple comparisons test.

### C4 interacts with Calcium Sensing Receptor in chloroplasts and suppresses downstream immune responses

Following activation of PTI, retrograde signalling allows communication of chloroplasts with the nucleus, activating expression of genes required for the biosynthesis of the defense hormone salicylic acid (SA) and ultimately leading to the accumulation of this compound (Qi et al., 2018; Zhang and Li, 2019; Nomura et al., 2012). Since activation of PTI leads to the re-localization of C4 from the PM to the chloroplast, and with the aim to detect a potential interference of C4 with this signalling cascade, we measured the expression of SA biosynthetic and responsive genes (*ICS1* and *PR1*, respectively) as well as the accumulation of SA after treatment with flg22 in transgenic Arabidopsis plants expressing the chloroplast-localized C4_G2A_ (Figure 2A,B and Supplemental figure 2A). As shown in Figure 2A and B, the presence of C4 in the chloroplast results in lower expression of SA-related genes and halved SA content after flg22 treatment. C4, however, does not impair SA perception or downstream responses, as demonstrated by exogenous SA treatments (Supplemental figure 2B). Importantly, the suppression of SA biosynthesis seems to be relevant for the viral infection, in agreement with previous observations (Li et al., 2019), since transgenic Arabidopsis or tomato plants depleted in SA (*NahG* transgenic plants) support higher viral accumulation and can partially complement a null mutation in C4, suggesting that the suppression of SA responses is one of the main functions of C4 in the context of the infection (Figure 2C, Supplemental figure 2C).

**Figure 2.**
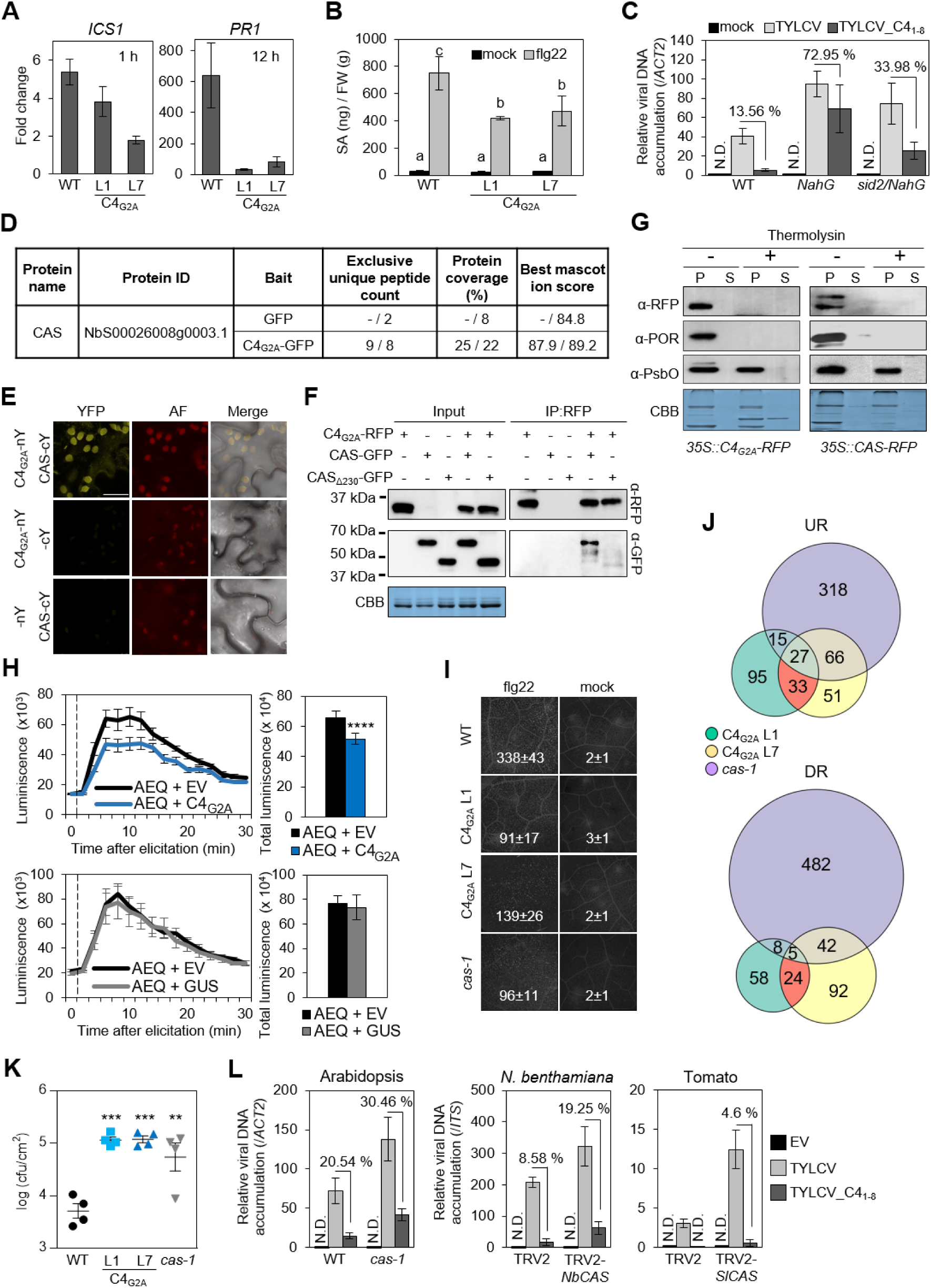
Chloroplast-localized C4 interacts with Calcium sensing receptor (CAS) and suppresses downstream immune responses. (**A**) Chloroplast-localized C4 suppresses retrograde signalling. Flg22-induced *ICS1* and *PR1* expression was analysed in ten-day-old wild-type or transgenic Arabidopsis lines expressing C4_G2A_ at the indicated time points by qRT–PCR. Seedlings were grown on ½ MS agar plates under long day conditions prior to transfer to liquid ½ MS for elicitation. *ACT2* was used as the normalizer. Data are mean ± SE of three independent biological replicates. (**B**) Chloroplast-localized C4 suppresses SA accumulation upon flg22 treatment. Total SA content was quantified in ten-day-old wild-type or transgenic Arabidopsis lines expressing C4_G2A_. Seedlings were grown on ½ MS agar plates under long day conditions prior to transfer to liquid ½ MS for elicitation with 1 µM flg22 during 12 h. Data are mean ± SE of three independent biological replicates. Lowercase letters indicate statistically significant differences between mean values (*P* <0.001), according to a one-way ANOVA with post-hoc Tukey’s multiple comparisons test. (**C**) Viral accumulation in Arabidopsis *NahG* and *sid2/NahG* plants infected with TYLCV wild type (TYCLV) or a mutant version carrying a premature stop codon in position 9 of the C4 gene (TYLCV_C4_1-8_). The relative viral DNA accumulation in plants was determined by qPCR of total DNA extracted from the six youngest rosette leaves at 21 days post-inoculation (dpi). Values represent the average of eight plants. Error bars represent SE. ND: not detectable. “%” indicates the percentage of TYLCV_C4_1-8_ accumulation compared to TYLCV (100%). Experiments were repeated at least three times with similar results; data from one biological replicate are shown. (**D**) CAS peptides identified in affinity purification-mass spectrometry (AP-MS) analysis after purification of C4_G2A_-GFP and GFP from *N. benthamiana* leaves in two independent experiments. “-” indicates no peptide was detected. (**E**) Biomolecular fluorescence complementation (BiFC) assay showing the interaction between C4_G2A_ and CAS. YFP fluorescence was observed in chloroplasts upon transient co-expression in *N. benthamiana* leaves at 2 dpi. Scale bar = 25 µm. AF: Autofluorescence. (**F**) C4_G2A_ interacts with the C-terminal part of CAS by co-immunoprecipitation upon transient co-expression in *N. benthamiana* leaves. Full-length or a truncated version of CAS lacking the rhodanese-like domain (CAS_∆230_) were used. Molecular weight is indicated. CBB: Comassie brilliant blue. (**G**) Topology analysis of chloroplast-localized C4 and CAS proteins. Thermolysin digestion of freshly isolated thylakoid membranes indicates that chloroplastic C4 is a stroma-facing thylakoid peripheral protein that interacts with the stroma-facing C-terminal part of CAS (P: thylakoid pellet; S: supernatant; POR: protochlorophyllide oxidoreductase (37 kDa), a thylakoid peripheral protein towards stroma; PsbO: photosystem II subunit O (33 kDa), a lumen localized protein). CBB: Comassie brilliant blue. (**H**) Chloroplast-localized C4 impairs the CAS-dependent cytoplasmic calcium transient in response to flg22. The calcium sensor aequorin (AEQ) was transiently co-expressed with C4_G2A_ or GUS (as negative control) in *N. benthamiana* leaves. The vertical dashed lines indicate the time at which treatment was initiated. AEQ luminescence was recorded during 30 min every two minutes (luminiscence in cp 120^−s^). Total luminescence was calculated at the end of the experiment. Values represent the average of six plants. Error bars represent SE. Asterisks indicate a statistically significant difference (\*\*\*\**P* < 0.0001) according to a two-tailed comparisons t-test. This experiment were repeated at least three times with similar results; data from one experiment are shown. EV: empty vector. (**I**) Chloroplast-localized C4 impairs flg22-induced callose deposition. Representative pictures and the average number (with SD) of callose deposits per 2.5 mm^2^ (n=10) are shown. This experiment was repeated three times with similar results; results from one experiment are shown. (**J**) Transcriptional overlap between transgenic Arabidopsis plants expressing C4_G2A_ and a *cas-1* mutant upon activation of plant immunity by treatment with the bacterial peptide elicitor flg22 (1 µM, 12 h). RNA-seq data were obtained from ten-day-old Arabidopsis seedlings grown on ½ MS agar plates under long-day conditions prior to transfer to ½ MS liquid for elicitation. UR: up-regulated; DR: down-regulated. Comparisons are made with treated control (wild-type) plants. (**K**) Transgenic Arabidopsis plants expressing C4_G2A_ display increased susceptibility to the bacterial pathogen *P. syringae* pv. *tomato* DC3000. Four-week-old short-day-grown plants were inoculated by infiltration with *Pto*DC3000. Three days later, bacteria were extracted from 7-mm leaf discs from three different leaves of four independent plants and incubated at 28 °C for two days. Data are mean ± SE of n = 4. This experiment was repeated 3 times with similar results; results from one experiment are shown. Asterisks indicate a statistically significant difference (\*\**P* < 0.01, \*\*\**P* < 0.001) according to a one-way ANOVA with post-hoc Dunnett’s multiple comparisons test. (**L**) Mutation in *CAS* favours infection by TYLCV and partially complements a C4 null mutant virus. Relative accumulation of TYLCV wild type (TYCLV) or a mutant version carrying a premature stop codon in position 9 of the C4 gene (TYLCV_C4_1-8_) in *cas-1* Arabidopsis plants and *N. benthamiana* and tomato plants in which *CAS* has been silenced by VIGS, as determined by qPCR. Total DNA was extracted from the 6 youngest apical leaves in Arabidopsis and from the 2 youngest apical leaves in *N. benthamiana* and tomato, at 21 dpi. Values represent the average of 6 plants. Error bars represent SE. ND: not detectable. EV: empty vector. Experiments were repeated at least three times with similar results; results from one experiment are shown.

We reasoned that the C4-mediated hampering of immune chloroplast retrograde signalling could be most likely based on the interaction with some plant protein, hence we performed affinity purification followed by mass spectrometry analysis (AP-MS) upon transient expression of C4_G2A_ in *N. benthamiana* to identify potential interactors of the chloroplast-localized C4. This approach unveiled the plant-specific Calcium Sensing Receptor (CAS) as a putative interactor of C4 in the chloroplast (Figure 2D); this protein-protein interaction was subsequently confirmed by bimolecular fluorescent complementation (BiFC) and co-immunoprecipitation (co-IP) analyses (Figure 2E, F). CAS is described as a thylakoid membrane-spanning protein (Nomura et al., 2012; Cutolo et al., 2019); topology analyses demonstrate that the C-terminus of CAS faces the stroma and interacts with C4, itself associated to the thylakoid membrane as peripheral protein (Figure 2F,G and Supplemental figure 2D-F). Interestingly, CAS has been previously demonstrated to be required for retrograde signalling in PTI and the ensuing activation of SA biosynthesis (Nomura et al., 2012). Therefore, we next tested known CAS-dependent responses to flg22 in our transgenic lines expressing chloroplast-localized C4. As shown in Figure 2H-K, plants expressing C4_G2A_ display immune defects at different levels, including reduced cytoplasmic calcium burst (Figure 2H and Supplemental figure 2G), lower callose deposition (Figure 2I), defective transcriptional reprogramming (Figure 2J and Supplemental figure 2H), and increased susceptibility to *P. syringae* pv. *tomato* DC3000 (Figure 2K and Supplemental figure 2I), all of them phenocopying a *cas* mutant and hence consistent with an inhibition of this protein. Of note, and as expected, flg22 perception is not affected in C4_G2A_ transgenic lines or in a *cas* mutant, since the early apoplastic burst of reactive oxygen species (ROS) as well as the late seedling growth inhibition occur normally in these plants in response to the peptide treatment (Supplemental figure 2J, K; Nomura et al., 2012). The biological relevance of the potential C4-mediated inhibition of CAS is illustrated by the fact that a *cas* Arabidopsis mutant as well as *N. benthamiana* or tomato plants in which *CAS* has been knocked down by virus-induced gene silencing (VIGS) are all more susceptible to TYLCV infection, indicating that suppression of CAS function has a positive impact on the virus’ performance (Figure 2L and Supplemental figure 2L).

### Proteins encoded by evolutionarily unrelated pathogens contain overlapping membrane- and chloroplast-targeting signals and suppress chloroplast-mediated defences

Our results show that the geminivirus TYLCV has evolved a strategy, through the action of the virus-encoded C4 protein and its shuttling from the PM to the chloroplast, to interfere with chloroplast-mediated SA-dependent defence responses and promote viral performance. Since SA has been proven to counter virus infection in different plant-virus interactions (e.g. Chen et al., 2010; Ji et al., 2001; Kachroo et al., 2000; Malamy et al., 1990) and chloroplasts are crucial players in this pathway, we hypothesized that other plant viruses might have independently evolved similar strategies to target this organelle when they are perceived by the plant. In an attempt to identify such hypothetical proteins, we searched public databases for proteins encoded by plant viruses containing both an N-myristoylation motif and a cTP. Surprisingly, we found ~400 viral proteins in which these two localization signals co-exist, encoded by viral species belonging to four different families (Figure 3A); most of these proteins are encoded by geminiviruses (Supplemental figure 3A). From this set of proteins, we selected four to test the functionality of their localization signals as well as their biological effect following chloroplast localization: C4 from the curtovirus *Beet curly top virus* (BCTV) (Fam. *Geminiviridae*); AC4 from the bipartite begomovirus *East African cassava mosaic virus* (EACMV) (Fam. *Geminiviridae*); the capsid protein (CP) from the cucumovirus *Cucumber mosaic virus* (CMV) (Fam. *Bromoviridae*); and the P3 protein from the nepovirus *Grapevine fanleaf virus* (GFLV) (Fam. *Secoviridae*) (Supplemental figure 3B). In all cases, the wild-type version of these viral proteins fused to GFP transiently expressed in *N. benthamiana* appeared as associated to membranes, while the non-myristoylable (G2A) mutant versions accumulated in the chloroplast (Figure 3B). Treatment with flg22 triggered the re-localization of C4 from BCTV and AC4 from EACMV to chloroplasts; this effect could not be clearly observed for CP and P3, suggesting that a different signal might be required in these two cases (Figure 3C). We next tested immune readouts in *N. benthamiana* leaves transiently expressing the chloroplast-localized (G2A) version of these proteins. Strikingly, the chloroplastic versions of all four viral proteins reduced the expression of *NbICS1*, *NbPR1*, and other defence-related genes, as well as the cytosolic calcium burst, in response to flg22, while all but P3 enhanced susceptibility against the pathogenic bacteria *P. syringae* pv tomato DC3000 *ΔhopQ1* (Figure D-F, Supplemental figure 3C-E), indicating that these phylogenetically unrelated viral effectors are capable of interfering with SA-mediated defences from the chloroplast. Expression of these proteins does not interfere with the apoplastic ROS burst that follows flg22 treatment, indicating that perception of the PAMP occurs normally (Figure 3G, Supplemental figure 3F).

**Figure 3.**
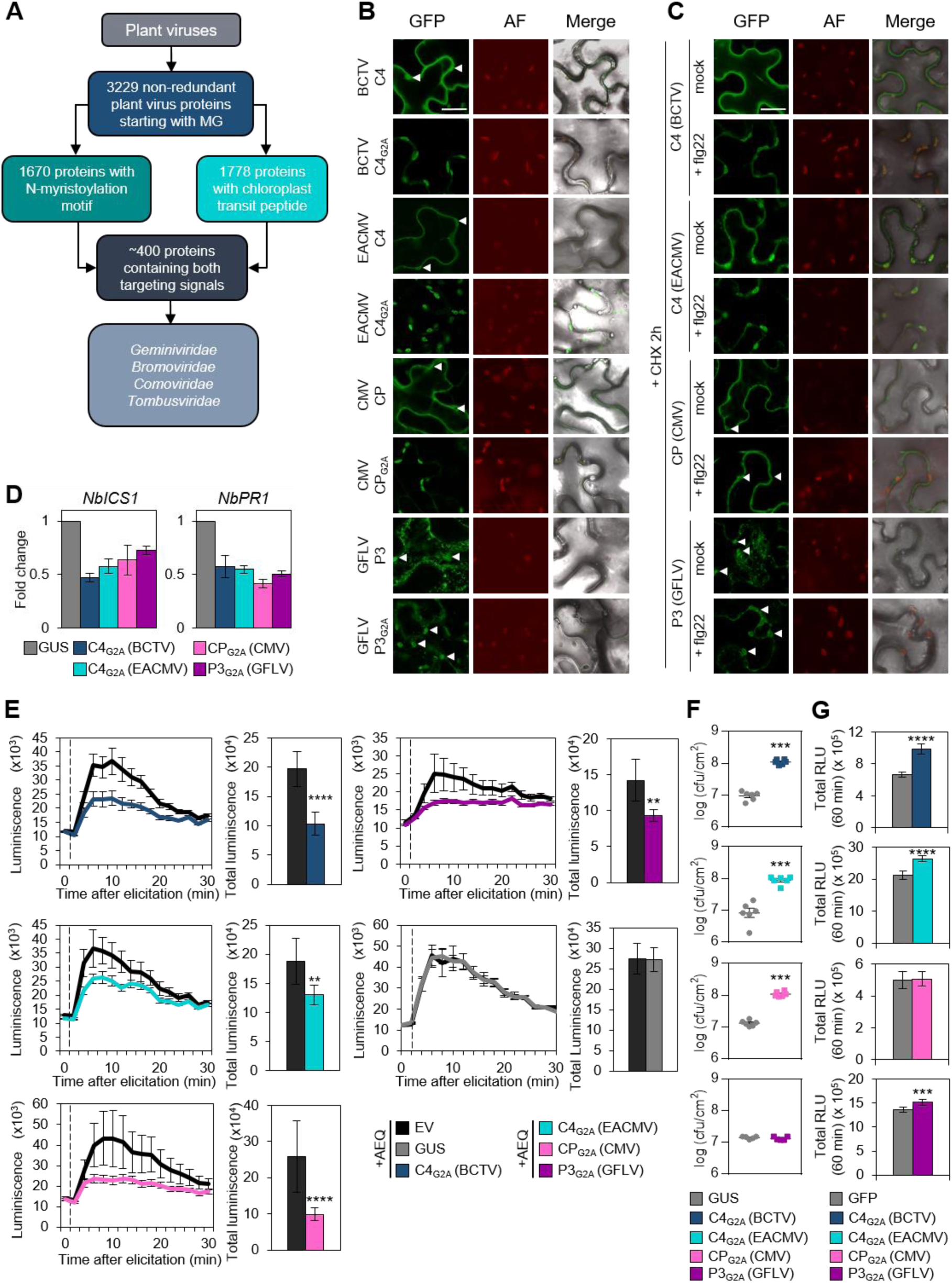
Proteins encoded by plant viruses contain overlapping membrane- and chloroplast-targeting signals and suppress chloroplast-mediated defences. (**A**) Flow diagram illustrating the number of proteins encoded by plant viruses and containing an N-myristoylation motif, a chloroplast transit peptide, or both, and the families they belong to. (**B**) Selected viral proteins with overlapping N-myristoylation motif and cTP (C4 from the curtovirus *Beet curly top virus* (BCTV; Fam. *Geminiviridae*); AC4 from the bipartite begomovirus *East African Cassava Mosaic Virus* (EACMV; Fam. *Geminiviridae*); the capsid protein (CP) from the cucumovirus *Cucumber mosaic virus* (CMV; Fam. *Bromoviridae*); and the P3 protein from the nepovirus *Grapevine fanleaf virus* (GFLV; Fam. *Secoviridae*) show PM/chloroplast dual localization upon transient expression in *N. benthamiana* leaves. Localization of the wild-type proteins and the non-myristoylable (G2A) versions fused to GFP was observed at two days post-infiltration (dpi). Scale bar = 25 µm. AF: Autofluorescence. This experiment was repeated 3 times with similar results. Arrowheads indicate chloroplasts. (**C**) Geminiviral proteins with overlapping N-myristoylation motif and cTP (C4 from BCTV and AC4 from EACMV) re-localize to chloroplasts upon activation of immunity. Localization of the wild-type proteins was monitored under different conditions (mock vs. 1 µM flg22, 24 h post-treatment). Scale bar = 25 µm. AF: Autofluorescence. This experiment was repeated 3 times with similar results. Arrowheads indicate chloroplasts. (**D**) Viral proteins with the overlapping N-myristoylation motif and cTP suppress expression of SA-related genes upon activation of plant immunity in *N. benthamiana* leaves. PAMP treatment (1 µM flg22) was performed 48 h after transient transformation. Leaf discs from three plants were collected separately 9 hours after treatment. Gene expression was analysed by qRT–PCR. *NbEF1α* was used as an internal standard. Values represent the average of 4 independent experiments with 3 plants used in each replicate. Data are mean ± SE of three independent experiments. (**E**) Chloroplast-localized viral proteins impair the CAS-dependent cytoplasmic calcium transient in response to flg22. The calcium sensor aequorin (AEQ) was coexpressed with the untagged non-myristoylable (G2A) versions of the viral proteins or GUS (as a negative control) in *N. benthamiana* leaves. The vertical dashed lines indicate the time at which flg22 treatment was initiated. AEQ luminescence was recorded during 30 min every two minutes (luminiscence in cp 120^−s^). Total luminescence was calculated at the end of the experiment. Values represent the average of 6 plants. Error bars represent SE. Asterisks indicate a statistically significant difference (\*\**P* < 0.01, \*\*\*\**P* < 0.0001) according to a two-tailed comparisons t-test. Experiments were repeated at least three times with similar results; results from one experiment are shown. EV: empty vector. (**F**) Chloroplast-localized viral proteins increase susceptibility to the plant pathogenic bacterial strain *P. syringae* pv. *tomato* DC3000 *∆hopQ1-1*. Leaves of four-week-old *N. benthamiana* plants were inoculated by infiltration with a *Pto*DC3000*∆hopQ1-1* suspension 24 h after transient transformation. Three days later, bacteria were extracted from 7-mm leaf discs from two different leaves of 3 independent plants and incubated at 28 °C for 2 days. Data are mean ± SE. Asterisks indicate a statistically significant difference (\*\*\**P* < 0.001) according to a two-tailed comparisons t-test. Experiments were repeated at least three times with similar results; results from one experiment are shown. (**G**) Total ROS production in the 60 min following flg22 treatment in *N. benthamiana* leaves 48 h after transient transformation. Error bars indicate SE (n = 24). Asterisks indicate a statistically significant difference (\*\*\**P* < 0.001, \*\*\*\**P* < 0.0001) according to a two-tailed comparisons t-test. Experiments were repeated at least three times with similar results; results from one experiment are shown.

Given that viral proteins from independent origins show a similar coexistence of targeting signals and share the capacity to interfere with SA-dependent defences from the chloroplast, and considering the general effect of SA on different plant-pathogen interactions, we next sought to answer the question of whether unrelated plant pathogens such as bacteria can encode effector proteins with a similar localization pattern and effect. With this purpose, we screened the entire predicted proteome of the plant pathogenic bacterium *Ralstonia solanacearum* GMI1000 for proteins containing both an N-myristoylation motif and a cTP. This search yielded 2 proteins (GALA1 and GALA3) (Figure 4A, Supplementary figure 4A). Interestingly, both proteins have been described as effector proteins secreted inside plant cells during bacterial infection (Mukaihara et al, 2010).

**Figure 4.**
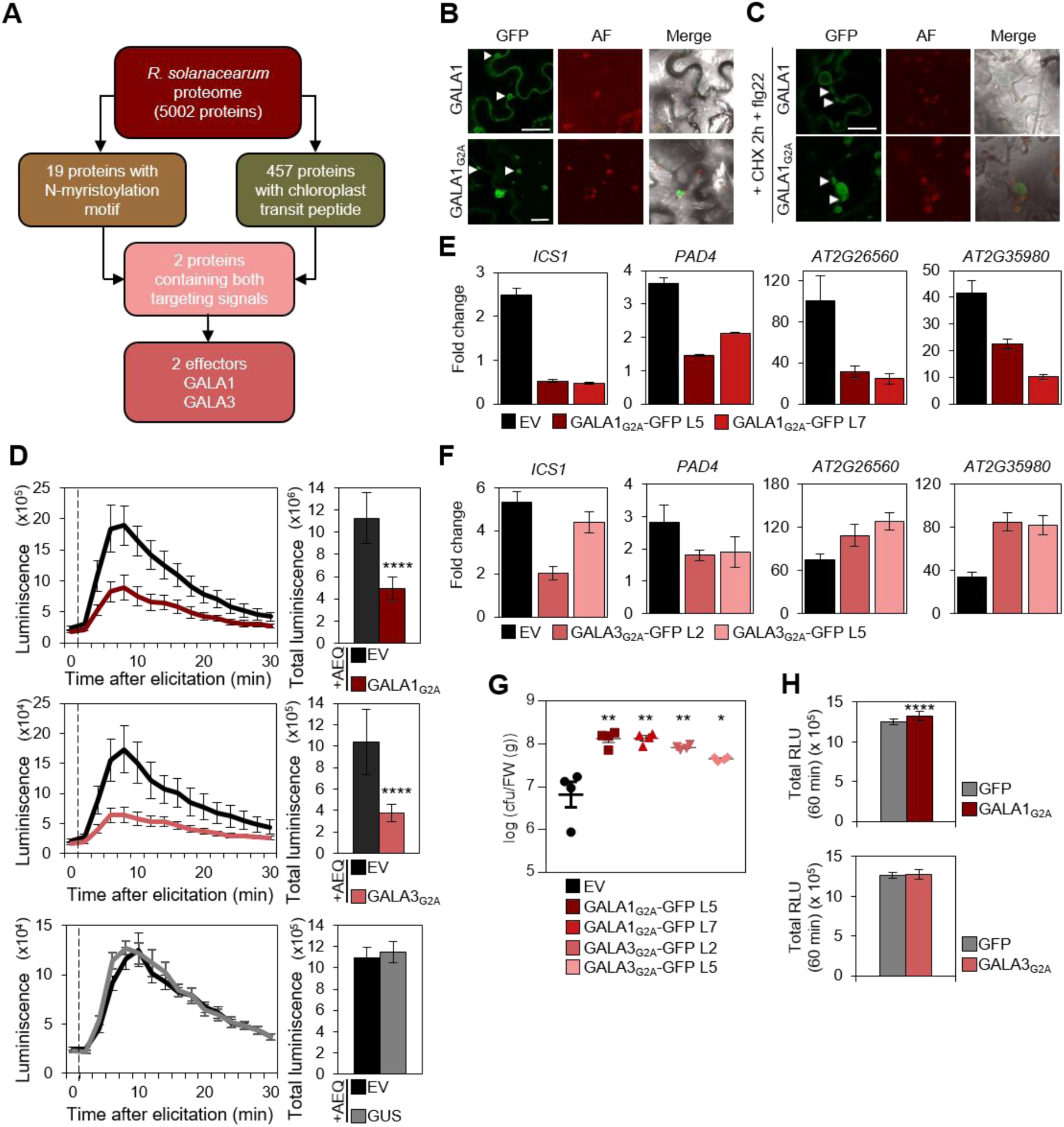
Effector proteins from plant pathogenic bacteria contain overlapping membrane- and chloroplast-targeting signals and suppress chloroplast-mediated defences. (**A**) Flow diagram illustrating the number of proteins encoded by the plant pathogenic bacterial strain *Ralstonia solanacearum* (GMI1000) and containing an N-myristoylation motif, a chloroplast transit peptide, or both. (**B**) The effector GALA1 from *R. solanacearum* GMI1000, which contains overlapping N-myristoylation motif and cTP, shows PM/chloroplast dual localization in *N. benthamiana* leaves. Localization of the wild-type (GALA1) and the non-myristoylable (GALA1_G2A_) versions fused to GFP was observed 2 days post-infiltration. Scale bar = 25 µm. AF: Autofluorescence. Arrowheads indicate chloroplasts. (**C**) GALA1 re-localizes to chloroplasts upon activation of immunity. Localization of the wild-type (GALA1) version fused to GFP was monitored 24 h upon activation of plant immunity by treating with 1 µM flg22. Scale bar = 25 µm. AF: Autofluorescence. Arrowheads indicate chloroplasts. (**D**) Chloroplast accumulation of GALA1 and GALA3 impairs the CAS-dependent cytoplasmic calcium transient in response to flg22. The calcium sensor aequorin (AEQ) was co-expressed with the non-myristoylable (G2A) versions of GALA1 and GALA3 or GUS (as negative control) in *N. benthamiana* leaves. The vertical dashed lines indicate the time at which flg22 treatment was initiated. AEQ luminescence was recorded during 30 min every two minutes (luminiscence in cp 120^−s^). Total luminescence was calculated at the end of the experiment. Values represent the average of 6 plants. Error bars represent SE. Asterisks indicate a statistically significant difference (\*\*\*\**P* < 0.0001) according to a two-tailed comparisons t-test. Experiments were repeated at least three times with similar results; results from one experiment are shown. EV: empty vector. (**E**) Chloroplast-localized GALA1 hampers expression of SA biosynthetic and CAS-dependent genes upon activation of plant immunity. Flg22-induced gene expression was analysed in 10-day-old Arabidopsis transgenic lines expressing GALA1_G2A_-GFP 1 h after flg22 treatment by qRT–PCR. Seedlings were grown under long-day conditions on ½ MS agar plates prior to their transfer to liquid ½ MS for elicitation with 1 µM flg22. *ACT2* was used as the normalizer. Data are mean ± SE of three independent biological replicates with three technical replicates each. EV: empty vector. (**F**) Chloroplast-localized GALA3 hampers expression of SA biosynthetic genes but enhances the expression of CAS-dependent genes upon activation of plant immunity. Flg22-induced gene expression was analysed in 10-day-old Arabidopsis transgenic lines expressing GALA3_G2A_-GFP 1 h after flg22 treatment by qRT–PCR. Seedlings were grown under long day conditions on ½ MS agar plates prior to their transfer to liquid ½ MS for elicitation with 1 µM flg22. *ACT2* was used as the normalizer. Data are mean ± SE of three independent biological replicates with three technical replicates each. EV: empty vector. (**G**) Transgenic Arabidopsis plants expressing chloroplast-localized GALA1 or GALA3 (GALA1_G2A_ and GALA3_G2A_ plants) display increased susceptibility to the bacterial pathogen *P. syringae* pv. *tomato* DC3000. Three-week-old short-day-grown plants were spray-inoculated with *Pto*DC3000. Three days later, bacteria were extracted from 4 independent plants and incubated at 28 °C for 2 days. Data are mean ± SE of n = 4. This experiment was repeated 4 times with similar results; results from one experiment are shown. Asterisks indicate a statistically significant difference (\**P* < 0.05, \*\**P* < 0.01) according to a one-way ANOVA with post-hoc Dunnett’s multiple comparisons test. EV: empty vector. (**H**) Total ROS production in the 60 min following flg22 treatment in *N. benthamiana* leaves 48 h after transient transformation. Error bars indicate SE (n = 24). Asterisks indicate a statistically significant difference (\*\*\*\**P* < 0.0001) according to a two-tailed comparisons t-test. Experiments were repeated at least three times with similar results; results from one experiment are shown.

For subsequent functional characterization, we selected *R. solanacearum* GALA1, since GALA3 could not be detected by confocal microscopy upon transient expression in *N. benthamiana* leaves. GALA1-GFP presented the predicted dual PM/chloroplast localization, and its accumulation in the chloroplast increased upon treatment with flg22 (Figure 4B, C and Supplemental figure 4B). Nevertheless, both bacterial effectors, GALA1 and GALA3, could be detected in chloroplasts upon expression of their G2A version fused to GFP following organelle purification and IP (Supplemental figure 4C). Importantly, chloroplast accumulation of either of these effectors led to a reduction in the cytoplasmic calcium burst in response to flg22 treatment (Figure 4D and Supplemental figure 4D). Since *R. solanacearum* is capable of efficiently infecting Arabidopsis, we generated stable transgenic Arabidopsis plants expressing GALA1_G2A_ or GALA3_G2A_ fused to GFP, which showed the expected chloroplast localization (Supplemental figure 4E, F). These plants display altered expression of SA-responsive genes following flg22 treatment, and are more susceptible to *P. syringae* pv. *tomato* DC3000 (Figure 4E-G and Supplemental figure 4G), indicating that these bacterial effector proteins can disturb SA-dependent defences when localized in the chloroplast. Nevertheless, perception of flg22 is not affected by the expression of the chloroplast-localized versions of these bacterial effectors, since production of apoplastic ROS upon elicitation with the peptide occurs normally (Figure 4H, Supplemental figure 4H).

### The plant-encoded CPK16 employs overlapping targeting signals to re-localize from PM to chloroplasts and promote chloroplast-mediated immune signalling

The finding that pathogens from different kingdoms of life seem to have convergently evolved to target chloroplasts and impair SA-dependent defences following a previous association with membranes raises the idea that a pathway linking PM to chloroplasts might exist in plants, and that such pathway might have been co-opted by different plant pathogens during host-pathogen co-evolution. Following this rationale, we decided to screen the predicted Arabidopsis proteome to identify proteins containing both an N-myristoylation motif and a cTP, and found 78 proteins fulfilling this criterion (Figure 5A; Supplemental table 1). Interestingly, functional enrichment analysis of this subset of proteins unveiled an over-representation of defence regulators and protein kinases (Supplemental figure 5A). We next performed a similar screen using the predicted tomato and rice proteomes, and found 68 and 107 proteins containing both localization signals, respectively (Supplemental figure 5B,C; Supplemental tables 2 and 3), among which defence regulators and protein kinases are also over-represented (Supplemental figure 5D,E). Strikingly, comparison of the identified proteins in Arabidopsis, tomato, and rice yielded a core of 26 homologous proteins common to all three species and containing an N-myristoylation site and a cTP (Table 1; Figure 5B), which shows an over-representation of proteins described as involved in SA signalling and systemic acquired resistance as well as protein kinases (Figure 5C). Of note, the green algae *Chlamydomonas reinhardtii*, the liverwort *Marchantia polymorpha*, and the moss *Physcomitrella patens* also encode proteins containing the N-myristoylation motif and the cTP (19, 30, and 21, respectively; Supplementary figure 5F; Supplementary tables 4, 5, and 6), with a significant over-representation of protein kinases (Supplementary figure 5G-I).

**Table 1.**
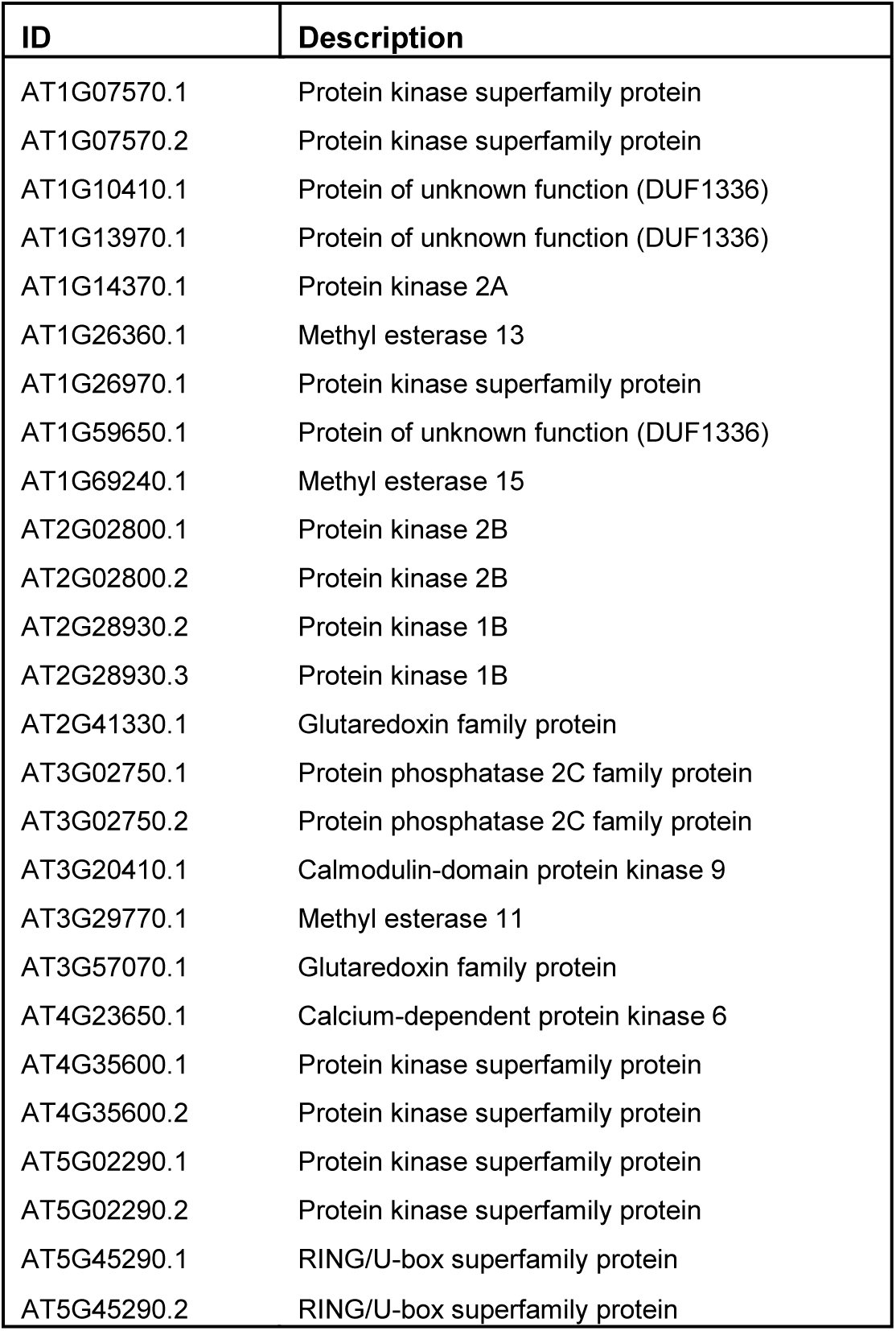
Identity of the 26 proteins corresponding to 19 unique genes common to the Arabidopsis, tomato, and rice proteomes containing both an N-myristoylation motif and a cTP.

**Figure 5.**
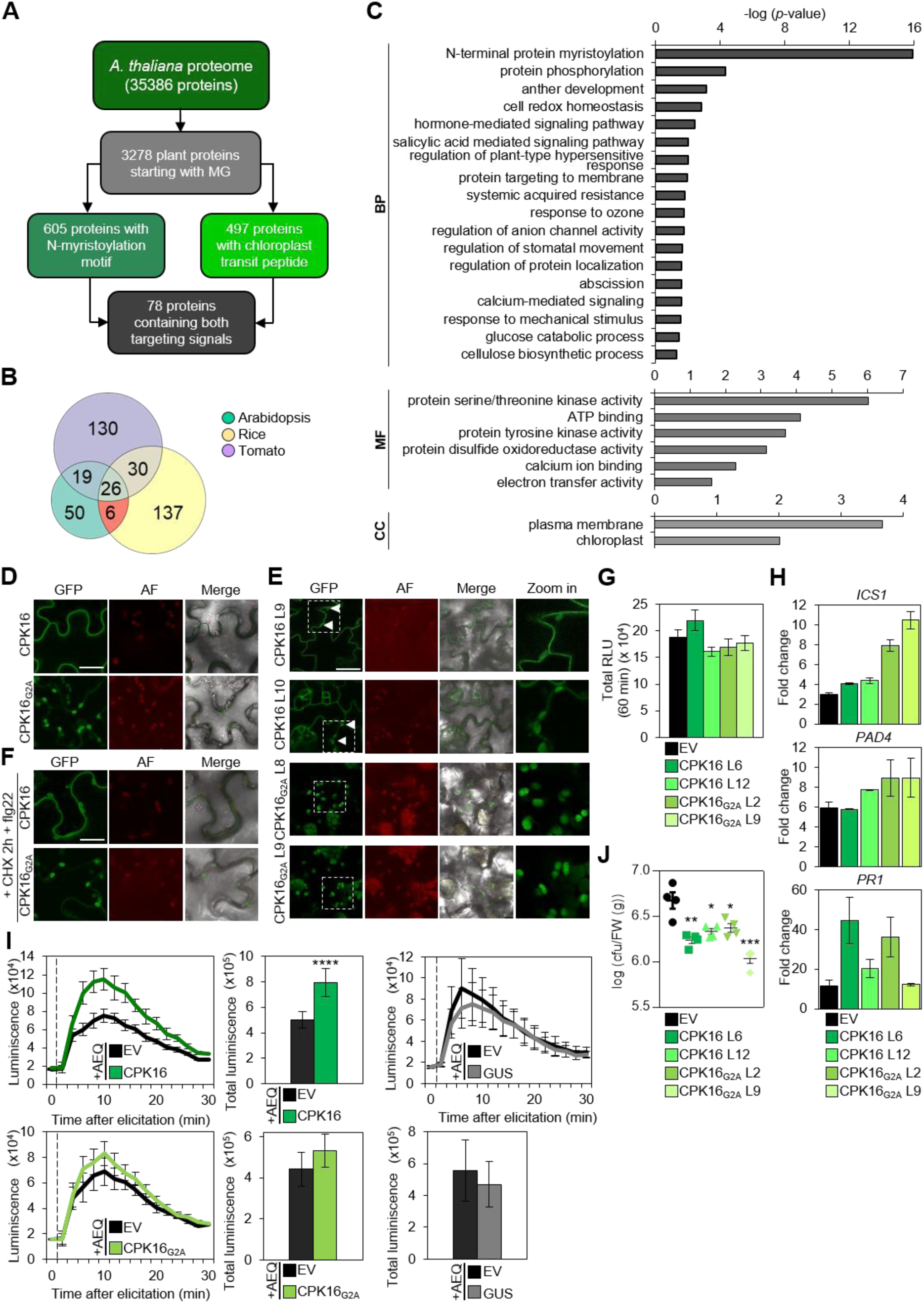
The plant Calcium protein kinase 16 (CPK16) contains overlapping plasma membrane- and chloroplast-targeting signals, shifts its localization from plasma membrane to chloroplasts upon activation of defence, and activates chloroplast-mediated defences. (**A**) Flow diagram illustrating the number of proteins encoded by *Arabidopsis thaliana* containing an N-myristoylation motif, a chloroplast transit peptide, or both. (**B**) Venn diagram of proteins with overlapping N-myristylation motifs and cTPs in the predicted proteomes of Arabidopsis, tomato, and rice. (**C**) The subset of 26 common proteins containing overlapping N-myristylation motifs and cTPs in the predicted proteomes of Arabidopsis, tomato, and rice is enriched in defence-related GO terms. BP: biological process ontology; MF: molecular function ontology; CC: cellular component ontology. (**D**) CPK16 shows PM localization upon transient expression in *N. benthamiana* leaves. Localization of the wild-type (CPK16) and the non-myristoylable (CPK16_G2A_) versions fused to GFP was observed at 2 days post-infiltration. Scale bar = 25 µm. AF: Autofluorescence. (**E**) CPK16 shows PM/chloroplast dual localization in transgenic 5-day-old *A. thaliana.* Localization of the wild-type (CPK16) and the non-myristoylable (CPK16_G2A_) versions fused to GFP is shown. Scale bar = 25 µm. AF: Autofluorescence. Dashed squares indicate region of interest corresponding to the zoom-in panels. Arrowheads indicate chloroplasts. (**F**) CPK16 re-localizes from the PM to the chloroplasts upon activation of PTI in *N. benthamiana* leaves. CPK16 localization was monitored 12 h after activation of PTI by treatment with 1 µM flg22. Scale bar = 25 µm. AF: Autofluorescence. This experiment was repeated three times with similar results. (**G**) Total ROS production in the 60 min following flg22 treatment in *N. benthamiana* leaves 48 h after transient transformation. Error bars indicate SE (n = 16). (*P* = 0.666, according to a one-way ANOVA). Experiments were repeated at least three times with similar results; results from one experiment are shown. EV: empty vector. (**H**) Constitutive chloroplast localization of CPK16 in transgenic CPK16_G2A_ *A. thaliana* plants promotes expression of defence-related marker genes upon activation of defence. Gene expression was analysed 1 h after 1 µM flg22 treatment by qRT–PCR. Seedlings were grown under long-day conditions on ½ MS agar plates prior to their transfer to liquid ½ MS for elicitation. *ACT2* was used as the normalizer. Data are mean ± SE of three independent biological replicates with three technical replicates each. EV: empty vector. (**I**) CPK16 modulates calcium-burst upon activation of plant immunity. The calcium sensor aequorin (AEQ) was coexpressed with the untagged versions of CPK16 or GUS (as negative control) in *N. benthamiana* leaves. The vertical dashed lines indicate the time at which treatment was initiated. AEQ luminescence was recorded during 30 min every two minutes (luminiscence in cp 120^−s^). Total luminescence was calculated at the end of the experiment. Values represent the average of 6 plants. Error bars represent SE. Asterisks indicate a statistically significant difference (\*\*\*\**P* < 0.0001) according to a two-tailed comparisons t-test. Experiments were repeated at least three times. Representative data are shown. EV: empty vector. (**J**) Transgenic Arabidopsis plants overexpressing CPK16 or CPK16_G2A_ display increased resistance against the bacterial pathogen *P. syringae* pv. *tomato* DC3000. Three-week-old short-day-grown plants were spray-inoculated with *Pto*DC3000. Three days later, bacteria were extracted from 4 independent plants and incubated at 28 °C for 2 days. Data are mean ± SE of n = 4. This experiment was repeated 4 times with similar results; results from one experiment are shown. Asterisks indicate a statistically significant difference (\**P* < 0.05, \*\**P* < 0.01, \*\*\**P* < 0.001) according to a one-way ANOVA with post-hoc Dunnett’s multiple comparisons test. EV: empty vector.

Among the proteins identified in Arabidopsis, one of them, Calcium-Dependent Kinase 16 (CPK16), had already been shown to associate to the PM in an N-myristoylation motif-dependent manner and to harbour a functional cTP (Stael et al., 2011); however, chloroplast localization could be observed only upon mutation of the myristoylation site, and hence its biological significance was unclear. Following transient expression in *N. benthamiana*, CPK16-GFP could be observed mostly at the PM, while, as previously described, a CPK16_G2A_ mutant version was detected in chloroplasts (Figure 5D); stable expression in Arabidopsis yielded similar results (Figure 5E). However, after flg22 treatment, CPK16-GFP accumulated in chloroplasts, similarly to C4-GFP (Figure 5F, Supplemental figure 5J); chloroplast localization could still be detected after CHX treatments, indicating that this process does not require new protein synthesis, and therefore the chloroplast-localized protein most likely derive from the PM pool (Figure 5F).

As proof-of-concept, we decided to investigate whether CPK16 plays a role in PTI responses when localized in the chloroplast. For this purpose, we generated transgenic Arabidopsis lines expressing wild-type CPK16 or its chloroplast-localized CPK16_G2A_ version under the 35S promoter (Supplemental figure 5K); these transgenic lines had no obvious developmental phenotype (Supplemental figure 5L). Following flg22 treatment, both CPK16 and CPK16_G2A_ plants displayed normal ROS burst but enhanced expression of SA marker genes (*ICS1*, *PAD4*, and *PR1*), an effect that was more pronounced for early-expressing genes when the chloroplastic version of CPK16 was used (Figure 5G, H; Supplemental figure 5M). Although a moderate increase in the amplitude of the flg22-triggered cytoplasmic calcium burst could be detected in *N. benthamiana* as the result of CPK16 overexpression, CPK16_G2A_ had no evident effect on this readout (Figure 5I, Supplemental figure 5N). Importantly, both CPK16- and CPK16_G2A_-overexpressing lines are also more resistant to *P. syringae* pv tomato DC3000 (Figure 5J, Supplementary figure 5O), indicating that CPK16 contributes to anti-bacterial immunity from the chloroplast.

## DISCUSSION

It has become widely recognized that, in addition to their function as energy providers, chloroplasts play an essential role in different aspects of plant biology as stress sensors and signal integrators; however, how these organelles, which reside inside the cell separated from the cytosol by a double membrane, can sense external cues is a long-standing question.

Taken together, the results presented here suggest the existence of a novel conserved pathway in plants that physically connects the PM and chloroplasts through protein re-localization upon a specific trigger, namely the perception of a biotic threat (Figure 6). This pathway, exemplified by CPK16, would involve proteins harbouring two overlapping and conflicting targeting signals: an N-myristoylation site, tethering the protein to the PM, and a cTP, targeting the protein to the chloroplast following release from the PM. Remarkably, the Arabidopsis N-myristoylome is enriched in defence-related proteins (Boisson et al., 2003). Our results show that CPK16 localizes to the PM in basal conditions, as previously observed (Stael et al., 2011), but can re-localize to chloroplasts following activation of PTI; chloroplast-localized CPK16 enhances chloroplast-dependent PTI responses, indicating that this protein can act as a modulator of defence through its function in the chloroplast. Many other Arabidopsis proteins containing both targeting signals are known defence regulators (e.g. BIK1, BSK1, AGI1, CPK28); whether these and the other proteins in this subset localize in the chloroplast under specific conditions, and if so, what their function in the organelle and their contribution to defence responses are remains to be investigated. Of note, the subsets of proteins containing these two targeting signals present an over-representation of protein kinases in all species analysed (*C. reinhardtii*, *M. polymorpha*, *P. patens*, *A. thaliana*, *O. sativa*, and *S. lycopersicum*), suggesting that kinases that work in other subcellular compartments can function in the chloroplast upon perception of certain stimuli, and underscoring the potential relevance of protein phosphorylation for signalling relay in this putative pathway.

**Figure 6.**
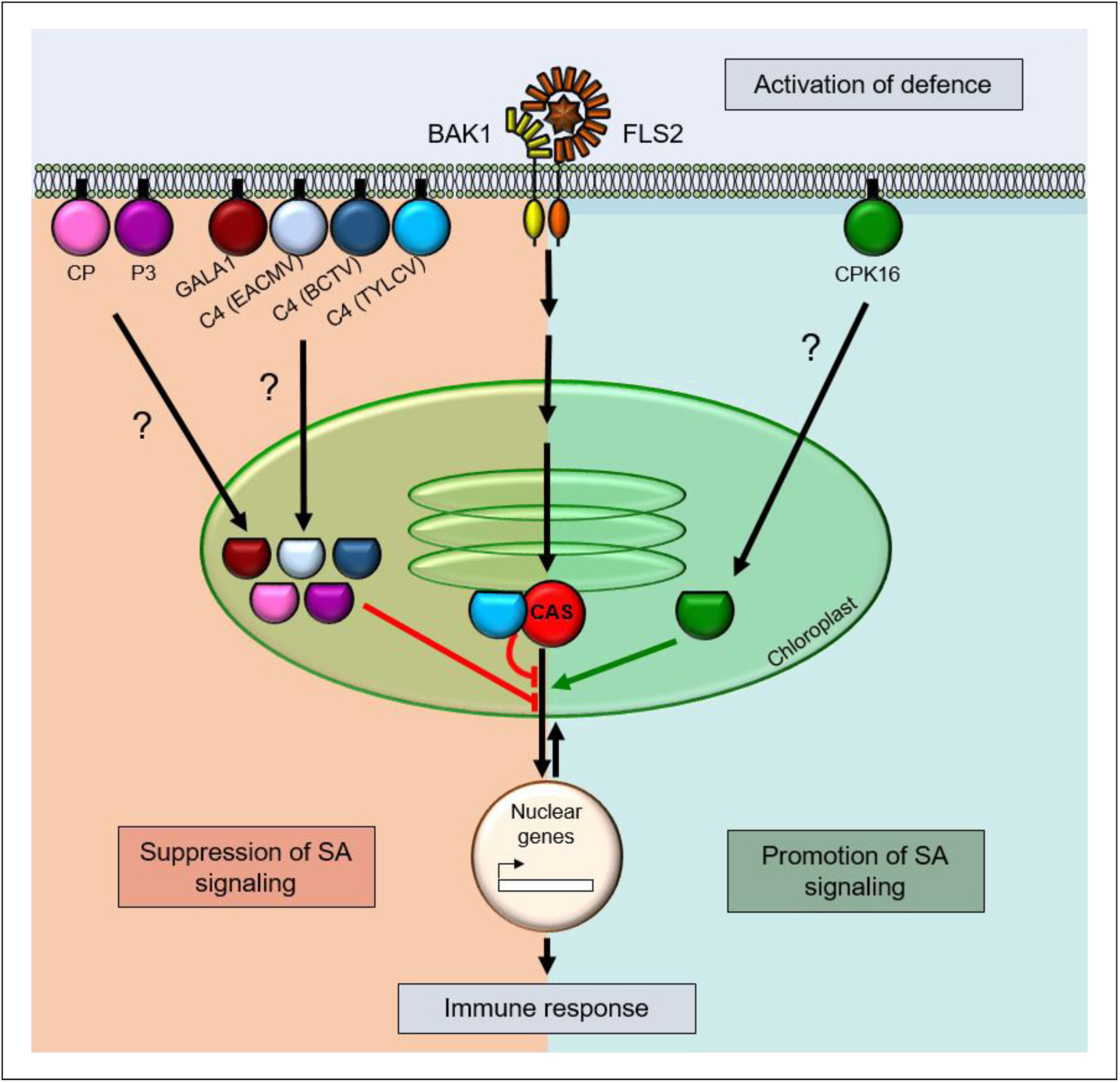
Model of the proposed pathway linking plasma membrane to chloroplasts and activating defence in plants and its co-option by plant pathogens to promote virulence through suppression of SA responses. On the right, the plant defence regulator CPK16; on the left, pathogen-encoded proteins (CP: CP protein from the cucumovirus *Cucumber mosaic virus* (Fam. *Bromoviridae*); P3: P3 protein from the nepovirus *Grapevine fanleaf virus* (Fam. *Secoviridae*); GALA1: GALA1 effector from *R. solanacearum* GMI1000; C4 (EACMV): C4 protein from the bipartite begomovirus *East African cassava Mosaic virus* (Fam. *Geminiviridae*); C4 (BCTV): C4 protein from the curtovirus *Beet curly top virus* (Fam. *Geminiviridae*); C4 (TYLCV): C4 from the monopartite begomovirus *Tomato yellow leaf curl virus* (Fam. *Geminiviridae*); CAS: Plant-encoded Calcium sensing receptor protein). Activation of defence following perception of a biotic threat at the plasma membrane would trigger release of the indicated proteins and their subsequent re-localization to chloroplasts, where they would activate (green arrow) or repress (red arrows) downstream SA-dependent defence responses.

Strongly supporting the notion of this pathway having a central function in plant defence is the finding that its co-option and potential disruption by pathogens has evolved multiple times independently: effectors containing an N-myristoylation motif and a cTP and suppressing chloroplast-dependent defences from this organelle have been identified as encoded by DNA viruses, RNA viruses, and bacteria. Future work will unveil how prevalent the coexistence of these two signals is among pathogen effectors, including those belonging to other kingdoms of life.

The pathway proposed here would allow for timely, rapid, and precise information relay from the PM to the chloroplast, fine-tuning retrograde signalling to orchestrate appropriate responses upon signal integration in the plastid, avoiding negative consequences of activating defence and shutting down photosynthesis in the absence of a threat. Additionally, it could allow for dual functions of the participating proteins, since they could exert a given role in basal conditions, at the PM, and a different one once a biotic threat is perceived and they are translocated into the chloroplast – for example, by keeping signalling in an off state in the absence of stimulus, but promoting downstream responses upon pathogen perception.

One example of how the subcellular compartmentalization enabled by protein re-localization can give rise to multifunctionality is illustrated by the C4 protein encoded by TYLCV. At the PM, C4 interacts with the receptor-like kinases BAM1 and BAM2 and hinders the intercellular spread of RNAi (Rosas-Diaz et al., 2018; Fan et al., 2019); following activation of defence, however, which is triggered by the presence or activity of the virus-encoded Rep protein, C4 is translocated to the chloroplast, where it interacts with CAS and interferes with the CAS-dependent defence responses, including SA biosynthesis. Therefore, C4 can exert at least two distinct biological functions depending on its subcellular localization, which is determined by the state of the cell, both of which promote viral pathogenicity. This model could also apply to other pathogen effectors, which could have additional virulence-promoting roles at the PM in the absence of a defensive trigger. The identification of the targets of independently evolved effectors following chloroplast translocation could make a powerful contribution to elucidating the molecular mechanisms involved in chloroplast-nucleus communication during plant-pathogen interactions.

In addition to expanding our view of the molecular and cellular underpinnings of plant defence, the unravelling of the direct physical connection between PM and chloroplasts following the perception of a biotic threat could also have practical applicability, paving the way to the engineering of improved resistance to pathogens without yield penalty. This idea is suggested by the results obtained with transgenic CPK16_G2A_ Arabidopsis plants, which show no apparent developmental phenotype, with no effect in growth, but are more resistant to bacterial infection. This observation also implies that chloroplast localization is not sufficient for CPK16 to activate defence responses in the absence of a trigger, and that some additional component, regulated by perception of a biotic threat, is required for the downstream effects.

An intriguing question is how these myristoylated, PM-localized proteins get released in order to be translocated to the chloroplast. Since myristoylation is an irreversible covalent lipidation, an additional post-translational modification would be required for the release to occur. In animals, myristoylation-dependent PM association has been shown to be regulated by phosphorylation of the protein (Thelen et al., 1991); since phosphorylation is a prevalent event during activation of defence signalling at the PM, this modification might be a plausible candidate to modulate the localization of plant defence regulators or mimicking pathogen effectors. As the results with the non-myristoylable versions of these plant- or pathogen-encoded proteins indicate, PM release would be sufficient to guarantee immediate chloroplast localization.

Multiple environmental and developmental cues are perceived at the PM. Considering their central role in plant biology, it would be conceivable that some of these signals get relayed to chloroplasts in order to coordinate appropriate downstream physiological responses. In this scenario, whether a similar, post-translational modification-based mechanism operates to enable communication between the PM and chloroplasts following perception of non-defence related cues is a question that remains to be explored.

## METHODS

### Transient expression in *Nicotiana benthamiana*

Transient expression assays were performed as described in Wang et al. (2017a) with minor modifications. In brief, the *Agrobacterium tumefaciens* strain GV3101 harbouring the corresponding binary vectors were liquid cultured in LB with the appropriate antibiotics at 28°C overnight. Bacterial cultures were centrifuged at 4,000 g for 10 min and resuspended in the infiltration buffer (10 mM MgCl_2_, 10 mM MES pH 5.6, 150 µM acetosyringone) to an OD_600_ = 0.2-1. Bacterial suspensions were incubated at room temperature in the dark for 2 h before infiltration into the abaxial side of 4-week-old *N. benthamiana* leaves with a 1 mL needle-less syringe. For experiments that required co-infiltration, the *Agrobacterium* suspensions carrying different constructs were mixed at 1:1 ratio before infiltration.

### Chloroplast isolation

Chloroplasts were isolated from 3-week-old Arabidopsis and 4-week-old *N. benthamiana* plants as described previously (Kauss et al., 2012). Isolated chloroplasts were separated into membrane and stroma fractions according to (Wang et al., 2016). During the course of the fractionation, SIGMAFAST^TM^ Protease Inhibitor was added to all required buffers (1 tablet per 300 ml buffer). Chlorophyll from separated membrane fractions was removed using acetone (Wang et al., 2016). The resulting proteins from pellet and stroma fractions were resuspended in 1x Laemmli SDS sample buffer and denatured for 10 min at 95 °C.

Equal amounts of the solubilized protein fractions were separated on 10% SDS-PAGE gels and blotted onto PVDF membranes (Bio-Rad). RFP-fused proteins were detected using a rat anti-RFP antibody (1:5,000; ChromoTek). Light harvesting complex protein (LhcB1) and Rubisco large subunit (RbcL) proteins were immunochemically detected using rabbit anti-LhcB1 (1:10,000; Agrisera) and rabbit anti-RbcL (1:10,000; Agrisera) antibodies, respectively.

To detect GALA1_G2A_ and GALA3_G2A_, chloroplasts were isolated from 4-week-old *N. benthamiana* plants expressing GALA_G2A_-GFP proteins (2 dpi) as described previously (Kauss et al., 2012). GFP-Trap beads (GFP-Trap Agarose; ChromoTek) were added to the isolated chloroplast proteins. The samples were first agitated at room temperature for 1.5-2 h, then the beads were washed 4 times for 5 min with wash buffer. Finally, the washed beads were resuspended in 100μl 1x Laemmli SDS sample buffer and incubated for 20 min at 70 °C. Equal amounts of the eluted proteins were separated on 10% SDS-PAGE gels and blotted onto PVDF membranes (Bio-Rad). GFP-fused proteins were detected using mouse monoclonal anti-GFP antibody (1:5,000; Abiocode).

### Thylakoid membrane isolation and topology analysis

Thylakoid membranes were isolated from 4-week-old plants as described by Kato et al. (2018). Leaves were homogenized in a blender with ice-cold homogenization buffer (0.35 M Sucrose, 50 mM HEPES pH 7.5, 0.5 mM MgCl_2_, 10 mM NaCl, and 2 mM EDTA). Homogenates were then filtered through Miracloth (Merck Millipore). After centrifugation at 2,380 *× g* for 10 min, the pellet was resuspended in the same buffer. After centrifugation at 300 *× g* for 1 min, the supernatant was centrifuged at 2,380 *× g* for 10 min. The pellets were resuspended in the homogenization buffer and used for further analyses.

For topology analysis, freshly isolated thylakoid membranes were resuspended (0.5 mg chlorophyll/ml) in HS buffer (0.35 M Sucrose, 50 mM HEPES, pH 7.5 mM) and treated with thermolysin (100 µg/ml) for 30 min on ice. The soluble- and pellet-fractions were then separated by centrifugation at 2,380 *× g* for 5 min. Chlorophyll from pellet fractions was removed using acetone. The resulting proteins from pellet and soluble fractions were resuspended in 1x Laemmli SDS sample buffer and denatured for 10 min at 95 °C. Equal amounts of the solubilized protein fractions were separated on 10% SDS-PAGE gels and blotted onto PVDF membranes (Bio-Rad). Protochlorophyllide oxidoreductase (POR) and photosystem II subunit O (PsbO) proteins were immunochemically detected using rabbit anti-POR (1:5,000; Agrisera) and rabbit anti-PsbO (1:10,000; Agrisera) antibodies, respectively.

### Co-immunoprecipitation (co-IP) and western blot

To detect the interaction between C4 and CAS, thylakoid membranes were first isolated to extract proteins. Co-immunoprecipitation (co-IP) was carried out as described previously (Dogra et al., 2019; Wang et al., 2016). Briefly, freshly isolated thylakoid membranes were resuspended in co-IP buffer and kept on ice for 20 min followed by centrifugation at 21,000 *× g* for 30 min at 4 °C. The supernatant was filtered through a 0.22 μm Millipore Express PES membrane and proteins were quantified using Pierce^TM^ BCA protein assay kit (Thermo Fisher Scientific). Small aliquots were taken as input samples whereas the remaining parts were used for co-IP. RFP-Trap beads (RFP-Trap Agarose; ChromoTek) were added to equal amounts of the co-IP samples. The samples were first agitated at room temperature for 1.5-2 h, then the beads were washed 4 times for 5 min with wash buffer. Finally, the washed beads were resuspended in 100μl 1x Laemmli SDS sample buffer and incubated for 20 min at 70 °C. Equal amounts of the eluted proteins were separated on 10% SDS-PAGE gels and blotted onto PVDF membranes (Bio-Rad). GFP- and RFP-fused proteins were detected using mouse monoclonal anti-GFP antibody (1:5,000; Abiocode) and rat monoclonal anti-RFP antibody (1:5,000; Chromotek), respectively.

### Calcium burst measurements

*N. benthamiana* leaf discs transiently co-expressing aequorin and the construct of interest were floated on Milli-Q water containing 5 µM coelenterazine (Sigma), then kept overnight at 22°C in the dark. A 1 µM solution of the elicitor peptide flg22 was applied to the leaf discs and the transient increase in calcium was immediately recorded. Aequorin luminescence was measured with a NightShade LB 985 In vivo Plant Imaging System (Berthold) equipped with an absolutely light-tight cabinet and a cooled CCD camera; data were analysed with Indigo v2 software.

### Bacterial infections

Three-week-old Arabidopsis plants grown under short-day condition were spray-inoculated with a *Pseudomonas syringae* pv. *tomato* DC3000 inoculum (OD_600_ = 0.2 in 10 mM MgCl_2_ with 0.02% Silwet L-77) and kept covered for 24 h. Four-week-old Arabidopsis plants grown under short-day condition were infiltrated with a *Pto*DC3000 inoculum (OD_600_ = 0.0002 in 10 mM MgCl_2_) using a needleless syringe and kept covered for 24 h.

Four-week-old *N. benthamiana* leaves were infiltrated with a *P. syringae* pv. *tomato* DC3000 *∆hopQ1-1* suspension (OD_600_ = 0.0002 in 10 mM MgCl_2_) using a needleless syringe upon transient expression of the construct of interest.

Bacterial growth was determined three days after inoculation by plating 1:10 serial dilutions of leaf extracts; plates were incubated at 28 °C for two days before the bacterial cfu were counted.

### Bioinformatic analyses

For the identification of N-myristoylation motifs and chloroplast transit peptides (cTPs) in the predicted proteomes analyzed in this work, detailed information is provided in the Supplemental methods section.

In all cases, the N-terminal myristoylation motif and the cTP were predicted by Expasy Myristoylator (https://web.expasy.org/myristoylator/) and ChloroP (http://www.cbs.dtu.dk/services/ChloroP/), respectively.

GO term analyses were conducted with topGO (https://bioconductor.org/packages/release/bioc/html/topGO.html); Venn diagrams were drawn by eulerr (https://cran.r-project.org/web/packages/eulerr/index.html).

## ACKNOWLEDGEMENTS

The authors thank Xinyu Jian, Aurora Luque, Tamara Jimenez-Gongora, Yujing (Ada) Liu, the PSC Proteomics and Metabolomics Core Facility, the PSC Cell Biology Core Facility, and the PSC Genomics Core Facility for technical assistance, Jose Rufian for experimental advice, and all members in Rosa Lozano-Duran’s and Alberto Macho’s groups for stimulating discussions and helpful suggestions. The authors thank Jiamin Luo and Alberto Macho for sharing the construct 35S:*GUS-3xHA*, and are grateful to Alberto Macho and Tamara Jimenez-Gongora for critical reading of the manuscript. This research was supported by the the National Natural Science Foundation of China (NSFC) (grant numbers 31671994 and 31870250 to RL-D), the 100 Talents Program from the Chinese Academy of Sciences (CAS) (to RL-D), the Shanghai Center for Plant Stress Biology, and a Young Investigator Grant from the NSFC to LM-P (grant number 31850410467). LM-P is the recipient of a President’s International Fellowship Initiative (PIFI) postdoctoral fellowship (No. 2018PB058) from CAS.

## AUTHOR CONTRIBUTIONS

RL-D conceived the project; LM-P, HT, VD, MW, TR-D, LW, XD, and DZ performed experiments; LM-P, VD, FX, and RL-D analysed data. RL-D wrote the manuscript with input from all authors.

## SUPPLEMENTAL INFORMATION

## SUPPLEMENTAL METHODS

## SUPPLEMENTAL FIGURES

**Supplemental figure 1. C4 shifts its localization from the plasma membrane to the chloroplast following activation of PTI.**

**Supplemental figure 2. C4 interacts with CAS in the chloroplast and suppresses SA-dependent defences.**

**Supplemental figure 3. Proteins encoded by plant viruses and containing overlapping N-myristoylation motifs and cTPs suppress defence responses from the chloroplast.**

**Supplemental figure 4. Proteins encoded by the plant pathogenic bacterium *Ralstonia solanacearum* and containing overlapping N-myristoylation motifs and cTPs suppress defence responses from the chloroplast.**

**Supplemental figure 5. A core of conserved plant defence-related proteins contain overlapping N-myristoylation motifs and cTPs.**

## SUPPLEMENTAL TABLES

**Supplemental table 1.** Identity of the 78 proteins containing both an N-myristoylation motif and a cTP from the Arabidopsis predicted proteome.

**Supplemental table 2.** Identity of the 68 proteins containing both an N-myristoylation motif and a cTP from the predicted *Solanum lycopersicum* proteome.

**Supplemental table 3.** Identity of the 107 proteins containing both an N-myristoylation motif and a cTP from the predicted *Oryza sativa* subsp. *japonica* proteome.

**Supplemental table 4.** Identity of the 19 proteins containing both an N-myristoylation motif and a cTP from the predicted *Chlamydomonas reinhardtii* proteome.

**Supplemental table 5.** Identity of the 30 proteins containing both an N-myristoylation motif and a cTP from the predicted *Marchantia polymorpha* proteome.

**Supplemental table 6.** Identity of the 21 proteins containing both an N-myristoylation motif and a cTP from the predicted *Physcomitrella patens* proteome.

**Supplemental table 7.** Plasmids and constructs used in this work.

**Supplemental table 8.** Primers used in this work.

