## Supplemental material for "A novel pathway linking plasma membrane and chloroplasts is co-opted by pathogens to suppress salicylic acid-dependent defences"

Laura Medina-Puche *et al*.

### SUPPLEMENTAL METHODS

#### Plant material

All Arabidopsis mutants and transgenic plants used in this work are in the Col-0 background. The Arabidopsis 35S:*C4* lines are described in Rosas-Diaz et al. (2018). To generate Arabidopsis transgenic lines, wild-type *Arabidopsis* plants were transformed with the corresponding construct (see Supplemental table 5) using the floral dipping method (Zhang et al., 2006; Clough, 2005; Clough and Bent, 1998). Arabidopsis plants were grown in a controlled growth chamber in long-day conditions (16 h light/8 h dark) at 22°C for all experiments except for bacterial infections and callose deposition, for which plants were grown in a controlled growth chamber in short-day conditions (8 h light/16 h dark) at 22°C. *Nicotiana benthamiana* and tomato (cv. Moneymaker) plants were grown in a controlled growth chamber in long-day conditions (16 h light/8 h dark) at 25°C.

#### Plasmids and cloning

Plasmids and primers used for cloning are summarized in Supplemental tables 7 and 8. The TYLCV clone used as template is AJ489258 (GenBank). The TYLCV_$C4_{1-8}$ infectious clone is described in Rosas-Diaz et al. (2018). Vectors from the pGWB and ImpGWB series were kindly provided by Tsuyoshi Nakagawa (Nakagawa et al., 2007a; Nakagawa et al., 2007b). pB7RWG2.0 is described in Karimi et al. (2002). pGTQL1211YN and pGTQL1221YC are described in Lu et al. (2010). The pBIN-CYA(K) plasmid expressing $AEQ_{cyt}$ is described in Mehlmer et al. (2012). In all cases, the pENTR™/D-TOPO® entry vector (Thermo Scientific) or the pDONR™221 entry vector (Thermo Scientific) containing the fragments of interest were recombined into the corresponding destination vectors through the Gateway LR reaction (Thermo Scientific).

**qPCR and qRT-PCR**

RNA was extracted using the Plant RNA kit (OMEGA Bio-Tek); cDNA was prepared using the iScript™ gDNA Clear cDNA Synthesis Kit (Bio-Rad) according to the manufacturer's instructions. DNA was extracted with 2xCTAB from leaf tissues. DNA and cDNA were analyzed by qPCR with Hieff™ qPCR SYBR® Green Master Mix (Yeasen). The reactions were done as follows: 3 min at 95 °C, 40 cycles consisting of 15 s at 95 °C, 30 s at 60 °C. The primers used are described in Supplemental table 8. *ACTIN* (*ACT2*) was used as internal reference for DNA extracted from Arabidopsis samples, and the *25S ribosomal DNA interspacer* (*ITS*) was used for *N. benthamiana* and tomato DNA samples (Mason et al., 2008). For cDNA samples, *ACT2*, *elongation factor-1 alpha* (*NbEF1α;* Nicot et al., 2005), and *SlACTIN* were used as normalizers for Arabidopsis, *N. benthamiana*, and tomato, respectively.

**RNA-sequencing**

Transcriptome analyses were performed at the Genomics Core Facility, Shanghai Center for Plant Stress Biology, CAS. Four biological replicates of each genotype were used. Total RNA (1 µg) from each sample was used for library preparation with NEBNext® Poly(A) mRNA Magnetic Isolation Module (New England BioLabs) and NEBNext Ultra II Directional RNA Library Prep Kit for Illumina (New England BioLabs) following the manufacturer's instructions. Prepared libraries were assessed for quality using NGS High-Sensitivity kit on a Fragment Analyzer (AATI) and for quantity using Qubit 2.0 fluorometer (Thermo Fisher Scientific). All libraries were sequenced in paired-end 150 bases protocol (PE150) on an Illumina HiSeq sequencer.

**Affinity-purification and mass spectrometry**

Affinity-purification and mass spectrometry (AP-MS) was performed according to Wang et al. (2017b). Data were analysed by Scaffold 4.9 with 99% peptide threshold.

**Total SA quantification**

25 mg of ten-day-old Arabidopsis seedlings were used for SA quantification. Total SA was extracted with 70% MeOH and 2 ng SA-d4 was used as internal standard. 50 µL of the solution were injected into the LC-MS that was performed in a Waters Liquid chromatography ACQUITY

UPLC I-class coupled with AB SCIEX TripleTOF® 5600+ equipped with an ACQUITY UPLC BECH C18 1.7 µm VanGuardTM Pre-Column 2.1x5 mm column. The analytical column used was ACQUITY UPLC BECH C18 1.7 µm 2.1x150 mm column. The results were analyzed by Peakview 1.2.

**Seedling growth inhibition**

Sterilized seeds were sown on ½ MS medium containing 1% sucrose and 0.7% agar. Seeds were stratified for 3 days in the dark at 4°C and grown in long-day conditions. Five-day-old seedlings were transferred into 48-well plates (Corning™ Costar™, Fisher Scientific) with liquid ½ MS containing 0.8% sucrose with or without the desired peptides (elf18 or flg22 at 10 nM and 100 nM concentration) and grown for another ten days. Fresh weight of 12-16 seedlings was measured using a precision balance.

**Reactive oxygen species measurement**

Measurement of PAMP-triggered reactive oxygen species (ROS) was performed as previously described (Sang and Macho, 2017). Leaf discs from 4- to 5-week-old Arabidopsis plants grown in short-day conditions were placed in white 96-well plates (OptiPlate™-96, PerkinElmer) with water overnight. The following day, the water was replaced by a solution containing 100 nM flg22, 100 µM luminol, and 20 µg/mL HRP, and luminescence was measured in a microplate reader (Varioskan flash, Thermo Scientifc) for 60 min. For data analysis, both relative luminescence units (RLU) produced every minute upon flg22 treatment and total RLU over the duration of 60 min were plotted.

**Callose deposition**

Three- to four-week-old Arabidopsis plants grown in short-day conditions were infiltrated with 1 µM flg22 solution or water using a needle-less syringe. Infiltrated leaves were collected 24 h later and placed in 6-well plates. Chlorophyll was removed by incubating the tissue in ethanol (90% v/v) at 37 °C, followed by two washes with ethanol (75% v/v) for 1 h, and one in $dH_2O$ for 1h. The distained leaves were vacuum-infiltrated with 0.05% aniline blue and incubated for 1h. Then, aniline blue was removed and leaves were embedded in 50% glycerol and conserved at 4 °C in the dark before visualization. Callose deposits were visualized under UV illumination (Ex: 405 nm, Em: 448–525 nm) and quantified using Image J software.

#### Transmission electron microscopy

10-day-old Arabidopsis seedlings were used for TEM observation. Samples were cut into sections of about 1mm×2mm and pre-fixed in 2.5% glutaraldehyde in 0.1M phosphate buffer (PBS, pH7.4) at 4°C for 3 days, followed by 3 washes with PBS buffer. Samples were post-fixed in 1% osmic acid overnight at 4°C, followed by 3 washes with 0.1M PBS, then dehydrated with a gradient ethanol-acetone series and gradient-infiltrated and embedded in Spurr resin. Ultra-thin resin sections (70 nm) were cut by a diamond knife on a Leica UC7 ultramicrotome, mounted on copper grids, and stained by 2% uranyl acetate and 0.5% lead citrate. The sections were observed and photographed using a transmission electron microscope H7700 (Hitachi, Japan).

#### Determination of maximum photochemical efficiency of photosystem II (Fv/Fm)

The Fv/Fm was determined with a FluorCam system (FC800-C/1010GFP; Photon Systems Instruments) containing a CCD camera and an irradiation system following the manufacturer's instructions. Seedlings were grown in ½ MS -sucrose agar medium under constant light (low light intensity: 40 µmol $m^{-2}$ $s^{-1}$; high light intensity: 80 µmol $m^{-2}$ $s^{-1}$) for 5 days under normal temperature (22°C), followed by 5 days in constant light (very high intensity: 300 µmol $m^{-2}$ $s^{-1}$) under cold stress conditions (10°C), when Fv/Fm was recorded every day (D: day; D0 to D5: from day 0 to day 5); seedlings were then transferred to the initial conditions, and Fv/Fm was recorded at the indicated time points (R: recovery; R_2h: 2 h after recovery; R_D1 to R_D5: from day 1 to day 5 after recovery).

#### Visualization of protein subcellular localization

For subcellular localization, plant tissues expressing GFP- or RFP-fused proteins were imaged with a Leica TCS SP8 confocal microscope (Leica Microsystems) using the preset settings for GFP (Ex: 488 nm, Em: 500-550 nm) and RFP (Ex: 554 nm, Em: 580-630 nm).

**Bimolecular fluorescence complementation (BiFC) assay**

BiFC assays were performed with a split-YFP system in *N. benthamiana* leaves as described previously (Lu et al., 2010). *A. tumefaciens* mixtures carrying the appropriate BiFC constructs were infiltrated into the abaxial side of 4-week-old *N. benthamiana* leaves with a 1 mL needle-less syringe. Two days later, samples were imaged with a Leica TCS SP8 confocal microscope (Leica Microsystems) using the preset settings for YFP (Ex: 514 nm, Em: 525-575 nm).

**Viral infections**

Viral infections were performed as described in Lozano-Duran et al. (2011a, b). In brief, for *N. benthamiana* and tomato (cv. Moneymaker), a suspension of *Agrobacterium* carrying the TYLCV infectious clone or a mutant version lacking C4 (TYLCV_C4$_{1-8}$) were syringe-inoculated in the shoot apical meristem of 2-week-old plants. For Arabidopsis, the corresponding *Agrobacterium* cells were collected from densely grown plates by scraping and inoculated in the centre of the rosette using a needle.

**Virus induced gene silencing (VIGS) assay**

VIGS assays were performed as described by Yu et al. (2019). Briefly, independent cultures of *A. tumefaciens* GV3101 carrying pTRV1 or pTRV2-based constructs were grown overnight in LB medium plus appropriate antibiotics. Cultures were resuspended in VIGS buffer (10 mM morpholineethanesulfonic acid pH 5.6, 10 mM $MgCl_2$, and 100 µM acetosyringone) to $OD_{600}=1$, and incubated 2 h at room temperature in the dark. Cultures were mixed at a 1:1 ratio. Approximately 1 mL of this suspension was used to inoculate the stem and underside of cotyledons of 2-week-old *N. benthamiana* and tomato (cv. Moneymaker) plants. Samples were collected 3 weeks after inoculation.

**Prediction of N-myrisotylation motif and chloroplast transit peptide**

In all cases, the N-terminal myristoylation motif and the cTP were predicted by Expasy Myristoylator (https://web.expasy.org/myristoylator/) and ChloroP (http://www.cbs.dtu.dk/services/ChloroP/), respectively. For plant viruses, we extracted the "Genbank organism" corresponding to plant virus from the tab-delimited text file downloaded from

DPV (www.dpvweb.net/seqs/seqlist.zip) and used NCBI Entrez Direct (https://www.ncbi.nlm.nih.gov/books/NBK179288/) to download all the plant virus protein sequences in FASTA format. Proteins starting with methionine (M) followed by glycine (G) were selected for further analyses. Then, redundancy was removed using CD-HIT (http://weizhongli-lab.org/cd-hit/). A total of 3,229 non-redundant plant virus proteins were used to predict the N-terminal myristoylation and cTP. For *Ralstonia solanacearum* GMI1000, a total of 5,002 protein sequences were downloaded from UniProt (https://www.uniprot.org/proteomes/UP000001436). For Arabidopsis, the annotated proteins (35,386) were downloaded from http://www.arabidopsis.org/download_files/Proteins/TAIR10_protein_lists/TAIR10_pep_20101214. Proteins starting with MG (3,278) were selected for further analyses. For tomato (*Solanum lycopersicum*), the annotated tomato proteins (35,768) were downloaded from ftp://ftp.solgenomics.net/tomato_genome/annotation/ITAG3.2_release/ITAG3.2_proteins.fasta. Proteins starting with MG (3,456) were selected for further analyses. For rice (*Oryza sativa* subsp. *japonica*), the annotated proteins (66,338) were downloaded from http://rice.plantbiology.msu.edu/pub/data/Eukaryotic_Projects/o_sativa/annotation_dbs/pseudomolecules/version_7.0/all.dir/all.pep. Proteins starting with MG (5,954) were selected for further analyses. For *Chlamydomonas reinhardtii*, a total of 18,829 protein sequences were downloaded from UniProt (https://www.uniprot.org/proteomes/UP000006906). For *Marchantia polymorpha*, a total of 40,580 protein sequences were downloaded from UniProt (https://www.uniprot.org/uniprot/?query=Marchantia%20polymorpha). For *Physcomitrella patens*, a total of 30,858 protein sequences were downloaded from UniProt (https://www.uniprot.org/proteomes/UP000006727).

**Accession Numbers**

Sequence information of the genes studied in this article can be found in Supplemental table 8. Accession numbers were obtained from the Arabidopsis TAIR database (https://www.arabidopsis.org), National Center for Biotechnology Information (https://www.ncbi.nlm.nih.gov), and the Sol Genomics Network website (https://solgenomics.net/).

### SUPPLEMENTAL FIGURES

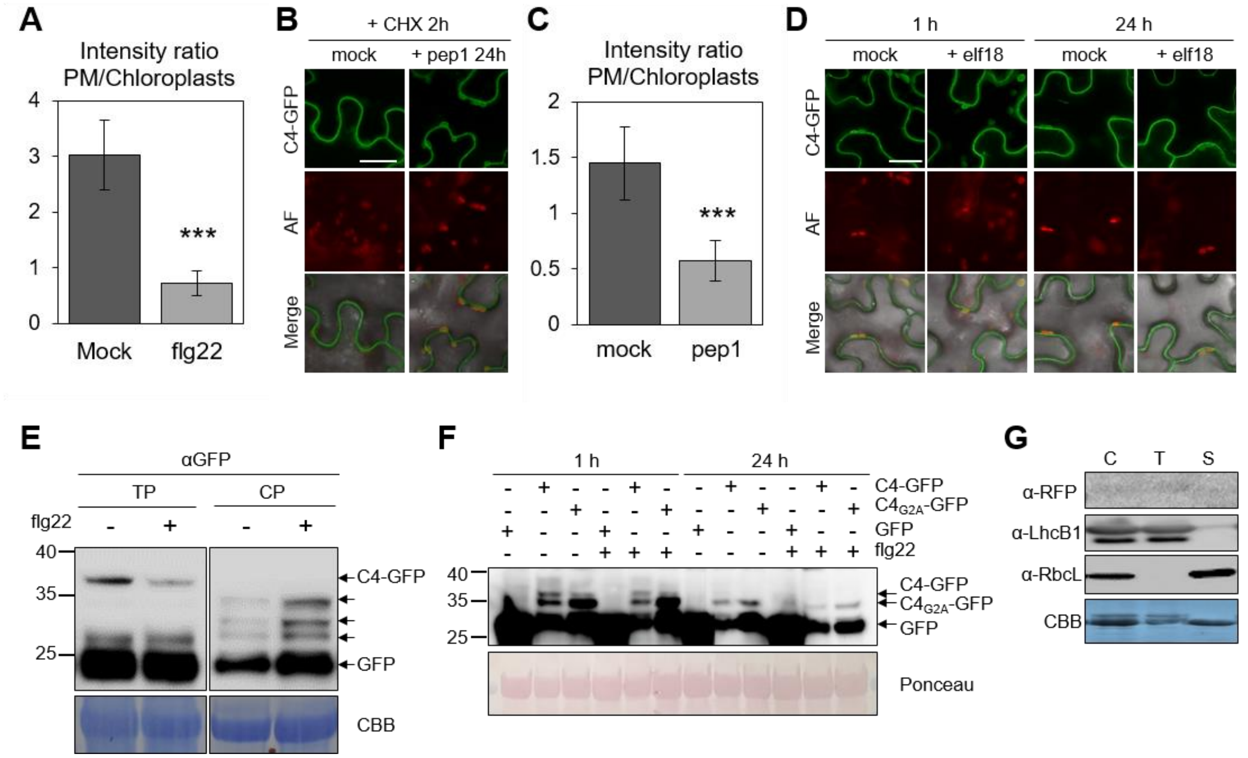


**Supplemental figure 1. C4 shifts its localization from the plasma membrane to the chloroplast following activation of PTI.** (**A**) C4-GFP PM/chloroplasts intensity ratio upon flg22 or mock treatment (Figure 1E) was quantified using ImageJ software. Bars represent SE of n = 6. Asterisks indicate a statistically significant difference (****P* value = 0.0004) according to a two-tailed comparisons *t*-test. (**B**) C4 re-localizes from PM to chloroplasts in response to treatment with the damage-associated molecular pattern Pep1 upon transient expression in *N. benthamiana* leaves. Localization of the wild-type C4 version fused to GFP was compared following treatment with 1 µM Pep1 or mock treatment (24 h post-treatment) in the presence of CHX (50 µg/ml, 2 h). Scale bar = 25 µm. AF: Autofluorescence. (**C**) C4-GFP PM/chloroplasts intensity ratio from C4 re-localization upon Pep1 or mock treatment (Supplemental figure 1B) was quantified using ImageJ software. Bars represent SE of n = 6. Asterisks indicate a statistically significant difference (****P* value = 0.0002) according to a two-tailed comparisons *t*-test. (**D**) C4 does not re-localize from PM to chloroplasts in response to elf18 peptide in *N. benthamiana* leaves. Localization of the wild-type C4 version fused to GFP was compared following treatment with 1 µM elf18 or mock treatment (1 h and 24 h post-treatment). Scale bar = 25 µm. AF: Autofluorescence. (**E**) Western blot of total and chloroplast-localized C4-GFP transiently expressed in *N. benthamiana* leaves with or without flg22 treatment. Leaves were incubated 2h with CHX (50 µg/ml) prior to PAMP treatment (flg22 1 µM, 12 h). TP: total protein. CP: chloroplast protein from an isolated chloroplast fraction. CBB: Comassie brilliant blue. Arrows indicate C4 mature chloroplast forms. (**F**) Western blot of C4-GFP and $C4_{G2A}$-GFP transiently expressed in *N.*

*benthamiana* leaves with or without flg22 treatment. Leaves were incubated 2h with CHX (50 µg/ml) prior to PAMP treatment (flg22 1 µM, 1 h and 24 h). (**G**) Chloroplast fractionation of a *35S:RFP* Arabidopsis transgenic line (as a negative control to Figure 1E). Isolated chloroplasts from three-week-old Arabidopsis transgenic lines expressing free RFP were separated into membrane and stroma fractions. (C: total chloroplast; T: thylakoid; S: stroma; LhcB1: light harvesting complex protein B1 (25 kDa), a thylakoid membrane protein; RbcL: rubisco large subunit (52.7 kDa), a stromal protein). CBB: Comassie brilliant blue.

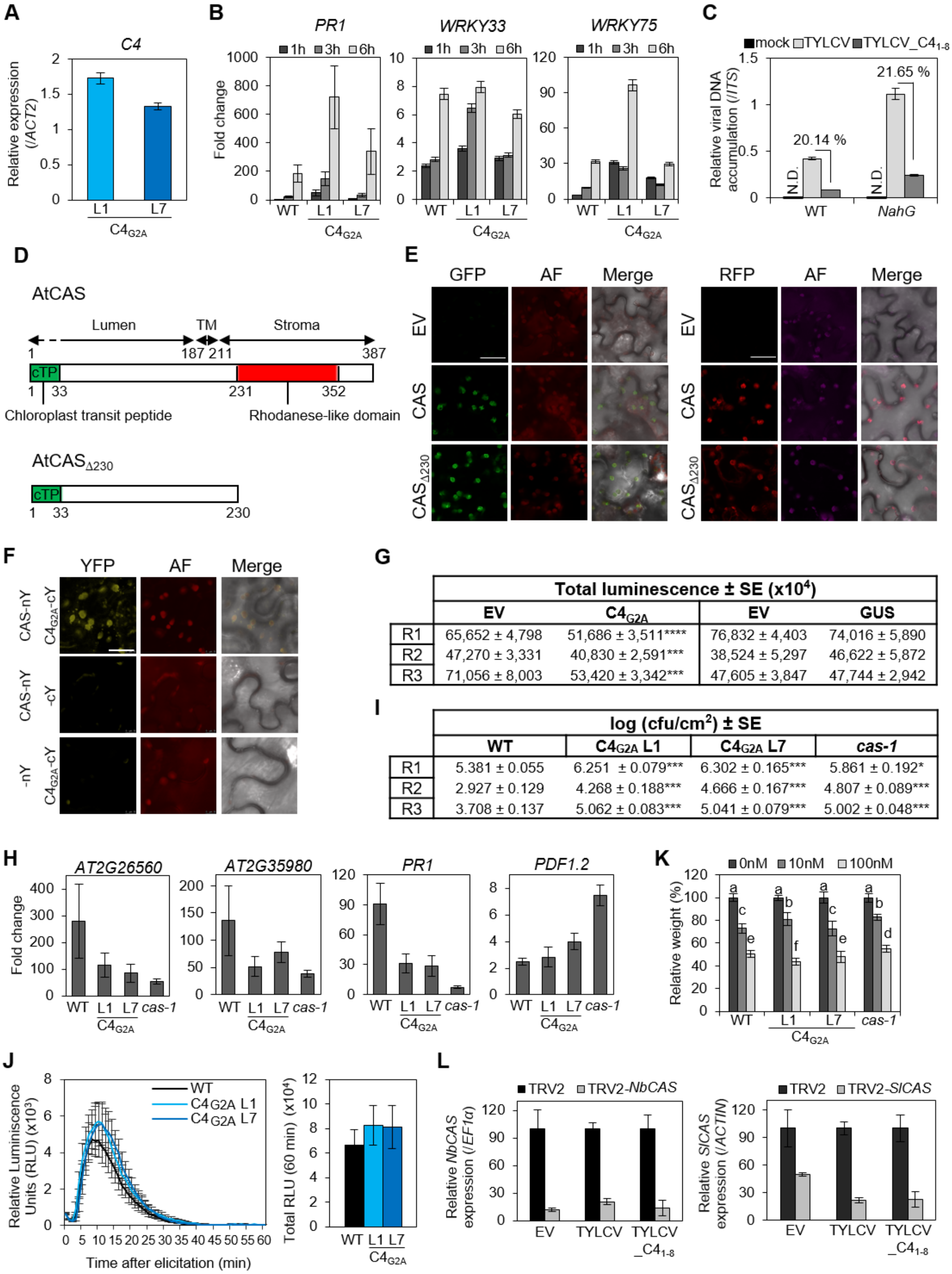


**G**

| | Total luminescence ± SE (x10$^4$) | | | |
|---|---|---|---|---|
| | EV | C4$_{G2A}$ | EV | GUS |
| R1 | 65,652 ± 4,798 | 51,686 ± 3,511**** | 76,832 ± 4,403 | 74,016 ± 5,890 |
| R2 | 47,270 ± 3,331 | 40,830 ± 2,591*** | 38,524 ± 5,297 | 46,622 ± 5,872 |
| R3 | 71,056 ± 8,003 | 53,420 ± 3,342*** | 47,605 ± 3,847 | 47,744 ± 2,942 |

**I**

| | log (cfu/cm$^2$) ± SE | | | |
|---|---|---|---|---|
| | WT | C4$_{G2A}$ L1 | C4$_{G2A}$ L7 | *cas-1* |
| R1 | 5.381 ± 0.055 | 6.251 ± 0.079*** | 6.302 ± 0.165*** | 5.861 ± 0.192* |
| R2 | 2.927 ± 0.129 | 4.268 ± 0.188*** | 4.666 ± 0.167*** | 4.807 ± 0.089*** |
| R3 | 3.708 ± 0.137 | 5.062 ± 0.083*** | 5.041 ± 0.079*** | 5.002 ± 0.048*** |

**Supplemental figure 2. C4 interacts with CAS in the chloroplast and suppresses SA-dependent defences.** (**A**) Expression of the C4 transgene detected by qRT-PCR in 35S:C4$_{G2A}$ transgenic Arabidopsis plants. Data are mean ± SE of n= 6. (**B**) Transgenic Arabidopsis plants expressing C4$_{G2A}$ respond normally to exogenous SA treatments. SA-induced gene expression was analysed at the indicated time points by qRT–PCR. *ACT2* was used as an internal standard. Data are mean ± SE of three experiments. (**C**) Viral accumulation in SA-depleted transgenic tomato lines. Three-week-old transgenic *NahG* tomato plants were infected with TYLCV wild type or a C4 null mutant (TYLCV_C4$_{1\text{-}8}$). The relative viral DNA accumulation in plants was determined by qPCR of total DNA extracted from the six youngest apical leaves at 21 dpi. Values represent the average of 8 plants. Error bars represent SE. ND: not detectable. "%" indicates the percentage of TYLCV_C4$_{1\text{-}8}$ accumulation compared to TYLCV (100%). Experiments were repeated at least three times; results from one representative experiment are shown. (**D**) Schematic representation of CAS and CAS$_{\Delta 230}$. (**E**) Subcellular localization of CAS and CAS$_{\Delta 230}$. GFP and RFP tags were fused to the Ct of the proteins. AF: autofluorescence. EV: empty vector. Scale bar = 25 µm. (**F**) C4$_{G2A}$ interacts with CAS by biomolecular fluorescence complementation (BiFC) assay. YFP fluorescence was observed in the chloroplasts upon transient coexpression in *N. benthamiana* leaves. AF: autofluorescence. Scale bar = 25 µm. (**G**) Total luminescence from calcium burst assays after elicitation with 1 µM flg22 in *N. benthamiana* leaves in three independent replicates (from Figure 2H). Asterisks indicate a statistically significant difference (***$P$ < 0.001, ****$P$ < 0.0001) according to a two-tailed comparisons *t*-test. EV: empty vector. (**H**) Chloroplast-localized C4 triggers transcriptional changes consistent with loss of function of CAS. qRT-PCR validation of RNA-seq data. *ACT2* was used as the normalizer. Data are mean ± SE of four independent biological replicates. (**I**) *Pto*DC3000 growth in transgenic Arabidopsis plants expressing C4$_{G2A}$ o the *cas-1* mutant in three independent replicates (from Figure 2K). Asterisks indicate a statistically significant difference (*$P$ < 0.05, ***$P$ < 0.001) according to a one-way ANOVA with post-hoc Dunnett's multiple comparisons test. (**J**) and (**K**) Transgenic Arabidopsis plants expressing C4$_{G2A}$ are not affected in PAMP perception. (**J**) ROS-burst as relative luminescence units (RLU) during 60 min and as total RLU after flg22 treatment. Error bars indicate SE (n = 8) ($P$ = 0.7276, considered not significant, one-way ANOVA). (**K**) Seedling growth inhibition (SGI) assay after flg22 treatment. Error bars indicate SE (n = 12). Lowercase letters indicate statistically significant differences between mean values ($P$ <0.0001), according to a one-way ANOVA with post-hoc Student's test. (**L**) Virus-induced gene silencing (VIGS) of *CAS* in *N. benthamiana* and tomato. Gene silencing was quantified by qRT-PCR. RNA was extracted from the three youngest apical leaves at 21 dpi. Values represent the average of six plants. *NbEF1α* and *SlActin* were used as internal standard for *N. benthamiana* and tomato, respectively. Error bars represent SE. Experiments were repeated at least three times with similar results; results from one experiment are shown. EV: empty vector.

**A**

| Family | Subfamily | Genus | Type of genome | Number of proteins containing both targeting signals |
|---|---|---|---|---|
| *Geminiviridae* | | Begomovirus | dsDNA | 390 |
| | | Capulavirus | dsDNA | 1 |
| | | Curtovirus | dsDNA | 5 |
| | | Topocuvirus | dsDNA | 1 |
| | | Turncurtovirus | dsDNA | 1 |
| *Bromoviridae* | | Cucumovirus | (+)ssRNA | 1 |
| *Secoviridae* | Comovirinae | Nepovirus | (+)ssRNA | 1 |
| *Tombusviridae* | Calvusvirinae | Umbravirus | (+)ssRNA | 1 |

**B**

| Genus | Species | Isolate | Accession number | Protein | Protein ID | AA | myr | cTP (AA) |
|---|---|---|---|---|---|---|---|---|
| Curtovirus | *Beet curly top virus (BCTV)* | California [Logan] | M24597 | C4 | ADD82466 | 85 | yes | 69 |
| Begomovirus | *East African cassava mosaic virus (EACMV)* | EACMV-KE2[K48] | AJ717542 | AC4 | CAJ78099 | 77 | yes | 49 |
| Cucumovirus | *Cucumber mosaic virus (CMV)* | DN2-1 | KT302169 | CP | ALT66618 | 218 | yes | 25 |
| Nepovirus | *Grapevine fanleaf virus (GFLV)* | SACH44 | KC900164 | P3 | AGT42202 | 338 | yes | 42 |

**C**

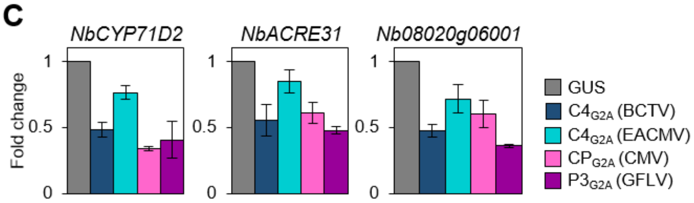


**D**

| | Total luminescence ± SE (x10$^4$) | | | | | | | |
|---|---|---|---|---|---|---|---|---|
| | EV | $C4_{G2A}$ (BCTV) | EV | $C4_{G2A}$ (EACMV) | EV | $CP_{G2A}$ (CMV) | EV | $P3_{G2A}$ (GFLV) |
| R1 | 19.672 ± 2.938 | 10.336 ± 1.946**** | 32.244 ± 3.464 | 22.606 ± 2.543**** | 17.464 ± 2.753 | 11.490 ± 7.297** | 14.208 ± 2.869 | 9.300 ± 0.832** |
| R2 | 17.314 ± 4.921 | 10.458 ± 1.469** | 18.804 ± 3.982 | 13.020 ± 1.632*** | 25.840 ± 9.995 | 9.856 ± 1.777**** | 40.301 ± 3.777 | 35.415 ± 4.368** |
| R3 | 42.844 ± 6.210 | 42.692 ± 5.539 | 11.242 ± 1.639 | 5.422 ± 0.972**** | 18.528 ± 2.561 | 15.648 ± 1.944* | 20.840 ± 3.569 | 17.982 ± 2.967 |

| | Total luminescence ± SE (x10$^4$) | |
|---|---|---|
| | EV | GUS |
| R1 | 27.370 ± 3.730 | 27.190 ± 2.906 |
| R2 | 55.724 ± 19.385 | 46.944 ± 14.336 |
| R3 | 19.132 ± 4.089 | 25.374 ± 4.499 |

**E**

| | log (cfu/cm$^2$) ± SE | | | | | | | |
|---|---|---|---|---|---|---|---|---|
| | GUS | $C4_{G2A}$ (BCTV) | GUS | $C4_{G2A}$ (EACMV) | GUS | $CP_{G2A}$ (CMV) | GUS | $P3_{G2A}$ (GFLV) |
| R1 | 6,918 ± 0,129 | 7,943 ± 0,048*** | 6,979 ± 0,052 | 8,043 ± 0,028*** | 7,124 ± 0,033 | 8,036 ± 0,028*** | 7,156 ± 0,017 | 7,129 ± 0,021 |
| R2 | 7,059 ± 0,102 | 7,952 ± 0,06*** | 7,037 ± 0,044 | 8,052 ± 0,072*** | 6,747 ± 0,149 | 7,621 ± 0,151*** | 6,908 ± 0,098 | 6,704 ± 0,072 |
| R3 | 7,088 ± 0,034 | 8,061 ± 0,034*** | 7,127 ± 0,051 | 8,005 ± 0,088*** | 7,007 ± 0,033 | 7,965 ± 0,055*** | 7,001 ± 0,019 | 6,989 ± 0,025 |

**F**

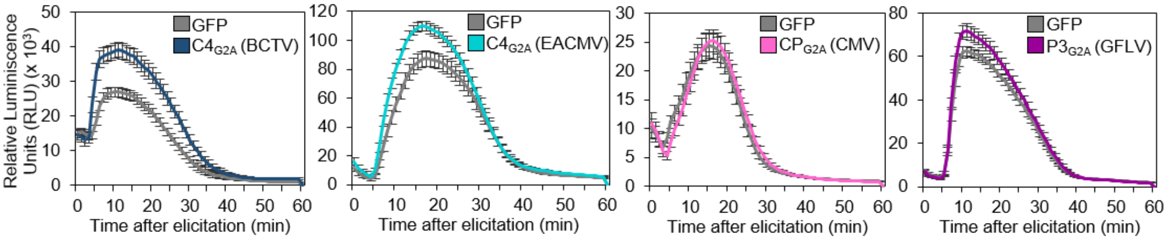

**Supplemental figure 3. Proteins encoded by plant viruses and containing overlapping N-myristoylation motifs and cTPs suppress defence responses from the chloroplast.** (**A**) Plant virus species encoding proteins with overlapping N-myristoylation motifs and cTPs. (**B**) Selected protein sequences from phylogenetically unrelated viral proteins with overlapping N-myristoylation motif and cTP. (**C**) Viral proteins with the overlapping N-myristoylation motif and cTP suppress expression of SA-related genes upon activation of plant immunity in *N. benthamiana* leaves. PAMP treatment (1 µM flg22) was performed 48 h after transient transformation. Leaf discs from three plants were collected separately nine hours after treatment. Gene expression was analysed by qRT–PCR. *NbEF1α* was used as an internal standard. Values represent the average of four independent experiments with three plants used in each replicate. Data are mean ± SE of three independent experiments. (**D**) Total luminescence from calcium burst after elicitation with 1 µM flg22 in *N. benthamiana* leaves in three independent replicates (from Figure 3E). Asterisks indicate a statistically significant difference (*$P < 0.05$, **$P < 0.01$, ***$P < 0.001$, ****$P < 0.0001$) according to a two-tailed comparisons *t*-test. EV: empty vector. (**E**) *Pto*DC3000 growth in *N. benthamiana* leaves transiently expressing selected chloroplast-localized viral proteins in three independent replicates (from Figure 3F). Asterisks indicate a statistically significant difference (***$P < 0{,}001$) according to a two-tailed comparisons *t*-test. (**F**) Kinetics of ROS production as relative luminescence units (RLU) during 60 min in response to the elicitor flg22 in *N. benthamiana* leaves transiently expressing chloroplast-localized proteins (from Figure 3G). Error bars indicate SE (n = 24). Experiments were repeated at least three times with similar results; results from one experiment are shown.

**A**

| *Ralstonia solanacearum* GMI1000 | | | | | |
|---|---|---|---|---|---|
| **Transcript_id** | **Gene name** | **Description** | **AA** | **myr** | **cTP (AA)** |
| Q8XRE0 | *RSp0914* | Irr-gala family type III effector protein (GALA1) | 661 | yes | 39 |
| Q8XTS6 | *RSp0028* | Irr-gala family type III effector protein (GALA3) | 603 | yes | 58 |

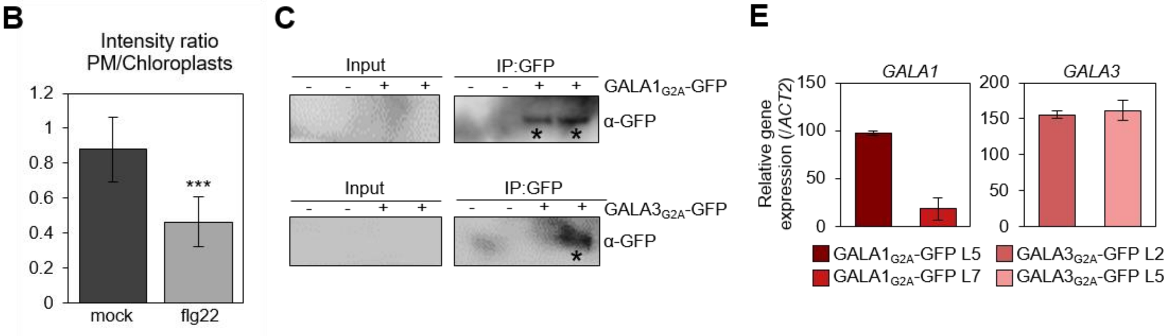


**D**

| | Total luminescence ± SE (x10$^4$) | | | | | |
|---|---|---|---|---|---|---|
| | **EV** | **GALA1$_{G2A}$** | **EV** | **GALA3$_{G2A}$** | **EV** | **GUS** |
| R1 | 32.840 ± 4.518 | 20.434 ± 1.254**** | 30.102 ± 5.778 | 16.640 ± 2.014*** | 55.724 ± 19.385 | 46.944 ± 14.336 |
| R2 | 38.644 ± 7.959 | 18.472 ± 1.159**** | 23.756 ± 2.284 | 17.838 ± 1.646*** | 21.776 ± 4.414 | 34.418 ± 9.276 |
| R3 | 112.576 ± 23.158 | 49.260 ± 10.499**** | 103.916 ± 30.522 | 37.532 ± 7.888**** | 69.060 ± 8.960 | 77.337 ± 8.762 |

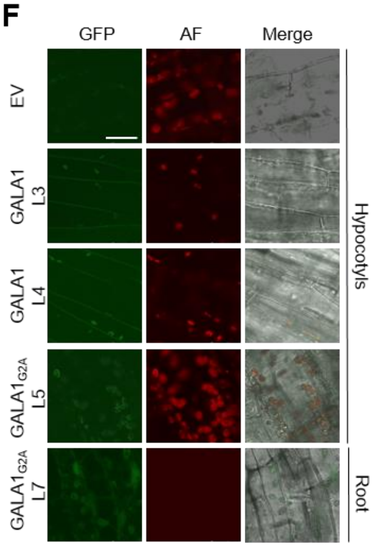


**G**

| | log (cfu/FW (g)) ± SE | | | | |
|---|---|---|---|---|---|
| | **EV** | **GALA1$_{G2A}$ L5** | **GALA1$_{G2A}$ L7** | **GALA3$_{G2A}$ L2** | **GALA3$_{G2A}$ L5** |
| R1 | 8.623 ± 0.152 | 9.083 ± 0.076* | 8.566 ± 0.114 | 9.031 ± 0.113* | 8.934 ± 0.094* |
| R2 | 6.892 ± 0.284 | 8.105 ± 0.126** | 8.091 ± 0.129** | 8.001 ± 0.111** | 7.592 ± 0.106** |
| R3 | 7.332 ± 0.178 | 7.601 ± 0.113** | 7.713 ± 0.118** | 7.109 ± 0.088 | 7.294 ± 0.11 |
| R4 | 6.231 ± 0.133 | 7.123 ± 0.218** | 6.359 ± 0.148 | 6.332 ± 0.151 | 6.708 ± 0.163*** |

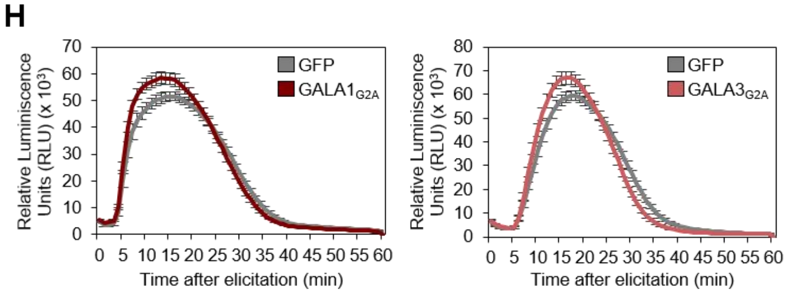


**Supplemental figure 4. Proteins encoded by the plant pathogenic bacterium *Ralstonia solanacearum* and containing overlapping N-myristoylation motifs and cTPs suppress defence responses from the chloroplast.** (**A**) Proteins encoded by *Ralstonia solanacearum* GMI1000 containing an N-myristoylation motif and a cTP. (**B**) GALA1-GFP PM/chloroplasts intensity ratio upon flg22 or mock treatment (Figure 4C) was quantified using ImageJ software. Bars represent SE of n = 6. Asterisks indicate a statistically significant difference (****P* value = 0.0003) according to a two-tailed comparisons *t*-test. (**C**) GALA1$_{G2A}$ and GALA3$_{G2A}$ accumulate in chloroplasts upon transient expression in *N. benthamiana* leaves. GALA$_{G2A}$-GFP proteins were detected in chloroplasts isolated from *N. benthamiana* leaves (2 dpi) after immunoprecipitation with GFP-Trap beads. “-” indicates samples infiltrated with empty vector. “+” indicates samples expressing $GALA1_{G2A}$ or $GALA3_{G2A}$. “*” marks the expected bands. (**D**) Total luminescence from calcium burst after elicitation with 1 µM flg22 in *N. benthamiana* leaves transiently expressing $GALA1_{G2A}$ or $GALA3_{G2A}$ in three independent replicates. Asterisks indicate a statistically significant difference (****P* < 0.001, *****P* < 0.0001) according to a two-tailed comparisons *t*-test. EV: empty vector. (**E**) Expression of *GALA1* and *GALA3* transgenes detected by qRT-PCR in 10-day-old transgenic Arabidopsis seedlings. Data are mean ± SE of n= 6. (**F**) Localization of the wild-type (GALA1) and the non-myristoylable ($GALA1_{G2A}$) versions of GALA1 fused to GFP in 3-day-old transgenic Arabidopsis lines. Scale bar = 25 µm. AF: autofluorescence. EV: empty vector. (**G**) *Pto*DC3000 growth in transgenic Arabidopsis plants expressing chloroplast-localized GALA1 or GALA3 ($GALA1_{G2A}$ and $GALA3_{G2A}$ plants) (from Figure 4G). Values represent the average of four independent replicates. Values of each replicate were calculated from four independent plants. Data are mean ± SE. Asterisks indicate a statistically significant difference (**P* < 0.05, ***P* < 0.01, ****P* < 0.001) according to a one-way ANOVA with post-hoc Dunnett's multiple comparisons test. EV: empty vector. (**H**) Kinetics of ROS production as relative luminescence units (RLU) during 60 min in response to the elicitor flg22 in *N. benthamiana* leaves transiently expressing $GALA1_{G2A}$ or $GALA3_{G2A}$ (from Figure 4H). Error bars indicate SE (n = 24). Experiments were repeated at least three times with similar results; results from one experiment are shown.

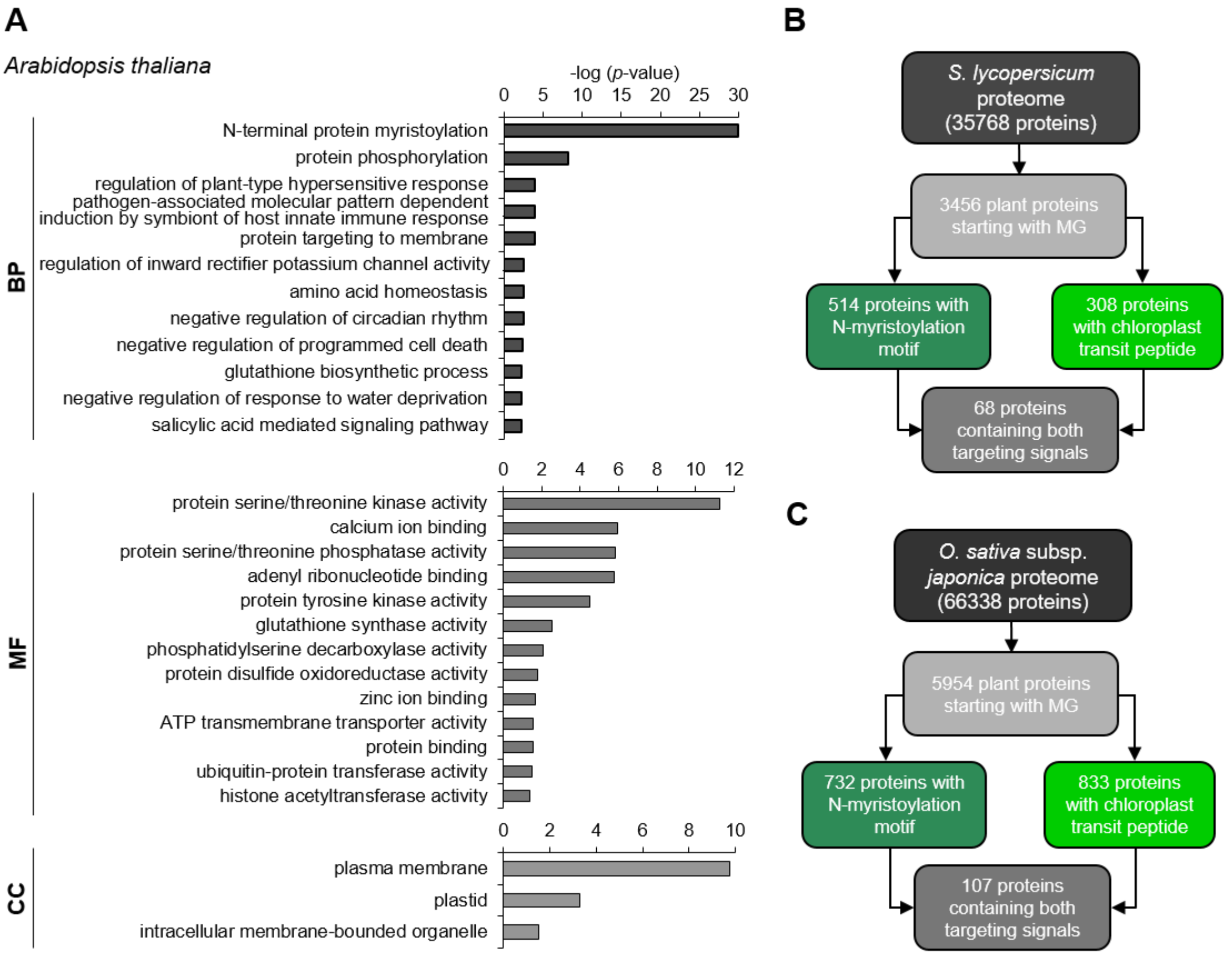

A
Arabidopsis thaliana
-log (p-value)
0 5 10 15 20 25 30
BP
N-terminal protein myristoylation
protein phosphorylation
regulation of plant-type hypersensitive response
pathogen-associated molecular pattern dependent induction by symbiont of host innate immune response
protein targeting to membrane
regulation of inward rectifier potassium channel activity
amino acid homeostasis
negative regulation of circadian rhythm
negative regulation of programmed cell death
glutathione biosynthetic process
negative regulation of response to water deprivation
salicylic acid mediated signaling pathway
0 2 4 6 8 10 12
MF
protein serine/threonine kinase activity
calcium ion binding
protein serine/threonine phosphatase activity
adenyl ribonucleotide binding
protein tyrosine kinase activity
glutathione synthase activity
phosphatidylserine decarboxylase activity
protein disulfide oxidoreductase activity
zinc ion binding
ATP transmembrane transporter activity
protein binding
ubiquitin-protein transferase activity
histone acetyltransferase activity
0 2 4 6 8 10
CC
plasma membrane
plastid
intracellular membrane-bounded organelle
B
S. lycopersicum proteome (35768 proteins)
3456 plant proteins starting with MG
514 proteins with N-myristoylation motif
308 proteins with chloroplast transit peptide
68 proteins containing both targeting signals
C
O. sativa subsp. japonica proteome (66338 proteins)
5954 plant proteins starting with MG
732 proteins with N-myristoylation motif
833 proteins with chloroplast transit peptide
107 proteins containing both targeting signals

**D**

*Solanum lycopersicum*

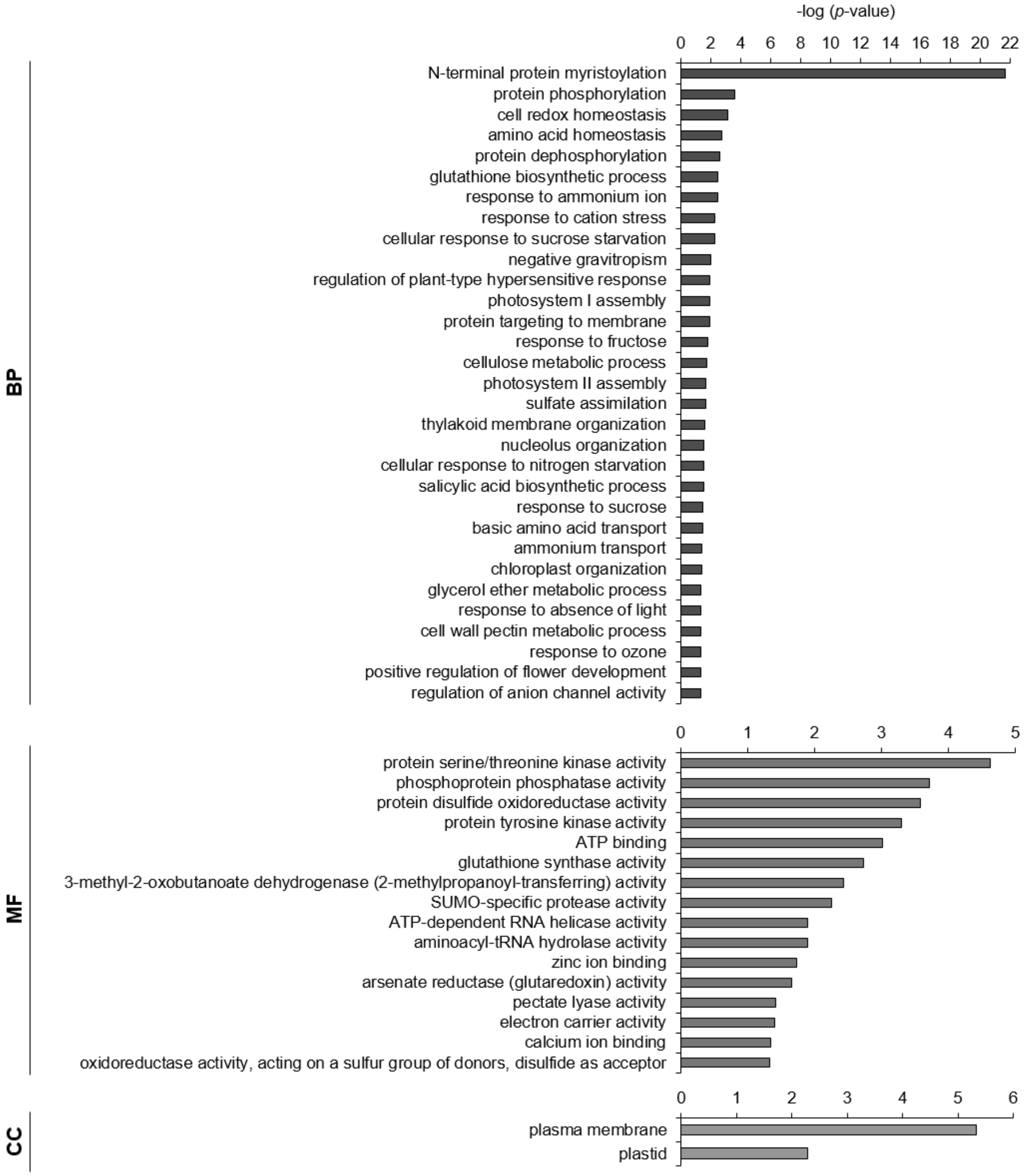

**E**

*Oryza sativa* subsp. *japonica*

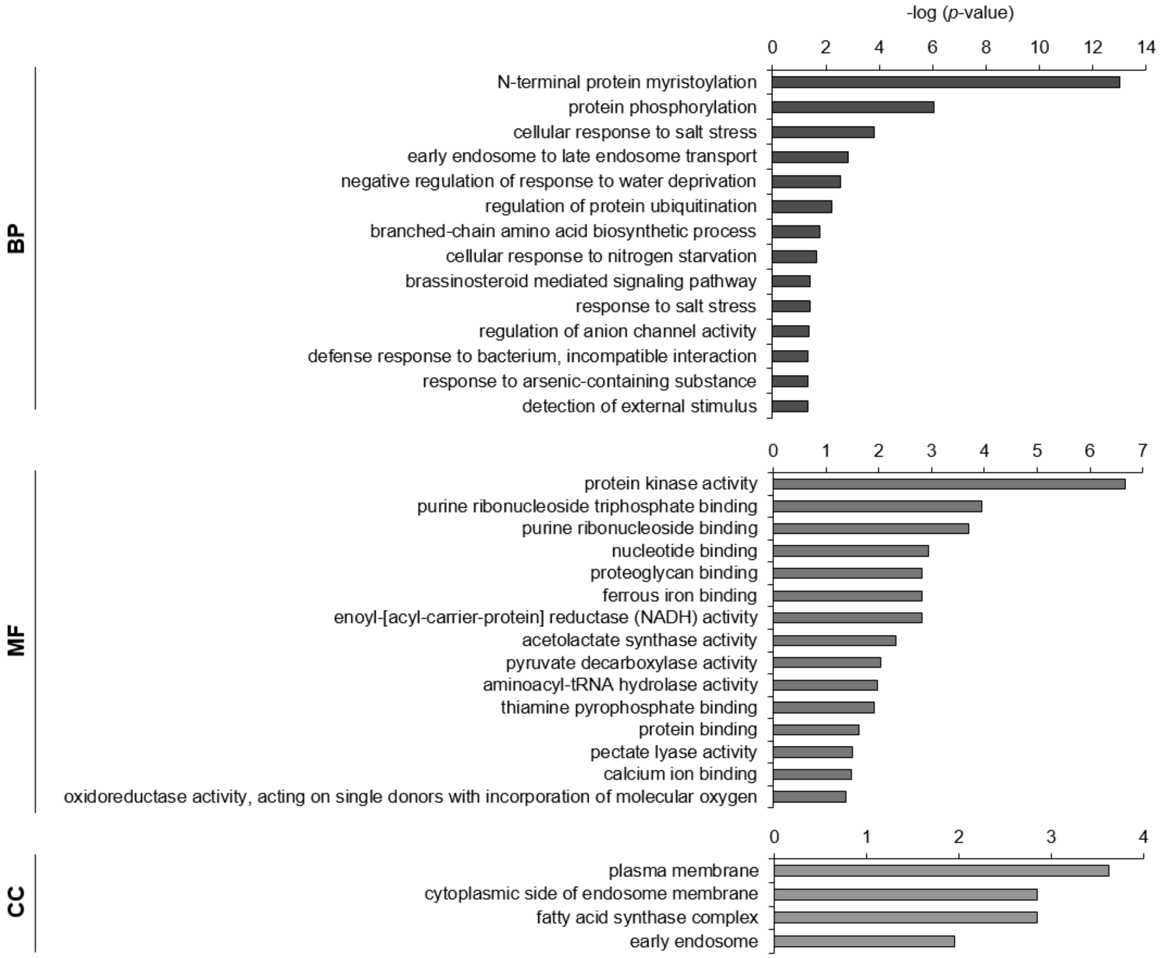

**F**

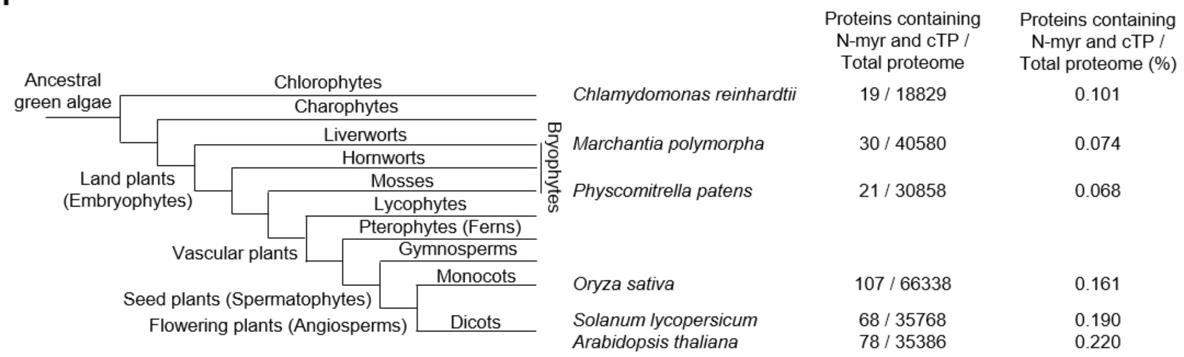


**G**

*Chlamydomonas reinhardtii*

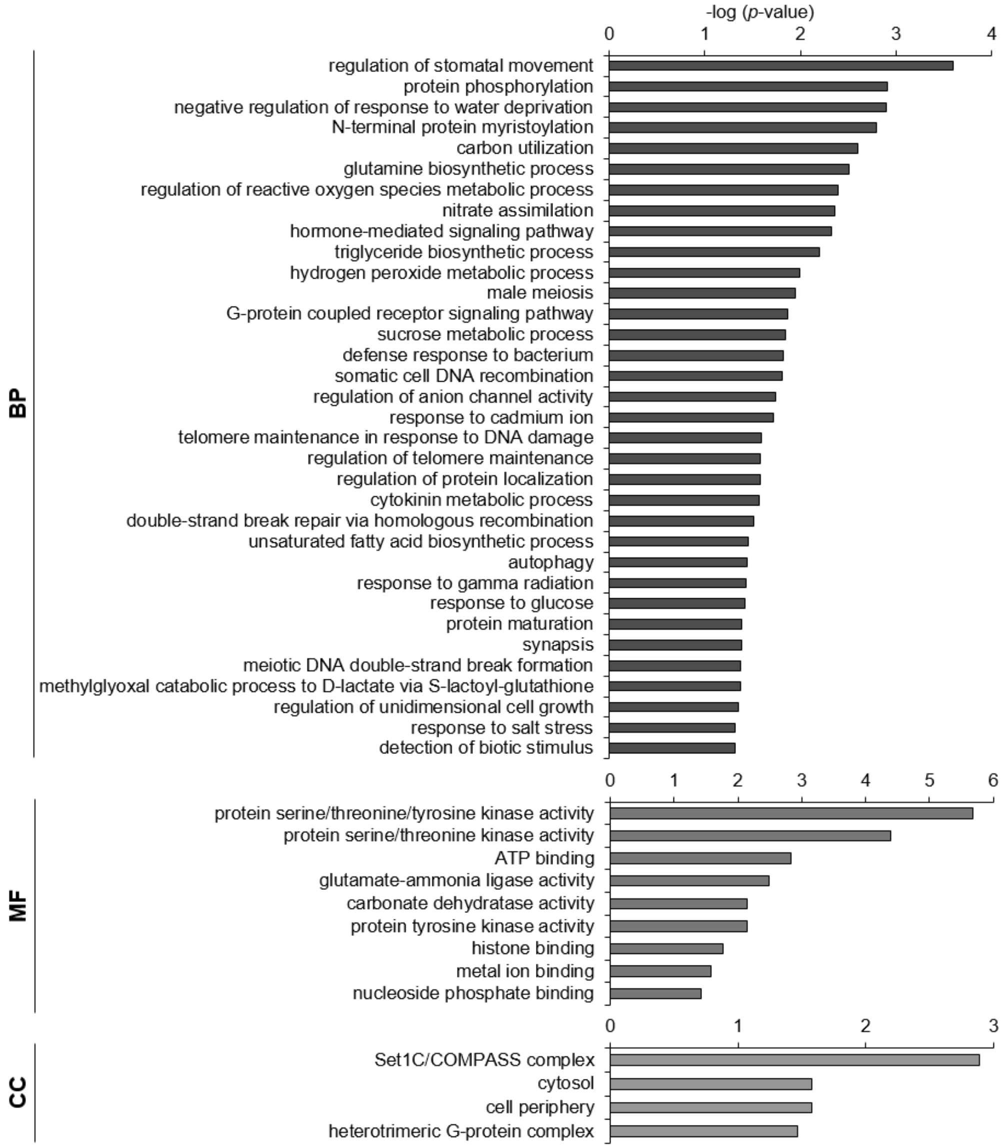

**H**

*Marchantia polymorpha*

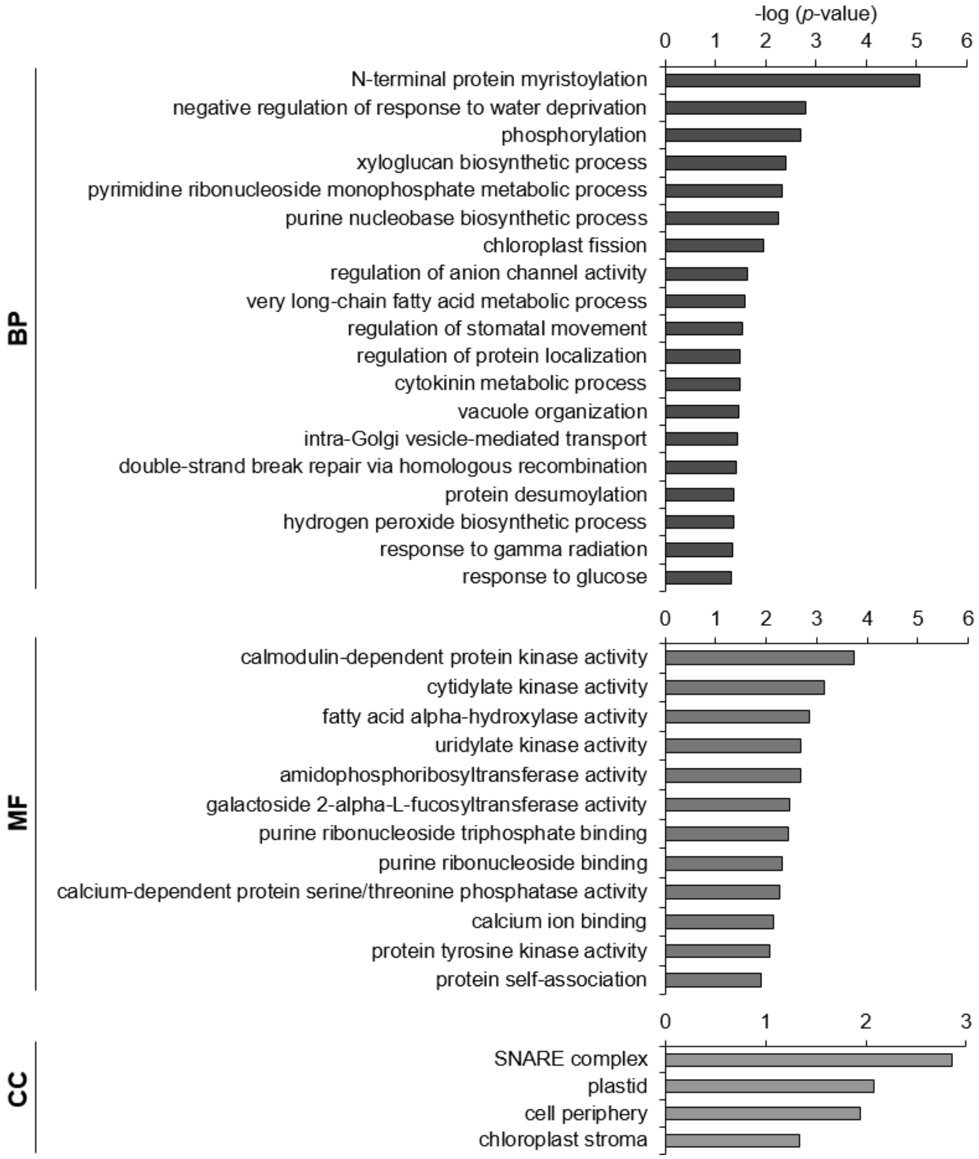

I

*Physcomitrella patens*

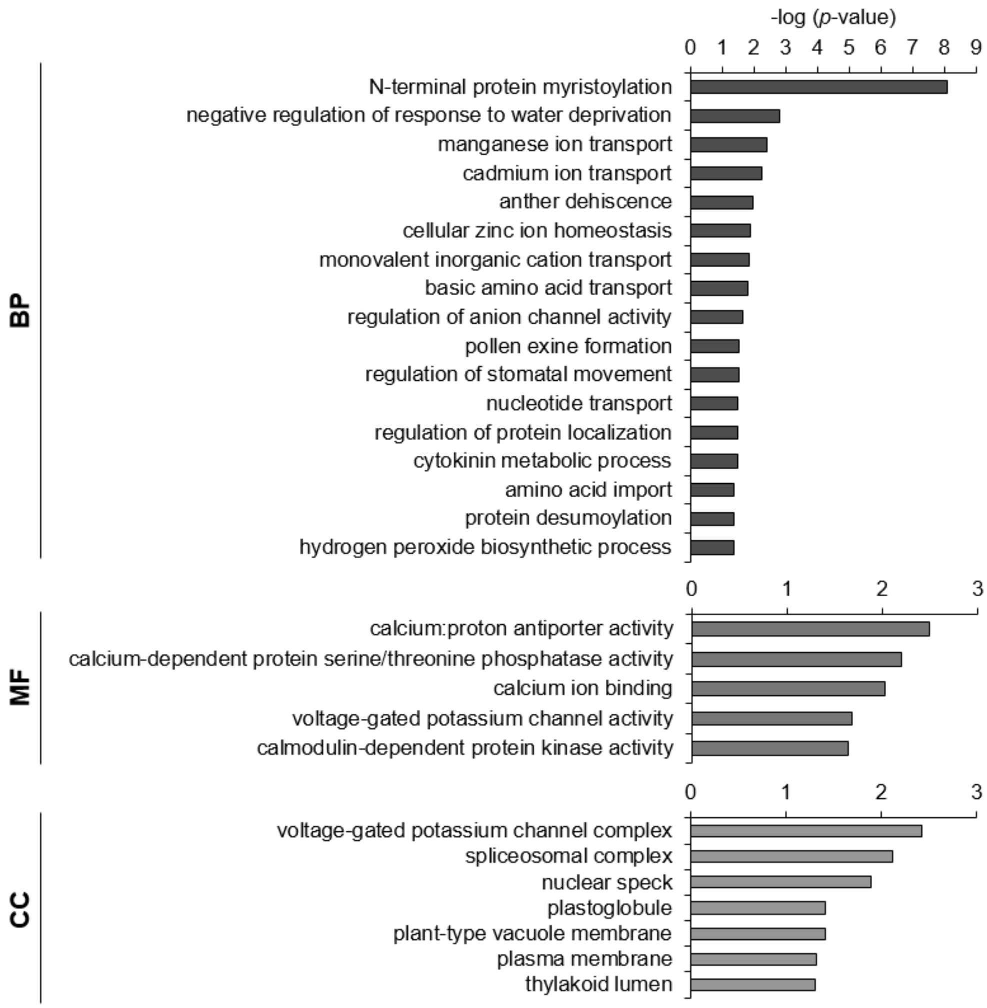

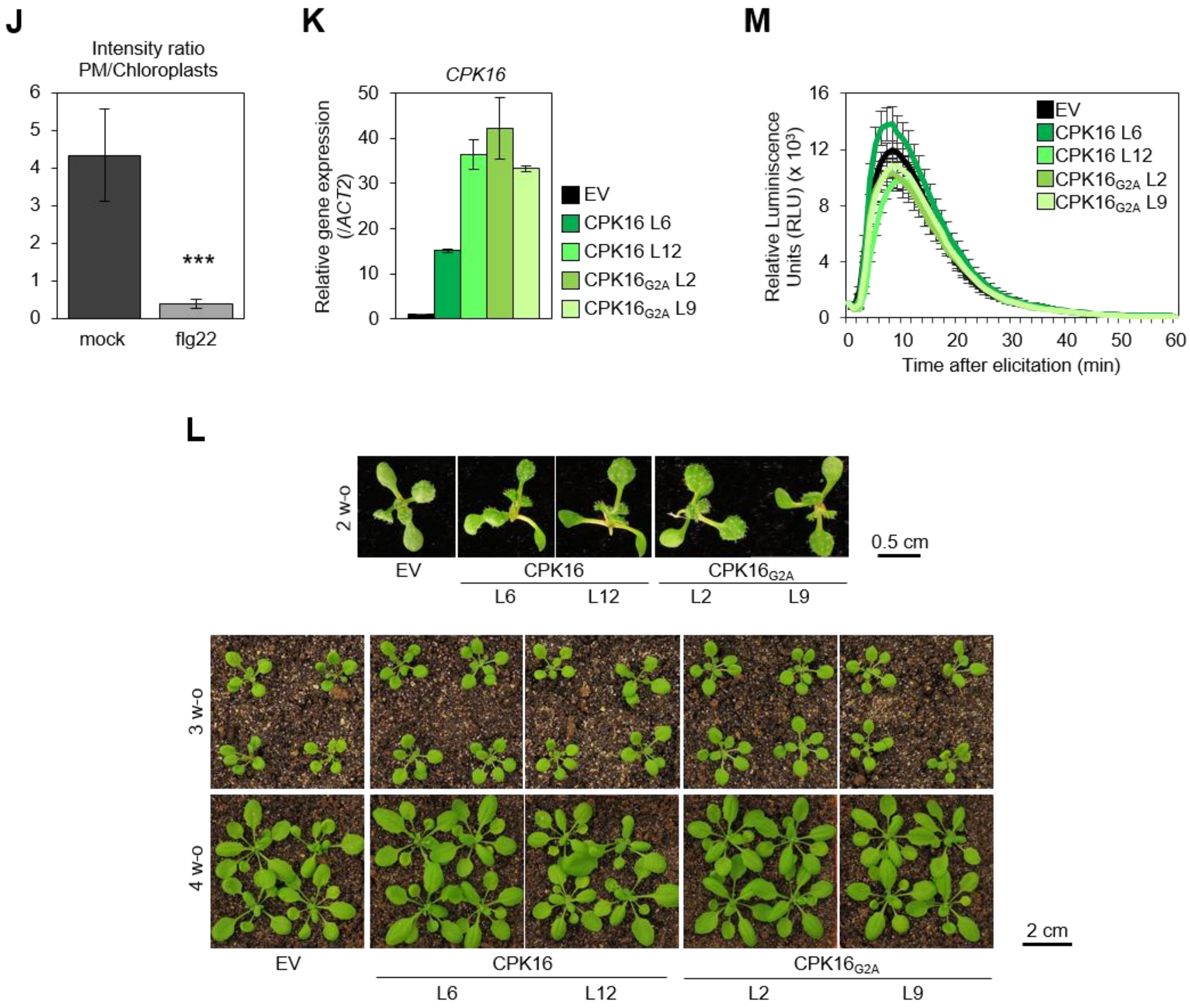


**N**

| | Total luminescence ± SE (x10$^4$) | | | | | |
|---|---|---|---|---|---|---|
| | EV | CPK16 | EV | CPK16$_{G2A}$ | EV | GUS |
| R1 | 50.172 ± 6.442 | 79.444 ± 11.089**** | 44.112 ± 8.267 | 53.006 ± 8.111 | 55.724 ± 19.385 | 46.944 ± 14.336 |
| R2 | 85.995 ± 16.614 | 125.572 ± 30.771* | 106.774 ± 14.077 | 115.980 ± 20.191 | 69.060 ± 8.961 | 77.337 ± 8.762 |
| R3 | 11.972 ± 1.707 | 19.391 ± 6.107* | 13.688 ± 1.807 | 12.630 ± 1.861 | 14.700 ± 2.277 | 12.333 ± 2.813 |

**O**

| | log (cfu/FW (g)) ± SE | | | | |
|---|---|---|---|---|---|
| | EV | CPK16 L6 | CPK16 L12 | CPK16$_{G2A}$ L2 | CPK16$_{G2A}$ L9 |
| R1 | 8.053 ± 0.070 | 8.037 ± 0.096 | 8.126 ± 0.053*** | 7.322 ± 0.058*** | 7.407 ± 0.072*** |
| R2 | 7.510 ± 0.129 | 7.575 ± 0.112 | 6.829 ± 0.060*** | 7.651 ± 0.100* | 6.789 ± 0.092*** |
| R3 | 6.990 ± 0.059 | 6.776 ± 0.008* | 6.668 ± 0.115** | 6.591 ± 0.147** | 5.691 ± 0.181*** |
| R4 | 6.669 ± 0.1512 | 6.248 ± 0.085*** | 6.291 ± 0.098*** | 6.294 ± 0.104*** | 5.983 ± 0.125*** |

**Supplemental figure 5. A core of conserved plant defence-related proteins contain overlapping N-myristoylation motifs and cTPs.** (**A**) Functional enrichment analysis of the 78 proteins in the *A. thaliana* proteome containing overlapping N-myristoylation motifs and cTPs. BP: biological process ontology; MF: molecular function ontology; CC: cellular component ontology. (**B**) Flow diagram illustrating the number of proteins encoded by tomato (*S. lycopersium*) containing an N-myristoylation motif, a chloroplast transit peptide, or both. (**C**) Flow diagram illustrating the number of proteins encoded by rice (*O. sativa*) containing an N-myristoylation motif, a chloroplast transit peptide, or both. (**D**) Plant phylogenetic tree indicating the relationship between the species analyzed, the number of proteins containing and N-myristoylation motif and a cTP in their predicted proteomes, and the prevalence of the coexistence of these two targeting signals. (**E**) Functional enrichment analysis of the 19 proteins in the *C. reinhardtii* proteome containing overlapping N-myristoylation motifs and cTPs. BP: biological process ontology; MF: molecular function ontology; CC: cellular component ontology. (**F**) Functional enrichment analysis of the 30 proteins in the *M. polymorpha* proteome containing overlapping N-myristoylation motifs and cTPs. BP: biological process ontology; MF: molecular function ontology; CC: cellular component ontology. (**G**) Functional enrichment analysis of the 21 proteins in the *P. patens* proteome containing overlapping N-myristoylation motifs and cTPs. BP: biological process ontology; MF: molecular function ontology; CC: cellular component ontology. (**H**) CPK16-GFP PM/chloroplasts intensity ratio upon flg22 or mock treatment (Figure 5F) was quantified using ImageJ software. Bars represent SE of n = 6. Asterisks indicate a statistically significant difference (****P* value = 0.0002) according to a two-tailed comparisons *t*-test. (**I**) Expression of *CPK16* detected by qRT-PCR in four-week-old plants over-expressing CPK16 or $CPK16_{G2A}$. Data are mean ± SE of n= 6. EV: empty vector. (**J**) Developmental phenotype of two-, three-, and four-week-old *35S:CPK16* and *35S:CPK16*$_{G2A}$ transgenic Arabidopsis lines. EV: empty vector. (**K**) Kinetics of ROS production as relative luminescence units (RLU) during 60 min in response to the elicitor flg22 in four-week-old Arabidopsis transgenic lines over-expressing CPK16 (from Figure 5G). Error bars indicate SE (n = 16) ($P = 0.666$, considered not significant, according to a one-way ANOVA). Experiments were repeated at least three times with similar results; results from one experiment are shown. EV: empty vector. (**M**) Total luminescence from calcium burst after elicitation with 1 µM flg22 in *N. benthamiana* leaves transiently overexpressing CPK16 or $CPK16_{G2A}$ in three independent replicates (from Figure 5I). Values represent the average of 6 plants. Asterisks indicate a statistically significant difference (*$P < 0.001$, ****$P < 0.0001$) according to a two-tailed comparisons *t*-test. EV: empty vector. (**L**) *Pto*DC3000 growth in transgenic Arabidopsis plants overexpressing CPK16 or $CPK16_{G2A}$ in four independent replicates (from Figure 4J). Values of each replicate were calculated from three different leaves of four independent plants. Data are mean ± SE. Asterisks indicate a statistically significant difference (*$P < 0.05$, **$P < 0.01$, ***$P < 0.001$) according to a one-way ANOVA with post-hoc Dunnett's multiple comparisons test. EV: empty vector.

### SUPPLEMENTAL TABLES

**Supplemental table 1.** Identity of the 78 proteins containing both an N-myristoylation motif and a cTP from the Arabidopsis predicted proteome.

| ID | Description |
|---|---|
| AT1G07460 | Concanavalin A-like lectin family protein |
| AT1G07570 | Protein kinase superfamily protein |
| AT1G07850 | Protein of unknown function (DUF604) |
| AT1G08860 | Calcium-dependent phospholipid-binding Copine family protein |
| AT1G09600 | Protein kinase superfamily protein |
| AT1G10410 | Protein of unknown function (DUF1336) |
| AT1G13970 | Protein of unknown function (DUF1336) |
| AT1G14370 | Protein kinase 2A |
| AT1G18290 | Unknown protein |
| AT1G18680 | HNH endonuclease domain-containing protein |
| AT1G21722 | Unknown protein |
| AT1G26360 | Methyl esterase 13 |
| AT1G26970 | Protein kinase superfamily protein |
| AT1G32190 | Alpha/beta-Hydrolases superfamily protein |
| AT1G32760 | Glutaredoxin family protein |
| AT1G33880 | Avirulence induced gene (AIG1) family protein |
| AT1G59650 | Protein of unknown function (DUF1336) |
| AT1G65050 | TRAF-like superfamily protein |
| AT1G65510 | Unknown protein |
| AT1G67800 | Copine (Calcium-dependent phospholipid-binding protein) family |
| AT1G69240 | Methyl esterase 15 |
| AT1G69650 | TRAF-like family protein |
| AT1G74380 | Xyloglucan xylosyltransferase 5 |
| AT2G02800 | Protein kinase 2B |
| AT2G17890 | Calcium-dependent protein kinase 16 |
| AT2G26190 | Calmodulin-binding family protein |
| AT2G28930 | Protein kinase 1B |
| AT2G31500 | Calcium-dependent protein kinase 24 |
| AT2G33860 | Transcriptional factor B3 family protein / auxin-responsive factor AUX/IAA-related |
| AT2G39660 | Botrytis-induced kinase1 |
| AT2G41330 | Glutaredoxin family protein |
| AT2G44230 | Plant protein of unknown function (DUF946) |
| AT2G46830 | Circadian clock associated 1 |
| AT2G46990 | Indole-3-acetic acid inducible 20 |
| AT3G01085 | Protein kinase superfamily protein |
| AT3G01860 | Unknown protein |
| AT3G02750 | Protein phosphatase 2C family protein |
| AT3G05050 | Protein kinase superfamily protein |

| | |
|---|---|
| AT3G09770 | RING/U-box superfamily protein |
| AT3G13800 | Metallo-hydrolase/oxidoreductase superfamily protein |
| AT3G19100 | Protein kinase superfamily protein |
| AT3G19460 | Reticulon family protein |
| AT3G20410 | Calmodulin-domain protein kinase 9 |
| AT3G27210 | Unknown protein |
| AT3G29770 | Methyl esterase 11 |
| AT3G50530 | CDPK-related kinase |
| AT3G54290 | Unknown protein |
| AT3G55450 | PBS1-like 1 |
| AT3G57070 | Glutaredoxin family protein |
| AT3G58210 | TRAF-like family protein |
| AT3G62100 | Indole-3-acetic acid inducible 30 |
| AT4G01210 | Glycosyl transferase family 1 protein |
| AT4G03415 | Protein phosphatase 2C family protein |
| AT4G23650 | Calcium-dependent protein kinase 6 |
| AT4G25970 | Phosphatidylserine decarboxylase 3 |
| AT4G31750 | HOPW1-1-interacting 2 |
| AT4G35230 | BR-signaling kinase 1 |
| AT4G35600 | Protein kinase superfamily protein |
| AT5G01500 | Thylakoid ATP/ADP carrier |
| AT5G01540 | Lectin receptor kinase a4 |
| AT5G02290 | Protein kinase superfamily protein |
| AT5G05400 | LRR and NB-ARC domains-containing disease resistance protein |
| AT5G06260 | TLD-domain containing nucleolar protein |
| AT5G09740 | Histone acetyltransferase of the MYST family 2 |
| AT5G14420 | RING domain ligase2 |
| AT5G15790 | RING/U-box superfamily protein |
| AT5G17350 | Unknown protein |
| AT5G20640 | Protein of unknown function (DUF567) |
| AT5G21170 | 5'-AMP-activated protein kinase beta-2 subunit protein |
| AT5G27380 | Glutathione synthetase 2 |
| AT5G36250 | Protein phosphatase 2C family protein |
| AT5G36710 | Unknown protein |
| AT5G36800 | Unknown protein |
| AT5G40880 | WD-40 repeat family protein / zfwd3 protein (ZFWD3) |
| AT5G45290 | RING/U-box superfamily protein |
| AT5G49665 | Zinc finger (C3HC4-type RING finger) family protein |
| AT5G56840 | Myb-like transcription factor family protein |
| AT5G66210 | Calcium-dependent protein kinase 28 |

**Supplemental table 2.** Identity of the 68 proteins containing both an N-myristoylation motif and a cTP from the predicted *Solanum lycopersicum* proteome.

| ***Solanum lycopersicum* ID** | ***A. thaliana* best match** | **Description** |
|---|---|---|
| Solyc01g091450.3.1 | AT1G01970 | Tetratricopeptide repeat (TPR)-like superfamily protein |
| Solyc10g008490.3.1 | AT1G03590 | Protein phosphatase 2C family protein |
| Solyc10g085410.2.1 | AT1G03910 | . |
| Solyc04g011520.3.1 | AT1G07570 | Protein kinase superfamily protein |
| Solyc05g006170.3.1 | AT1G10510 | RNI-like superfamily protein |
| Solyc08g080360.3.1 | AT1G12710 | Phloem protein 2-A12 |
| Solyc02g093470.3.1 | AT1G13000 | Protein of unknown function (DUF707) |
| Solyc12g009220.2.1 | AT1G19180 | Jasmonate-zim-domain protein 1 |
| Solyc05g012180.3.1 | AT1G26360 | Methyl esterase 13 |
| Solyc05g054880.3.1 | AT1G28060 | Pre-mRNA-splicing factor 3 |
| Solyc03g079860.2.1 | AT1G28240 | Protein of unknown function (DUF616) |
| Solyc08g076350.3.1 | AT1G32190 | alpha/beta-Hydrolases superfamily protein |
| Solyc00g027970.1.1 | AT1G32190 | alpha/beta-Hydrolases superfamily protein |
| Solyc02g092990.1.1 | AT1G47056 | VIER F-box proteine 1 |
| Solyc11g044560.2.1 | AT1G49720 | Abscisic acid responsive element-binding factor 1 |
| Solyc07g053910.3.1 | AT1G53050 | Protein kinase superfamily protein |
| Solyc02g087480.3.1 | AT1G59650 | Protein of unknown function (DUF1336) |
| Solyc01g008870.1.1 | AT1G69790 | Protein kinase superfamily protein |
| Solyc04g082500.3.1 | AT1G76360 | Protein kinase superfamily protein |
| Solyc05g053870.3.1 | AT1G79380 | Ca(2)-dependent phospholipid-binding protein (Copine) family |
| Solyc03g116500.3.1 | AT1G80170 | Pectin lyase-like superfamily protein |
| Solyc11g062400.2.1 | AT2G02800 | Protein kinase 2B |
| Solyc00g007060.3.1 | AT2G17220 | Protein kinase superfamily protein |
| Solyc01g112220.3.1 | AT2G17220 | Protein kinase superfamily protein |
| Solyc03g033540.3.1 | AT2G17890 | Calcium-dependent protein kinase 16 |
| Solyc05g047590.3.1 | AT2G26440 | Plant invertase/pectin methylesterase inhibitor superfamily |
| Solyc06g005500.3.1 | AT2G28930 | Protein kinase 1B |
| Solyc03g115400.2.1 | AT2G35340 | Helicase domain-containing protein |
| Solyc03g005030.3.1 | AT3G01860 | . |
| Solyc05g055790.3.1 | AT3G02750 | Protein phosphatase 2C family protein |
| Solyc09g007700.3.1 | AT3G09770 | RING/U-box superfamily protein |
| Solyc01g028900.3.1 | AT3G13450 | Transketolase family protein |
| Solyc01g109200.3.1 | AT3G18930 | RING/U-box superfamily protein |
| Solyc10g006390.3.1 | AT3G20550 | SMAD/FHA domain-containing protein |
| Solyc02g069560.3.1 | AT3G25690 | Hydroxyproline-rich glycoprotein family protein |
| Solyc02g067040.3.1 | AT3G29180 | Protein of unknown function (DUF1336) |
| Solyc02g089060.3.1 | AT3G29770 | Methyl esterase 11 |
| Solyc09g005850.3.1 | AT3G55140 | Pectin lyase-like superfamily protein |
| Solyc10g079560.1.1 | AT3G57070 | Glutaredoxin family protein |

| | | |
|---|---|---|
| Solyc12g056410.2.1 | AT3G61060 | Phloem protein 2-A13 |
| Solyc07g054300.3.1 | AT4G03415 | Protein phosphatase 2C family protein |
| Solyc12g099530.2.1 | AT4G15880 | Cysteine proteinases superfamily protein |
| Solyc08g008170.3.1 | AT4G23650 | Calcium-dependent protein kinase 6 |
| Solyc03g032150.3.1 | AT4G35600 | Protein kinase superfamily protein |
| Solyc10g084770.2.1 | AT5G02290 | Protein kinase superfamily protein |
| Solyc09g010850.3.1 | AT5G02290 | Protein kinase superfamily protein |
| Solyc09g010110.3.1 | AT5G06130 | Chaperone protein dnaJ-related |
| Solyc12g096390.2.1 | AT5G10700 | Peptidyl-tRNA hydrolase II (PTH2) family protein |
| Solyc03g082500.3.1 | AT5G24430 | Calcium-dependent protein kinase (CDPK) family protein |
| Solyc01g098610.3.1 | AT5G27380 | Glutathione synthetase 2 |
| Solyc10g081470.2.1 | AT5G35220 | Peptidase M50 family protein |
| Solyc04g005810.3.1 | AT5G39950 | Thioredoxin 2 |
| Solyc04g011590.3.1 | AT5G40780 | Lysine histidine transporter 1 |
| Solyc02g076710.3.1 | AT5G43060 | Granulin repeat cysteine protease family protein |
| Solyc08g075080.3.1 | AT5G45290 | RING/U-box superfamily protein |
| Solyc04g076860.3.1 | AT5G49665 | Zinc finger (C3HC4-type RING finger) family protein |
| Solyc07g026680.2.1 | AT5G56840 | Myb-like transcription factor family protein |
| Solyc09g007680.3.1 | AT5G60580 | RING/U-box superfamily protein |
| Solyc03g112770.3.1 | AT5G63030 | Thioredoxin superfamily protein |
| Solyc01g108990.3.1 | NA | NA |
| Solyc00g021650.1.1 | NA | NA |
| Solyc03g123550.1.1 | NA | NA |
| Solyc02g066970.1.1 | NA | NA |
| Solyc01g088640.3.1 | NA | NA |
| Solyc04g025500.2.1 | NA | NA |
| Solyc09g011280.1.1 | NA | NA |
| Solyc11g032110.1.1 | NA | NA |
| Solyc11g017478.1.1 | NA | NA |

**Supplemental table 3.** Identity of the 107 proteins containing both an N-myristoylation motif and a cTP from the predicted *Oryza sativa* subsp. *japonica* proteome.

| *Oryza sativa* subsp. *japonica* ID | *A. thaliana* best match | Description |
|---|---|---|
| LOC_Os03g04550.1 | AT1G04030 | . |
| LOC_Os08g44950.1 | AT1G04360 | RING/U-box superfamily protein |
| LOC_Os04g12720.1 | AT1G05675 | UDP-Glycosyltransferase superfamily protein |
| LOC_Os03g16740.1 | AT1G07570 | Protein kinase superfamily protein |
| LOC_Os03g16740.2 | AT1G07570 | Protein kinase superfamily protein |
| LOC_Os05g02020.1 | AT1G07570 | Protein kinase superfamily protein |
| LOC_Os03g16760.1 | AT1G07630 | pol-like 5 |
| LOC_Os05g40790.2 | AT1G07705 | NOT2 / NOT3 / NOT5 family |
| LOC_Os05g40790.1 | AT1G07705 | NOT2 / NOT3 / NOT5 family |
| LOC_Os02g39970.4 | AT1G15740 | Leucine-rich repeat family protein |
| LOC_Os02g39970.2 | AT1G15740 | Leucine-rich repeat family protein |
| LOC_Os02g39970.1 | AT1G15740 | Leucine-rich repeat family protein |
| LOC_Os02g39970.3 | AT1G15740 | Leucine-rich repeat family protein |
| LOC_Os08g39100.1 | AT1G16220 | Protein phosphatase 2C family protein |
| LOC_Os02g50060.1 | AT1G18210 | Calcium-binding EF-hand family protein |
| LOC_Os07g40550.1 | AT1G18670 | Protein kinase superfamily protein |
| LOC_Os07g40550.2 | AT1G18670 | Protein kinase superfamily protein |
| LOC_Os12g38860.1 | AT1G18670 | Protein kinase superfamily protein |
| LOC_Os04g42670.1 | AT1G47056 | VIER F-box proteine 1 |
| LOC_Os01g22590.1 | AT1G48100 | Pectin lyase-like superfamily protein |
| LOC_Os07g07600.1 | AT1G48120 | Hydrolases;protein serine/threonine phosphatases |
| LOC_Os01g27020.1 | AT1G54610 | Protein kinase superfamily protein |
| LOC_Os01g27020.3 | AT1G54610 | Protein kinase superfamily protein |
| LOC_Os01g27020.2 | AT1G54610 | Protein kinase superfamily protein |
| LOC_Os01g27020.4 | AT1G54610 | Protein kinase superfamily protein |
| LOC_Os01g27020.5 | AT1G54610 | Protein kinase superfamily protein |
| LOC_Os07g43950.1 | AT1G55310 | SC35-like splicing factor 33 |
| LOC_Os03g48660.1 | AT1G59650 | Protein of unknown function (DUF1336) |
| LOC_Os07g48730.1 | AT1G69790 | Protein kinase superfamily protein |
| LOC_Os04g24140.1 | AT1G71100 | Ribose 5-phosphate isomerase, type A protein |
| LOC_Os10g26520.1 | AT2G02800 | Protein kinase 2B |
| LOC_Os09g10600.1 | AT2G05990 | NAD(P)-binding Rossmann-fold superfamily protein |
| LOC_Os06g51170.1 | AT2G17220 | Protein kinase superfamily protein |
| LOC_Os10g29440.1 | AT2G39760 | BTB/POZ/MATH-domains containing protein |
| LOC_Os04g55860.1 | AT3G03010 | Peptidyl-tRNA hydrolase II (PTH2) family protein |
| LOC_Os11g45560.1 | AT3G06190 | BTB-POZ and MATH domain 2 |
| LOC_Os03g15770.2 | AT3G09010 | Protein kinase superfamily protein |
| LOC_Os03g15770.1 | AT3G09010 | Protein kinase superfamily protein |
| LOC_Os03g15770.3 | AT3G09010 | Protein kinase superfamily protein |

| | | |
|---|---|---|
| LOC_Os05g51400.1 | AT3G28690 | Protein kinase superfamily protein |
| LOC_Os07g41230.1 | AT3G29770 | Methyl esterase 11 |
| LOC_Os06g42730.1 | AT3G30380 | alpha/beta-Hydrolases superfamily protein |
| LOC_Os06g51280.1 | AT3G48560 | Chlorsulfuron/imidazolinone resistant 1 |
| LOC_Os03g42710.1 | AT3G50390 | Transducin/WD40 repeat-like superfamily protein |
| LOC_Os03g05740.1 | AT3G54840 | Ras-related small GTP-binding family protein |
| LOC_Os07g29640.3 | AT3G54880 | . |
| LOC_Os07g29640.2 | AT3G54880 | . |
| LOC_Os01g36620.1 | AT3G55140 | Pectin lyase-like superfamily protein |
| LOC_Os02g20140.1 | AT4G10010 | Protein kinase superfamily protein |
| LOC_Os06g08600.1 | AT4G13180 | NAD(P)-binding Rossmann-fold superfamily protein |
| LOC_Os01g65680.1 | AT4G15093 | Catalytic LigB subunit of aromatic ring-opening dioxygenase family |
| LOC_Os03g30934.5 | AT4G18230 | . |
| LOC_Os03g30934.2 | AT4G18230 | . |
| LOC_Os03g30934.3 | AT4G18230 | . |
| LOC_Os03g30934.4 | AT4G18230 | . |
| LOC_Os03g30934.1 | AT4G18230 | . |
| LOC_Os01g06500.1 | AT4G19840 | Phloem protein 2-A1 |
| LOC_Os01g06500.2 | AT4G19840 | Phloem protein 2-A1 |
| LOC_Os01g43410.1 | AT4G23650 | Calcium-dependent protein kinase 6 |
| LOC_Os03g10240.2 | AT4G34320 | Protein of unknown function (DUF677) |
| LOC_Os03g10240.1 | AT4G34320 | Protein of unknown function (DUF677) |
| LOC_Os03g04050.1 | AT4G35230 | BR-signaling kinase 1 |
| LOC_Os10g39670.1 | AT4G35230 | BR-signaling kinase 1 |
| LOC_Os02g58520.1 | AT4G35310 | Calmodulin-domain protein kinase 5 |
| LOC_Os07g23244.1 | AT4G39880 | Ribosomal protein L23/L15e family protein |
| LOC_Os01g74200.1 | AT5G02290 | Protein kinase superfamily protein |
| LOC_Os04g32310.1 | AT5G13160 | Protein kinase superfamily protein |
| LOC_Os06g40650.3 | AT5G14420 | RING domain ligase2 |
| LOC_Os06g40650.4 | AT5G14420 | RING domain ligase2 |
| LOC_Os06g40650.1 | AT5G14420 | RING domain ligase2 |
| LOC_Os06g40650.2 | AT5G14420 | RING domain ligase2 |
| LOC_Os06g12190.1 | AT5G39865 | Glutaredoxin family protein |
| LOC_Os02g51370.1 | AT5G39865 | Glutaredoxin family protein |
| LOC_Os08g37104.1 | AT5G42690 | Protein of unknown function, DUF547 |
| LOC_Os05g32350.1 | AT5G45290 | RING/U-box superfamily protein |
| LOC_Os01g04500.1 | NA | NA |
| LOC_Os04g28050.3 | NA | NA |
| LOC_Os01g71040.1 | NA | NA |
| LOC_Os01g07660.1 | NA | NA |
| LOC_Os01g08464.1 | NA | NA |
| LOC_Os01g08630.1 | NA | NA |
| LOC_Os02g33190.1 | NA | NA |
| LOC_Os03g11080.1 | NA | NA |

| | | |
|---|---|---|
| LOC_Os08g07450.1 | NA | NA |
| LOC_Os08g14010.1 | NA | NA |
| LOC_Os04g13874.1 | NA | NA |
| LOC_Os05g48210.1 | NA | NA |
| LOC_Os04g28050.1 | NA | NA |
| LOC_Os04g28050.2 | NA | NA |
| LOC_Os12g30555.1 | NA | NA |
| LOC_Os06g14160.1 | NA | NA |
| LOC_Os06g37200.1 | NA | NA |
| LOC_Os02g19390.1 | NA | NA |
| LOC_Os10g25502.1 | NA | NA |
| LOC_Os03g39670.1 | NA | NA |
| LOC_Os05g30940.1 | NA | NA |
| LOC_Os11g02810.1 | NA | NA |
| LOC_Os10g40820.1 | NA | NA |
| LOC_Os11g01540.1 | NA | NA |
| LOC_Os11g01950.1 | NA | NA |
| LOC_Os09g32000.1 | NA | NA |
| LOC_Os09g04470.1 | NA | NA |
| LOC_Os04g49880.1 | NA | NA |
| LOC_Os09g13050.1 | NA | NA |
| LOC_Os07g49420.1 | NA | NA |
| LOC_Os03g08030.1 | NA | NA |
| LOC_Os10g18310.1 | NA | NA |

**Supplemental table 4.** Identity of the 19 proteins containing both an N-myristoylation motif and a cTP from the predicted *Chlamydomonas reinhardtii* proteome.

| ***Chlamydomonas reinhardtii* ID** | ***A. thaliana* best match** | **Dual targeting signal in *A. thaliana*** | **Description** |
|---|---|---|---|
| tr\|A2I2U7\|A2I2U7_CHLRE | AT1G01580 | no | ferric reduction oxidase 2 |
| tr\|A0A2K3E1G3\|A0A2K3E1G3_CHLRE | AT1G02690 | no | importin alpha isoform 6 |
| tr\|A0A2K3DXF0\|A0A2K3DXF0_CHLRE | AT1G67890 | no | PAS domain-containing protein tyrosine kinase family protein |
| tr\|A0A2K3D1R0\|A0A2K3D1R0_CHLRE | AT1G78290 | no | Protein kinase superfamily protein |
| tr\|A0A2K3CRC2\|A0A2K3CRC2_CHLRE | AT2G03430 | no | Ankyrin repeat family protein |
| tr\|A0A2K3D8J1\|A0A2K3D8J1_CHLRE | AT3G03305 | no | Calcineurin-like metallo-phosphoesterase superfamily protein |
| tr\|A0A2K3E350\|A0A2K3E350_CHLRE | AT3G17820 | no | glutamine synthetase 1.3 |
| tr\|A0A2K3DRM4\|A0A2K3DRM4_CHLRE | AT3G46920 | no | Protein kinase superfamily protein with octicosapeptide/Phox/Bem1p domain |
| tr\|A0A2K3CXD2\|A0A2K3CXD2_CHLRE | AT3G49660 | no | Transducin/WD40 repeat-like superfamily protein |
| tr\|A0A2K3D913\|A0A2K3D913_CHLRE | AT4G23650 | yes | calcium-dependent protein kinase 6 |
| tr\|A0A2K3CRL8\|A0A2K3CRL8_CHLRE | AT4G31170 | no | Protein kinase superfamily protein |
| tr\|A0A2K3CPE6\|A0A2K3CPE6_CHLRE | AT4G33950 | no | Protein kinase superfamily protein |
| tr\|A0A2K3D5I1\|A0A2K3D5I1_CHLRE | AT5G01390 | no | DNAJ heat shock family protein |
| tr\|A0A2K3DCY0\|A0A2K3DCY0_CHLRE | AT5G14420 | yes | RING domain ligase2 |
| tr\|Q6S7R9\|Q6S7R9_CHLRE | AT5G14740 | no | carbonic anhydrase 2 |
| tr\|A0A2K3CQG1\|A0A2K3CQG1_CHLRE | AT5G52820 | no | WD-40 repeat family protein / notchless protein, putative |
| tr\|A0A2K3DUD7\|A0A2K3DUD7_CHLRE | AT5G52820 | no | WD-40 repeat family protein / notchless protein, putative |
| tr\|A0A2K3DCV1\|A0A2K3DCV1_CHLRE | AT5G52820 | no | WD-40 repeat family protein / notchless protein, putative |
| tr\|A0A2K3E7F9\|A0A2K3E7F9_CHLRE | AT5G57610 | no | Protein kinase superfamily protein with octicosapeptide/Phox/Bem1p domain |

**Supplemental table 5.** Identity of the 30 proteins containing both an N-myristoylation motif and a cTP from the predicted *Marchantia polymorpha* proteome.

| ***Marchantia polymorpha* ID** | ***A. thaliana* best match** | **Dual targeting signal in *A. thaliana*** | **Description** |
|---|---|---|---|
| tr\|A0A176VWK2\|A0A176VWK2_MARPO | AT1G04760 | no | Vesicle-associated membrane protein 726 |
| tr\|A0A176WMK8\|A0A176WMK8_MARPO | AT1G14370 | yes | Protein kinase 2A |
| tr\|A0A2R6XPC6\|A0A2R6XPC6_MARPO | AT1G14370 | yes | Protein kinase 2A |
| tr\|A0A176WLT0\|A0A176WLT0_MARPO | AT1G22040 | no | Galactose oxidase/kelch repeat superfamily protein |
| tr\|A0A176VBR5\|A0A176VBR5_MARPO | AT1G27200 | no | Domain of unknown function (DUF23) |
| tr\|A0A176W8L1\|A0A176W8L1_MARPO | AT1G32583 | no | . |
| tr\|A0A2R6XE67\|A0A2R6XE67_MARPO | AT1G32583 | no | . |
| tr\|A0A2R6XRM5\|A0A2R6XRM5_MARPO | AT1G53050 | no | Protein kinase superfamily protein |
| tr\|A0A176VPD0\|A0A176VPD0_MARPO | AT1G53050 | no | Protein kinase superfamily protein |
| tr\|A0A2R6X0Y5\|A0A2R6X0Y5_MARPO | AT1G53400 | no | Ubiquitin domain-containing protein |
| tr\|A0A176VS21\|A0A176VS21_MARPO | AT1G56080 | no | . |
| tr\|A0A2R6WIW5\|A0A2R6WIW5_MARPO | AT1G59650 | yes | Protein of unknown function (DUF1336) |
| tr\|A0A2R6WIX1\|A0A2R6WIX1_MARPO | AT1G59650 | yes | Protein of unknown function (DUF1336) |
| tr\|A0A176VJ71\|A0A176VJ71_MARPO | AT2G03220 | no | Fucosyltransferase 1 |
| tr\|A0A176VE71\|A0A176VE71_MARPO | AT2G16570 | no | GLN phosphoribosyl pyrophosphate amidotransferase 1 |
| tr\|A0A2R6XS49\|A0A2R6XS49_MARPO | AT2G16570 | no | GLN phosphoribosyl pyrophosphate amidotransferase 1 |
| tr\|A0A2R6WF60\|A0A2R6WF60_MARPO | AT2G17890 | yes | Calcium-dependent protein kinase 16 |
| tr\|A0A176VJU5\|A0A176VJU5_MARPO | AT2G34770 | no | Fatty acid hydroxylase 1 |
| tr\|A0A176WHQ9\|A0A176WHQ9_MARPO | AT2G36250 | no | Tubulin/FtsZ family protein |
| tr\|A0A176WLS8\|A0A176WLS8_MARPO | AT3G14900 | no | . |
| tr\|A0A176WPR4\|A0A176WPR4_MARPO | AT3G50530 | yes | CDPK-related kinase |
| tr\|A0A176VW83\|A0A176VW83_MARPO | AT3G54300 | no | Vesicle-associated membrane protein 727 |
| tr\|A0A176VW93\|A0A176VW93_MARPO | AT4G23650 | yes | Calcium-dependent protein kinase 6 |
| tr\|A7VM55\|A7VM55_CLOEH | AT5G02290 | yes | Protein kinase superfamily protein |
| tr\|A0A2R6W2Z6\|A0A2R6W2Z6_MARPO | AT5G14420 | yes | RING domain ligase2 |
| tr\|A0A176VHG7\|A0A176VHG7_MARPO | AT5G21950 | no | Alpha/beta-Hydrolases superfamily protein |
| tr\|A0A176VKC1\|A0A176VKC1_MARPO | AT5G24280 | no | Gamma-irradiation and mitomycin c induced 1 |
| tr\|A0A2R6WCA9\|A0A2R6WCA9_MARPO | AT5G24280 | no | Gamma-irradiation and mitomycin c induced 1 |
| tr\|A0A176VQF7\|A0A176VQF7_MARPO | AT5G26667 | no | P-loop containing nucleoside triphosphate hydrolases superfamily protein |
| tr\|A0A176W6B2\|A0A176W6B2_MARPO | AT5G44670 | no | Domain of unknown function (DUF23) |

**Supplemental table 6.** Identity of the 21 proteins containing both an N-myristoylation motif and a cTP from the predicted *Physcomitrella patens* proteome.

| ***Physcomitrella patens* ID** | ***A. thaliana* best match** | **Dual targeting signal in *A. thaliana*** | **Description** |
|---|---|---|---|
| tr\|A0A2K1JHV8\|A0A2K1JHV8_PHYPA | AT1G22410 | no | Class-II DAHP synthetase family protein |
| tr\|A0A2K1L0F6\|A0A2K1L0F6_PHYPA | AT1G24040 | no | Acyl-CoA N-acyltransferases (NAT) superfamily protein |
| tr\|A0A2K1IGW0\|A0A2K1IGW0_PHYPA | AT1G26360 | yes | Methyl esterase 13 |
| tr\|A0A2K1JE29\|A0A2K1JE29_PHYPA | AT1G53050 | no | Protein kinase superfamily protein |
| tr\|A0A2K1K3J2\|A0A2K1K3J2_PHYPA | AT1G60000 | no | RNA-binding (RRM/RBD/RNP motifs) family protein |
| tr\|A9TB79\|A9TB79_PHYPA | AT1G80280 | no | alpha/beta-Hydrolases superfamily protein |
| tr\|A9T548\|A9T548_PHYPA | AT2G03500 | no | Homeodomain-like superfamily protein |
| tr\|A0A2K1JML1\|A0A2K1JML1_PHYPA | AT3G13320 | no | Cation exchanger 2 |
| tr\|A0A2K1LA82\|A0A2K1LA82_PHYPA | AT3G21400 | no | . |
| tr\|A0A2K1JHT0\|A0A2K1JHT0_PHYPA | AT3G26935 | no | DHHC-type zinc finger family protein |
| tr\|A0A2K1JKM2\|A0A2K1JKM2_PHYPA | AT3G50530 | yes | CDPK-related kinase |
| tr\|A0A2K1KKY3\|A0A2K1KKY3_PHYPA | AT3G57400 | no | . |
| tr\|A0A2K1KKF5\|A0A2K1KKF5_PHYPA | AT4G10630 | no | Glutaredoxin family protein |
| tr\|A0A2K1J7T9\|A0A2K1J7T9_PHYPA | AT4G19180 | no | GDA1/CD39 nucleoside phosphatase family protein |
| tr\|A0A2K1IVF7\|A0A2K1IVF7_PHYPA | AT4G22240 | no | Plastid-lipid associated protein PAP / fibrillin family protein |
| tr\|A7X9K6\|A7X9K6_PHYPA | AT4G23650 | yes | Calcium-dependent protein kinase 6 |
| tr\|A0A2K1J6M3\|A0A2K1J6M3_PHYPA | AT4G25500 | no | Arginine/serine-rich splicing factor 35 |
| tr\|A0A2K1JR35\|A0A2K1JR35_PHYPA | AT5G01100 | no | O-fucosyltransferase family protein |
| tr\|A0A2K1L1F6\|A0A2K1L1F6_PHYPA | AT5G14420 | yes | RING domain ligase2 |
| tr\|A0A2K1IEY5\|A0A2K1IEY5_PHYPA | AT5G45290 | yes | RING/U-box superfamily protein |
| tr\|A9S7Y1\|A9S7Y1_PHYPA | AT5G55000 | no | Potassium channel tetramerisation domain-containing protein / pentapeptide repeat-containing protein |

**Supplemental table 7.** Plasmids and constructs used in this work.

| PLASMIDS | | |
|---|---|---|
| **Expression cassette/virus** | **Source** | **Destination vector** |
| TYLCV | Rosas-Diaz et al., 2018 | pGWB501 |
| TYLCV_C4$_{1-8}$ | Rosas-Diaz et al., 2018 | pGWB501 |
| 35S:*Rep-FLAG* | This paper | pGWB511 |
| 35S:*C2-FLAG* | This paper | pGWB511 |
| 35S:*C3-FLAG* | This paper | pGWB511 |
| 35S:*CP-FLAG* | This paper | pGWB511 |
| 35S:*V2-FLAG* | This paper | pGWB511 |
| 35S:*C4* | Rosas-Diaz et al., 2018 | pGWB2 |
| 35S:*C4*$_{G2A}$ | Rosas-Diaz et al., 2018 | pGWB2 |
| 35S:*C4-GFP* | Rosas-Diaz et al., 2018 | pGWB5 |
| 35S:*C4*$_{G2A}$*-GFP* | Rosas-Diaz et al., 2018 | pGWB5 |
| 35S:*C4-RFP* | Rosas-Diaz et al., 2018 | pB7RWG2 |
| 35S:*C4*$_{G2A}$*-RFP* | This paper | pB7RWG2 |
| 35S:*CPK16* | This paper | pGWB2 |
| 35S:*CPK16*$_{G2A}$ | This paper | pGWB2 |
| 35S:*CPK16-GFP* | This paper | pGWB5 |
| 35S:*CPK16*$_{G2A}$*-GFP* | This paper | pGWB5 |
| 35S:*C4(EACMV)* | This paper | pGWB2 |
| 35S:*C4*$_{G2A}$*(EACMV)* | This paper | pGWB2 |
| 35S:C4(EACMV) | This paper | pGWB502 |
| 35S:*C4*$_{G2A}$*(EACMV)* | This paper | pGWB502 |
| 35S:*C4(EACMV)-GFP* | This paper | pGWB505 |
| 35S:*C4*$_{G2A}$*(EACMV)-GFP* | This paper | pGWB505 |
| 35S:*C4(BCTV)* | This paper | pGWB2 |
| 35S:*C4*$_{G2A}$*(BCTV)* | This paper | pGWB2 |
| 35S:*C4(BCTV)* | This paper | pGWB502 |
| 35S:*C4*$_{G2A}$*(BCTV)* | This paper | pGWB502 |
| 35S:*C4(BCTV)-GFP* | This paper | pGWB505 |
| 35S:*C4*$_{G2A}$*(BCTV)-GFP* | This paper | pGWB505 |
| 35S:*CP(CMV)* | This paper | pGWB2 |
| 35S:*CP*$_{G2A}$*(CMV)* | This paper | pGWB2 |
| 35S:*CP(CMV)* | This paper | pGWB502 |
| 35S:*CP*$_{G2A}$*(CMV)* | This paper | pGWB502 |
| 35S:*CP(CMV)-GFP* | This paper | pGWB505 |
| 35S:*CP*$_{G2A}$*(CMV)-GFP* | This paper | pGWB505 |
| 35S:*P3(GFLV)* | This paper | pGWB2 |
| 35S:*P3*$_{G2A}$*(GFLV)* | This paper | pGWB2 |
| 35S:*P3(GFLV)* | This paper | pGWB502 |
| 35S:*P3*$_{G2A}$*(GFLV)* | This paper | pGWB502 |
| 35S:*P3(GFLV)-GFP* | This paper | pGWB505 |

| | | |
|---|---|---|
| 35S:*P3*$_{G2A}$*(GFLV)-GFP* | This paper | pGWB505 |
| 35S:*GALA1*$_{G2A}$ | This paper | pGWB2 |
| 35S:*GALA3*$_{G2A}$ | This paper | pGWB2 |
| 35S:*GALA1-GFP* | This paper | pGWB505 |
| 35S:*GALA1*$_{G2A}$*-GFP* | This paper | pGWB505 |
| 35S:*GALA3-GFP* | This paper | pGWB505 |
| 35S:*GALA3*$_{G2A}$*-GFP* | This paper | pGWB505 |
| 35S:*CAS-GFP* | This paper | pGWB5 |
| 35S:*CAS*$_{\Delta 230}$*-GFP* | This paper | pGWB505 |
| 35S:*CAS-RFP* | This paper | pB7RWG2 |
| 35S:*CAS*$_{\Delta 230}$*-RFP* | This paper | pB7RWG2 |
| 35S:*C4*$_{G2A}$*-YFP-N* | This paper | pGTQL1211YN |
| 35S:*C4*$_{G2A}$*-YFP-C* | This paper | pGTQL1221YC |
| 35S:*CAS-YFP-N* | This paper | pGTQL1211YN |
| 35S:*CAS-YFP-C* | This paper | pGTQL1221YC |
| 35S:*AEQ* | Mehlmer et al., 2012 (NASC code: N799874) | pBIN-CYA(K) |
| 35S:*GUS-3xHA* | This paper | pGWB514 |
| pDONR221-*CAS* | This paper | pDONR221 |
| TRV2:*NbCAS* | This paper | pTRV2 |
| TRV2:*SlCAS* | This paper | pTRV2 |

**Supplemental table 8.** Primers used in this work.

| AMPLIFICATION TARGET | SOURCE | SEQUENCE 5'-3' |
|---|---|---|
| **Oligonucleotides to clone in pENTR™/D-TOPO® (1 F: CACC) and the pDONR™221 entry vector (2 F: GGGGACAAGTTTGTACAAAAAAGCAGGCTTG, R: GGGGACCACTTTGTACAAGAAAGCTGGGTC) entry vector (Thermo Scientific)** | | |
| TOPO-CPK16 (with stop codon) | This paper | F: CACCATGGGTCTCTGTTTCTCCTCCGCCGCC<br>R: TTAGACCTTGCGAGAAATAAGATAACCAGG |
| TOPO-CPK16$_{G2A}$ (with stop codon) | This paper | F: CACCATGGCTCTCTGTTTCTCCTCCGCCGCC<br>R: TTAGACCTTGCGAGAAATAAGATAACCAGG |
| TOPO-*CPK16* (without stop codon) | This paper | F: CACCATGGGTCTCTGTTTCTCCTCCGCCGCC<br>R: GACCTTGCGAGAAATAAGATAACCAGG |
| TOPO-*CPK16$_{G2A}$* (without stop codon) | This paper | F: CACCATGGCTCTCTGTTTCTCCTCCGCCGCC<br>R: GACCTTGCGAGAAATAAGATAACCAGG |
| TOPO-*Rep(TYLCV)* (without stop codon) | This paper | F: CACCATGCCTCGTTTATTTAA<br>R: CGCCTTATTGGTTTC |
| TOPO-*C2(TYLCV)* (without stop codon) | This paper | F: CACCATGCAACCTTCGTC<br>R: AATAGTGTTAAGAAATG |
| TOPO-*C3(TYLCV)* (without stop codon) | This paper | F: CACCATGGATTCACGCACAG<br>R: ATAAAATTTATATTTTATATC |
| TOPO-*CP(TYLCV)* (without stop codon) | This paper | F: CACCATGTCGAAGCGACCAG<br>R: ATTTGATATTGAATC |
| TOPO-*V2(TYLCV)* (without stop codon) | This paper | F: CACCATGTGGGACCCACTTC<br>R: GGGCTTCGATACATTC |
| TOPO-*C4(EACMV)* (with stop codon) | This paper (Template: TOPO-*C4(EACMV)* synthesized; NCBI:txid374778) | F: CACCATGGGGTGCCTCATCTCCATG<br>R: CTAAATGCTGGCCCTCCCCCCT |
| TOPO-*C4$_{G2A}$(EACMV)* (with stop codon) | This paper (Template: TOPO-*C4(EACMV)* synthesized; NCBI:txid374778) | F: CACCATGGCGTGCCTCATCTCCATG<br>R: CTAAATGCTGGCCCTCCCCCCT |
| TOPO-*C4(BCTV)* (with stop codon) | This paper (Template: TOPO-BCTV infectious clone synthesized; NCBI:txid268960) | F: CACCATGGGCAACCTCATCTCCACG<br>R: TTAACGCCTTGGCATATGAGTCG |
| TOPO-*C4$_{G2A}$(BCTV)* (with stop codon) | This paper (Template: TOPO-BCTV infectious clone synthesized; NCBI:txid268960) | F: CACCATGGCCAACCTCATCTCCACG<br>R: TTAACGCCTTGGCATATGAGTCG |
| TOPO-*CP(CMV)* (with stop codon) | This paper (Template: TOPO-*CP(CMV)* synthesized; NCBI:txid12305) | F: CACCATGGGCAAATCTGAATCAACC<br>R: TCAAACTGGGAGCACCCCTGATG |
| TOPO-*CP$_{G2A}$(CMV)* (with stop codon) | This paper (Template: TOPO-*CP(CMV)* synthesized; NCBI:txid12305) | F: CACCATGGCCAAATCTGAATCAACC<br>R: TCAAACTGGGAGCACCCCTGATG |
| TOPO-*P3(GFLV)* (with stop codon) | This paper (Template: TOPO-*P3(GFLV)* synthesized; NCBI:txid12274) | F: CACCATGGGTTCTCGTATTAACG<br>R: CTACCTCAAGTCGCCAAAGCAACC |
| TOPO-*P3$_{G2A}$(GFLV)* (with stop codon) | This paper (Template: TOPO-*P3(GFLV)* synthesized; NCBI:txid12274) | F: CACCATGGCTTCTCGTATTAACG<br>R: CTACCTCAAGTCGCCAAAGCAACC |
| TOPO-*C4(EACMV)* (without stop codon) | This paper (Template: TOPO-*C4(EACMV)* synthesized; NCBI:txid374778) | F: CACCATGGGGTGCCTCATCTCCATG<br>R: AATGCTGGCCCTCCCCCCT |
| TOPO-*C4$_{G2A}$(EACMV)* (without stop codon) | This paper (Template: TOPO-*C4(EACMV)* synthesized; NCBI:txid374778) | F: CACCATGGCGTGCCTCATCTCCATG<br>R: AATGCTGGCCCTCCCCCCT |
| TOPO-*C4(BCTV)* (without stop codon) | This paper (Template: TOPO-BCTV infectious clone synthesized; NCBI:txid268960) | F: CACCATGGGCAACCTCATCTCCACG<br>R: ACGCCTTGGCATATGAGTCG |
| TOPO-*C4$_{G2A}$(BCTV)* (without stop codon) | This paper (Template: TOPO-BCTV infectious clone synthesized; NCBI:txid268960) | F: CACCATGGCCAACCTCATCTCCACG<br>R: ACGCCTTGGCATATGAGTCG |

| | | |
|---|---|---|
| TOPO-*CP(CMV)* (without stop codon) | This paper (Template: TOPO-*CP(CMV)* synthesized; NCBI:txid12305) | F: CACCATGGGCAAATCTGAATCAACC<br>R: AACTGGGAGCACCCCTGATG |
| TOPO-*CP*$_{G2A}$*(CMV)* (without stop codon) | This paper (Template: TOPO-*CP(CMV)* synthesized; NCBI:txid12305) | F: CACCATGGCCAAATCTGAATCAACC<br>R: AACTGGGAGCACCCCTGATG |
| TOPO-*P3(GFLV)* (without stop codon) | This paper (Template: TOPO-*P3(GFLV)* synthesized; NCBI:txid12274) | F: CACCATGGGTTCTCGTATTAACG<br>R: CCTCAAGTCGCCAAAGCAACC |
| TOPO-*P3*$_{G2A}$*(GFLV)* (without stop codon) | This paper (Template: TOPO-*P3(GFLV)* synthesized; NCBI:txid12274) | F: CACCATGGCTTCTCGTATTAACG<br>R: CCTCAAGTCGCCAAAGCAACC |
| TOPO-*GALA1* (with stop codon) | This paper | F: CACCATGGGAAACCAGTTTTCGATCA<br>R: CTACAGGGAATGGAACGTCATGC |
| TOPO-*GALA1*$_{G2A}$ (with stop codon) | This paper | F: CACCATGGCAAACCAGTTTTCGATCA<br>R: CTACAGGGAATGGAACGTCATGC |
| TOPO-*GALA3* (with stop codon) | This paper | F: CACCATGGGAAATGGTTTTTCAGTG<br>R: TCAAATCCGCAGCGTCACGCCGAT |
| TOPO-*GALA3*$_{G2A}$ (with stop codon) | This paper | F: CACCATGGCAAATGGTTTTTCAGTG<br>R: TCAAATCCGCAGCGTCACGCCGAT |
| TOPO-*GALA1* (without stop codon) | This paper | F: CACCATGGGAAACCAGTTTTCGATCA<br>R: CAGGGAATGGAACGTCATGC |
| TOPO-*GALA1*$_{G2A}$ (without stop codon) | This paper | F: CACCATGGCAAACCAGTTTTCGATCA<br>R: CAGGGAATGGAACGTCATGC |
| TOPO-*GALA3* (without stop codon) | This paper | F: CACCATGGGAAATGGTTTTTCAGTG<br>R: AATCCGCAGCGTCACGCCGAT |
| TOPO-*GALA3*$_{G2A}$ (without stop codon) | This paper | F: CACCATGGCAAATGGTTTTTCAGTG<br>R: AATCCGCAGCGTCACGCCGAT |
| pDONR221-*CAS* (without stop codon) (*At5g23060*) | This paper | F: GGGGACAAGTTTGTACAAAAAAGCAGGCTTG ATGGCTATGGCGGAAATGGCAACG<br>R: GGGGACCACTTTGTACAAGAAAGCTGGGTC GTCGGAGCTAGGAAGGAACTTGG |
| TOPO-*CAS*$_{\Delta 230}$ (without stop codon) | This paper (Template pDONR221-*CAS*) | F: CACCATGGCTATGGCGGAAATGGCAACG<br>R: GCAAAGAAGGTCAAGCGTTTGAGCC |
| **Oligonucleotides to clone in pTRV2 entry vector** (3 F: *EcoR*I, R: *Sac*I) | | |
| TRV2:*NbCAS* | This paper | F: ATCGGAATTCATGGACGCTCAACCAGTG<br>R: ATCGGAGCTCCTCTGTTCTAATATCAATC |
| TRV2:*SlCAS* | This paper | F: ATCGGAATTCACTTCATCACCACCTAAAG<br>R: ATCGGAGCTCTGCCTGCTTTACTAAAGG |
| **Oligonucleotides for gene expression** | | |
| *ICS1* (*At1g74710*) | Nomura et al., 2012 | F: GCGTCGTTCGGTTACAGG<br>R: ACAGCGAGGCTGAATCTCAT |
| *PAD4* (*At3g52430*) | Nomura et al., 2012 | F: TGCCATACTCAAACTCTTTCTTCA<br>R: CCAAAGTGCGGTGAAAGC |
| *PR1* (*At2g14610*) | Nomura et al., 2012 | F: TGATCCTCGTGGGAATTATGT<br>R: TGCATGATCACATCATTACTTCAT |
| *WRKY33* (*At2g38470*) | Nomura et al., 2012 | F: GGGAAACCCAAATCCAAGA<br>R: GTTTCCCTTCGTAGGTTGTGA |
| *WRKY75* (*At5g13080*) | Nomura et al., 2012 | F: CGTCAAGAACAACAAGTTCCCTA<br>R: CTTTGCACTTGCTTCTTCACAT |
| *PDF1.2* (*At5g44420*) | This paper | F: CTTATCTTCGCTGATCTTGT<br>R: CGTAACAGATACACTTGTGTGC |

| | | |
|---|---|---|
| *AT2G26560* | This paper | F: GAAGTAGCTGGTTGGGGACT<br>R: TCCAGATTCTCGACGGTAGC |
| *AT2G35980* | This paper | F: AACCTAGCCCTCACTGTTCC<br>R: CCTTGGAACGTTGGTGTGAG |
| *C4* | Rosas-Diaz et al., 2018 | F: TGCTGACCTCCTCTAGCTGA<br>R: ATCCGAACATTCAGGCAGCT |
| *CPK16* (*At2g17890*) | This paper | F: GCAACGACACTTGATGAGGA<br>R: GCTTGAAGAATCTCGGCAAC |
| *GALA1* (*RSp0914*) | This paper | F: CTCTACGCCTGCGACAATTT<br>R: AACCGATGCCATCAACTCTT |
| *GALA3* (*RSp0028*) | This paper | F: GCAAGACGATCAAGACGCTG<br>R: TGGGATTGGCGGAGATTCAG |
| *ACTIN2* (*At3g18780*) | This paper | F: CTAAGCTCTCAAGATCAAAGGCTTA<br>R: ACTAAAACGCAAAACGAAAGCGGTT |
| *Rep* | Rosas-Diaz et al., 2018 | F: TGAGAACGTCGTGTCTTCCG<br>R: TGACGTTGTACCACGCATCA |
| *25S ribosomal DNA interspacer (ITS)* | Mason et al., 2008 | F: ATAACCGCATCAGGTCTCCA<br>R: CCGAAGTTACGGATCCATTT |
| *Nb08020g06001* | This paper | F: AACAAAGTCAAGTTCTACG<br>R: TAGTATTCTTGTGACCCTG |
| *NbCAS* (*NbS00026008g0003.1*) | This paper | F: TGCGAAGAAAGTGGAAGCTG<br>R: GCTTACTCTGCAACCAACCC |
| *SlCAS* (*Solyc03g114450.3*) | This paper | F: CTCCGCTCTCTCTTTCAGCT<br>R: TCGGCAAATCTTCCAAAGGG |
| *NbEF1*α | Nicot et al., 2005 | F: ATTGGAAACGGATATGCTCCA<br>R: TCCTTACCTGAACGCCTGTCA |
| *SlACTIN* | Exposito-Rodriguez et al., 2008 | F: CCTCAGCACATTCCAGCAG<br>R: CCACCAAACTTCTCCATCCC |
| *NbICS1* | Sang et al., 2019 | F: GTGTCGGCTCTGCTGTCTTCT<br>R: CTGCGTATAGCACGCCAATC |
| *NbPR1* | Maimbo et al., 2007 | F: GGTCAACACGGCGAAAACC<br>R: GCCTTAGCAGCCGTCATGA |
| *NbCYP71D20* | Segonzac et al., 2011 | F: AAGGTCCACCGCACCATGTCCTTAGAG<br>R: AAGAATTCCTTGCCCCTTGAGTACTTGC |
| *NbACRE31* | Segonzac et al., 2011 | F: AAGGTCCCGTCTTCGTCGGATCTTCG<br>R: AAGAATTCGGCCATCGTGATCTTGGTC |
